# *Tff2* marks gastric corpus progenitors that give rise to pyloric metaplasia/SPEM following injury

**DOI:** 10.1101/2025.04.09.647847

**Authors:** Ruhong Tu, Hualong Zheng, Biyun Zheng, Qing Zhong, Jin Qian, Feijing Wu, Toshiro Shiokawa, Yosuke Ochiai, Hiroki Kobayashi, Quin T. Waterbury, Leah B. Zamechek, Satoru Takahashi, Seiya Mizuno, Changming Huang, Ping Li, Yoku Hayakawa, Timothy C. Wang

**Affiliations:** Division of Digestive and Liver Diseases, Department of Medicine, Columbia University Medical Center, New York, NY 10032, USA; Department of Gastric Surgery, Fujian Medical University Union Hospital, Fuzhou, 350001, China; Graduate School of Medicine, Department of Gastroenterology, The University of Tokyo, 7-3-1, Hongo, Bunkyo-ku, Tokyo, 113-8655, Japan; Laboratory Animal Resource Center in Transborder Medical Research Center, Institute of Medicine, University of Tsukuba, 305-8577, Japan

**Author notes:** **Correspondence:** Timothy C. Wang, M.D.,. Division of Digestive and Liver Diseases, Department of Medicine, Columbia University, College of Physicians and Surgeons, New York, U.S.A. R.T., H.Z., B.Z. and Q.Z. contributed equally as co-first authors of this article. **Abbreviations:** CNV, copy number variation; DT, diphtheria toxin; GC, gastric cancer; GIF, gastric intrinsic factor; HDT, high-dose tamoxifen; H.p, Helicobacter pylori; IF, immunofluorescence; IM, intestinal metaplasia; ISH, in situ hybridization; MNCs, mucous neck cells; MSCs, metaplastic stem-like cells; MNU, N-Methyl-N-nitrosourea; PAS, periodic acid-Schiff; PCs, proliferating cells; TA, transit-amplifying; SP, spasmolytic polypeptide; SPEM, spasmolytic polypeptide-expressing metaplasia; TFF2, trefoil factor family 2; TA, Transit-Amplifying.

## Abstract

**In Brief:** Tu et al. show that Tff2^+^ corpus isthmus cells are TA progenitors, and they, not chief cells, are the primary source of SPEM following injury. Upon Kras mutation, these progenitors directly progress to dysplasia, bypassing metaplasia, highlighting them as a potential origin of gastric cancer.

**Highlights:**

- Tff2^+^ corpus cells are TA progenitors that give rise to secretory cells.
- Tff2^+^ progenitors, not chief cells, are the primary source of SPEM after injury.
- Kras-mutant Tff2^+^ progenitors progress directly to dysplasia, bypassing metaplasia.
- Multi-omics analysis reveals distinct trajectories for SPEM and gastric cancer.

Graphical abstract

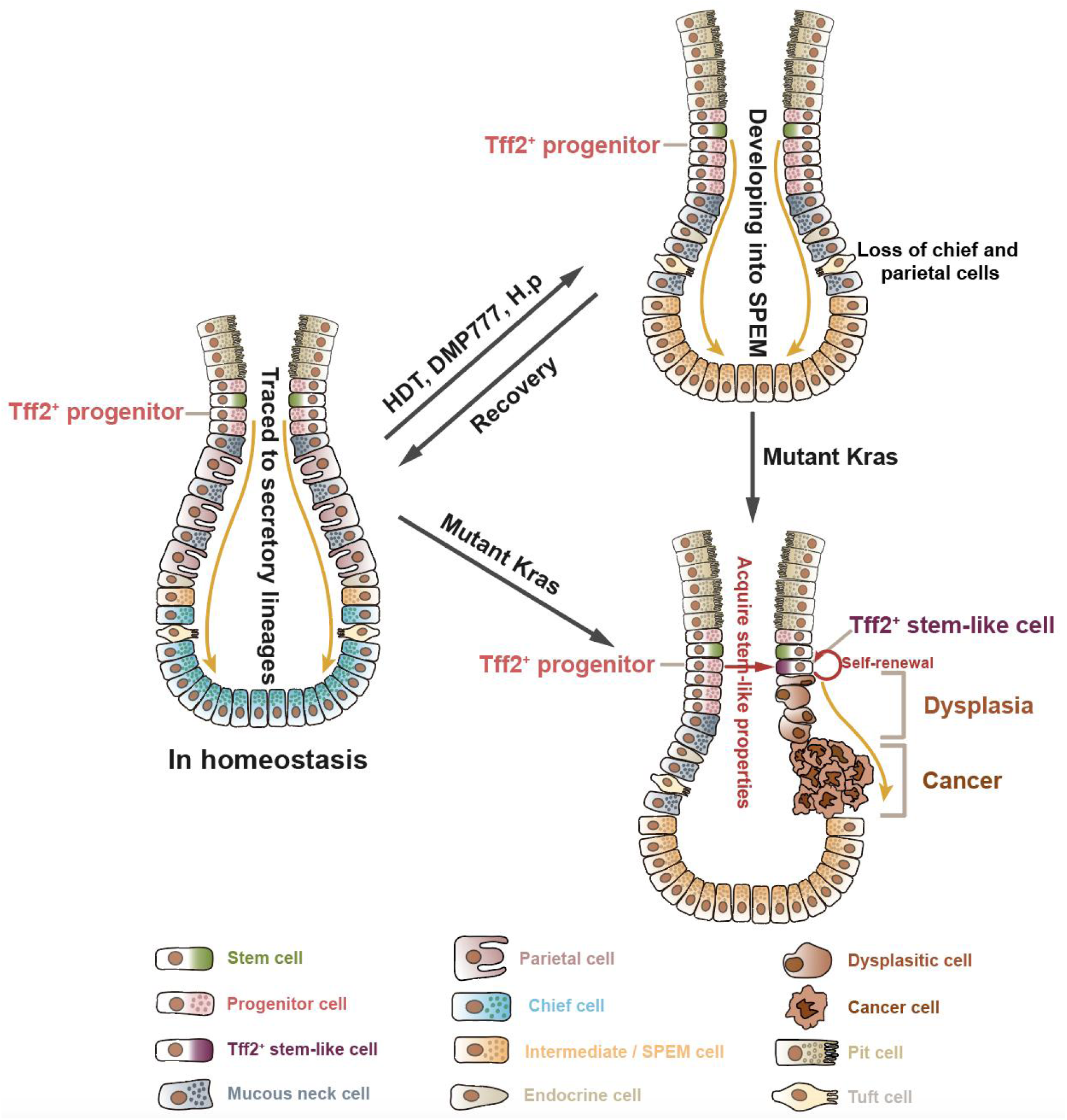

Pyloric metaplasia, also known as spasmolytic polypeptide-expressing metaplasia (SPEM), arises in the corpus in response to oxyntic atrophy, but its origin and role in gastric cancer remain poorly understood. Using *Tff2-CreERT* knockin mice, we identified highly proliferative Tff2^+^ progenitors in the corpus isthmus that give rise to multiple secretory lineages, including chief cells. While lacking long-term self-renewal ability, Tff2^+^ corpus progenitors rapidly expand to form short-term SPEM following acute injury or loss of chief cells. Genetic ablation of Tff2^+^ progenitors abrogated SPEM formation, while genetic ablation of GIF^+^ chief cells enhanced SPEM formation from Tff2^+^ progenitors. In response to *H. pylori* infection, Tff2^+^ progenitors progressed first to metaplasia and then later to dysplasia. Interestingly, induction of Kras^G12D^ mutations in Tff2^+^ progenitors facilitated direct progression to dysplasia in part through the acquisition of stem cell-like properties. In contrast, Kras-mutated SPEM and chief cells were not able to progress to dysplasia. Tff2 mRNA was downregulated in isthmus cells during progression to dysplasia. Single-cell RNA sequencing and spatial transcriptomics of human tissues revealed distinct differentiation trajectories for SPEM and gastric cancer. These findings challenge the conventional interpretation of the stepwise progression through metaplasia and instead identify Tff2^+^ progenitor cells as potential cells of origin for SPEM and possibly for gastric cancer.

## Introduction

Gastric cancer most commonly develops through a stepwise histopathologic pathway from chronic gastritis to atrophy, metaplasia, dysplasia and finally cancer, with atrophy representing the key step in this process.^1,2^ Gastric atrophy represents the loss of glandular secretory cells, a condition that can affect both the antrum and the corpus, usually due to chronic inflammation. In the gastric corpus with atrophic gastritis, both parietal cells and chief cells are lost, and are immediately replaced with epithelial cells resembling more closely the pyloric antrum, termed pyloric metaplasia or SPEM (for spasmolytic polypeptide-expressing metaplasia).^3^ Intestinal metaplasia (IM), where the stomach begins to resemble the intestinal mucosa, usually develops following the emergence of atrophy and SPEM.^1,2,4^ SPEM is strongly associated with the development of dysplasia and cancer, although it remains unresolved whether the lesion is a precursor to dysplasia or a distinct regenerative lesion that progresses no further.^5–11^

SPEM was first described in *Helicobacter felis*-infected mice and is characterized by marked expansion of an aberrant mucous cell lineage expressing a spasmolytic polypeptide (SP), later renamed trefoil factor family 2 (TFF2).^3,12^ Around the same time, the same metaplastic cell lineage was recognized in patients with gastric adenocarcinoma, and shown to be strongly associated with chronic *Helicobacter pylori* (H.p) infection.^9^ Given the expression by the SPEM lineage of TFF2, it was noted early on that SPEM resembled TFF2-expressing mucous neck cells that normally reside in the upper glands, just below the isthmus region. These mucous neck cells, which stain with the GS-II lectin and MUC6, are generated by corpus stem and progenitor cells in the isthmus. The TFF2^+^GSII^+^ neck cells quickly move downward, acquiring expression of gastric intrinsic factor (GIF) and becoming GSII^+^GIF^+^ intermediate cells before losing GSII expression, giving rise to GIF^+^ chief cells at the base.^13–15^ In preclinical models of short-term SPEM, most of the TFF2^+^ metaplastic cells stain positively with both GSII and GIF, thus resembling the normal intermediate cells downstream of mucous neck cells^15^. Nevertheless, long-term SPEM cells show differences from normal neck cells, and can also be identified by expression of APQ5 and CD44v9, more characteristic of deep antral gland cells.^16–18^

The TFF2 peptide is a member of the larger trefoil peptide family, but unlike the other two members (TFF1 and TFF3) contains two trefoil domains and is expressed in the stomach, duodenum and pancreas in mucous secreting cells.^12^ The TFF2 peptide is believed to play roles in gastric cytoprotection and mucosal repair, but is also an important anti-inflammatory peptide.^19,20^ Indeed, studies in TFF2 knockout mice suggest greater responses to *Helicobacter* infection and more rapid gastric carcinogenesis, suggesting that TFF2 may be an atypical tumor suppressor gene.^6^ In the stomach, *Tff2* mRNA is expressed in the lower isthmus region, suggesting expression in an early progenitor cell.^21^ Interestingly, past studies indicated discrepancies between localization of expression of *Tff2* mRNA and TFF2 peptide, with *Tff2* mRNA^+^ cells found above the glandular location of TFF2^+^ mucous neck cells.^21,22^ In addition, lineage tracing studies in the pancreas have indicated that *Tff2* marks a Transit-Amplifying (TA) acinar progenitor that lacks self-renewal but gives rise to exocrine acinar glands in the pancreas.^23^ Nevertheless, the evidence has been limited that Tff2^+^ cells in the stomach are indeed mitotic progenitor cells that give rise to other zymogenic lineages in the gastric corpus. And while the possibility has been acknowledged in the past,^16^ to date no study has directly addressed whether Tff2^+^ cells could be a source of SPEM.

The origins of SPEM have been controversial, with some studies arguing for an origin from chief cells^16,24–26^ while others have advocated for an origin from isthmus progenitor cells.^13,14,27,28^ Since the first report proposing a chief cell origin for SPEM in 2010,^26^ a number of studies have concluded that chief cells may undergo trans-differentiation to generate SPEM.^16,24,25^ However, many of these studies have been based on inducible lineage tracing, with transgenic lines such as Mist1-CreERT,^27^ eR1-CreERT,^28^ and Pgc-CreERT,^29^ which utilize tamoxifen and that express in both isthmus progenitors and chief cells, obscuring the ability to precisely determine the true origin of SPEM. The strongest evidence for a chief cell origin has been the study^24^ that utilized the Gif-rtTA mouse line to provide evidence for a SPEM contribution by traced chief cells. Nevertheless, off-target tracing may be difficult to exclude, and most of the interventions that have been shown to trigger the development of SPEM (high dose tamoxifen, *H. pylori* infection, Kras mutation) have in other studies, been shown to induce chief cell loss rather than transdifferentiation.^13,14,30,31^ Indeed, several of these studies suggest that it is the loss of chief cells that triggers SPEM development.^14,30,31^ Finally, while gastric cancer often develops in the setting of SPEM, the relationship of Tff2^+^ progenitors or SPEM to dysplasia remains uncertain.

Here, we developed a *Tff2-T2A-CreERT2* knock-in mouse line and demonstrated that Tff2-CreERT^+^ specifically marks corpus isthmus progenitors that rapidly expand and give rise to a greater number of SPEM cells following acute or chronic injury. Furthermore, we observed that Tff2^+^ progenitors can acquire stem-like properties and directly progress to dysplasia upon Kras mutation. In contrast, SPEM cells lack the ability to proliferate or undergo passage in vitro, even in the presence of a Kras mutation.

## Results

### Tff2 ^+^ cells mark corpus progenitors and generate multiple secretory lineages

To investigate the origins of SPEM and its relationship to isthmus cells, we generated a *Tff2-CreERT2* knockin mouse line, putting the inducible Cre recombinase ERT2 cassette under the control of the *Tff2*^21^ genetic regulatory elements (Fig. 1A). We crossed *Tff2-CreERT* mice with *Rosa26-tdTomato* mice to generate *Tff2-CreERT; Rosa26-tdTomato* mice and administered a low dose of tamoxifen (75 mg/kg). Lineage tracing of Tff2-expressing cells was performed at selected time points (Fig. S1A). Twenty-four hours post-induction, tdTomato expression was restricted to the lower isthmus region, near the upper third of the corpus glands. A majority of these cells (63.6%) co-expressed Ki67, indicating active proliferation (Fig. 1B, Fig. S1B). Previous studies have suggested that *Tff2* mRNA is expressed in neck cell progenitors, prior to the onset of TFF2 peptide synthesis.^21,22^ To validate this, we combined immunofluorescence (IF) staining for TFF2, which labels mucous neck cells, with in situ hybridization (ISH) to identify Tff2-mRNA*^+^*cells. We observed *Tff2* mRNA expression in the lower isthmus region, with no overlap between TFF2 antibody-positive mucous neck cells (Fig. 1C). Moreover, tdTomato^+^ cells were restricted to Tff2-mRNA*^+^* progenitors and overlapped minimally with TFF2 antibody-positive mucous neck cells (Fig. 1C, Fig. S1C), suggesting that *Tff2-CreERT* specifically labels Transit-Amplifying (TA) progenitors in the isthmus of the corpus.

**Figure 1.**
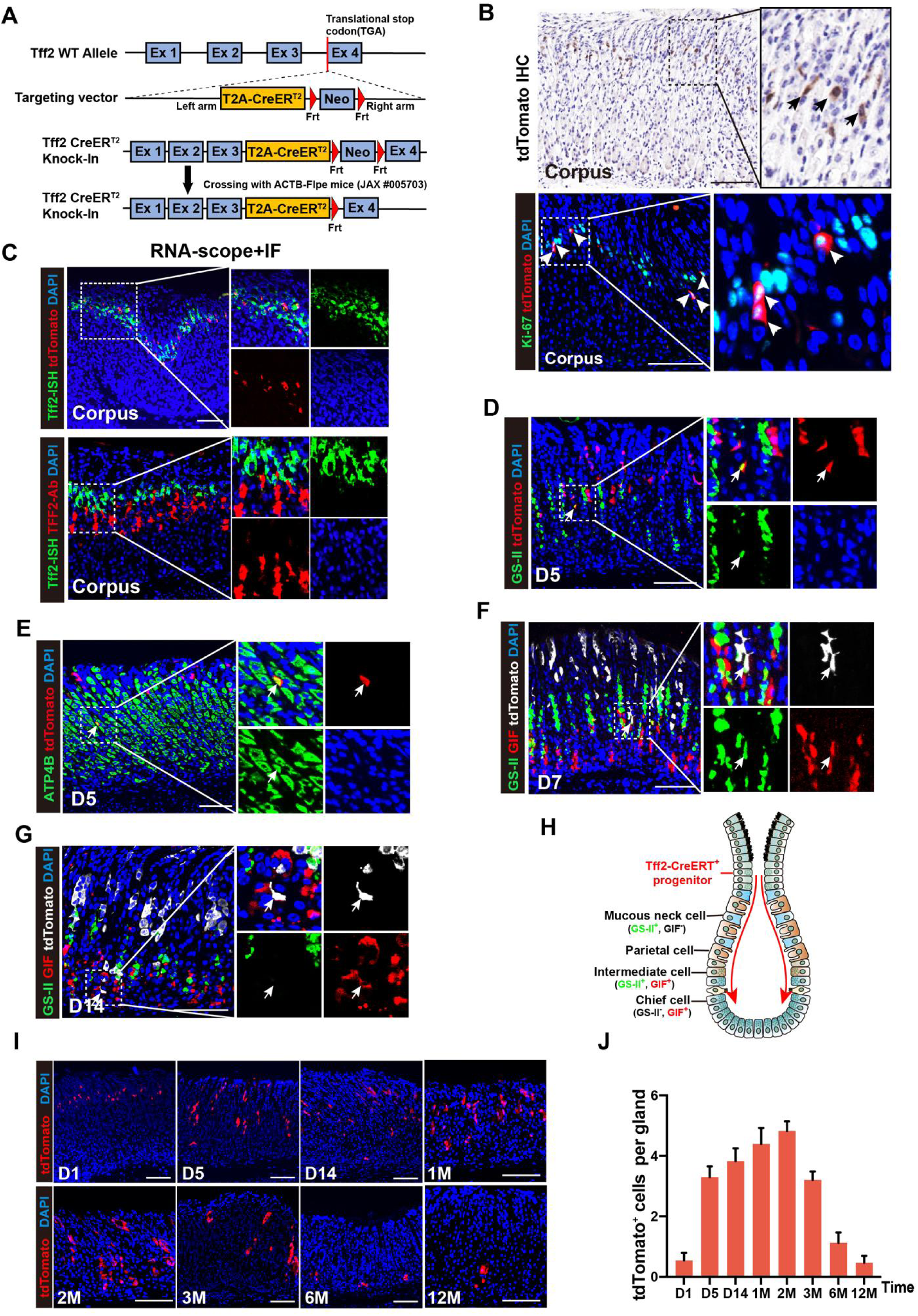
Tff2-CreERT+ cells specifically mark corpus progenitors and generate multiple secretory lineages. (**A**) Schematic of the Tff2 knock-in mouse created for this study. (**B**) (Top) Immunohistochemical staining for tdTomato (brown) and (bottom) immunofluorescence staining for Ki-67 (green), tdTomato (red), and DAPI (blue) in the corpus glands of *Tff2-CreERT2; R26-tdTomato* mice at 24 hours after tamoxifen (75 mg/Kg) induction. Black arrow indicates tdTomato^+^ cell in the isthmus region. White arrow indicates Ki-67^+^ tdTomato^+^ cells in the isthmus region. Scale bars, 100 μm. (**C**) (Top) In situ hybridization of Tff2 (green) and immunofluorescence staining for tdTomato (red), and DAPI (blue) in the corpus glands of *Tff2-CreERT2; R26-tdTomato* mice at 24 hours after tamoxifen (75 mg/Kg) induction. Scale bars, 100 μm. (Bottom) In situ hybridization of Tff2 (green) and immunofluorescence staining for TFF2 (red), and DAPI (blue) in the corpus glands of nontreated wile type (WT) mice. Scale bars, 100 μm. (**D**) Immunofluorescence staining for GS-II (green), tdTomato (red), and DAPI (blue) in the corpus glands of *Tff2-CreERT2; R26-tdTomato* mice at 5 days (D5) after tamoxifen (75 mg/Kg) induction. White arrow indicates GS-II^+^ tdTomato^+^ mucous neck cells. GS-II, lectin GS-II from Griffonia simplicifolia. Scale bars, 100 μm. (**E**) Immunofluorescence staining for ATP4B (green), tdTomato (red), and DAPI (blue) in the corpus glands of *Tff2-CreERT2; R26-tdTomato* mice at 5 days (D5) after tamoxifen (75 mg/Kg) induction. White arrow indicates ATP4B^+^ tdTomato^+^ parietal cells. Scale bars, 100 μm. (**F** and **G**) Immunofluorescence staining for GS-II (green), GIF (red), tdTomato (white), and DAPI (blue) in the corpus glands of *Tff2-CreERT2; R26-tdTomato* mice at 7 days (D7) (F) and 14 days (D14) (G) after tamoxifen (75 mg/Kg) induction. Scale bars, 100 μm. (F) White arrow indicates GS-II^+^ GIF^+^ tdTomato^+^ intermediate cells. (G) White arrow indicates GS-II^-^ GIF^+^ tdTomato^+^ mature chief cells. (**H**) Schematic representation of the lineage tracing of Tff2^+^ progenitors. In homeostasis, after tamoxifen (75 mg/kg) induction, Tff2^+^ corpus isthmus progenitors migrate downward to form first mucous neck cells (GS-II^+^, GIF^-^) and parietal cells (ATP4B^+^), then intermediate cells (GIF^+^, GS-II^+^), and finally mature chief cells (GIF^+^, GS-II^-^). This indicates that Tff2-derived clones are the source of several secretory lineages, including chief cells. (**I** and **J**) Long-term lineage tracing of Tff2^+^ progenitors using the *Tff2-CreERT2; R26-tdTomato* mice, by following their progeny after tamoxifen (75 mg/Kg) induction at different time points (D1, D5, D14, 1M, 2M, 3M, 6M and 12M). (I) Immunofluorescence staining for tdTomato (red) and DAPI (blue) in the corpus glands. Scale bars, 100 μm. (J) Quantification of the number of tdTomato^+^ cells per gland over time as in (I). Data are shown as mean ± SD; n = 3 mice per time point per group.

Twenty-four hours after tamoxifen induction, tdTomato^+^ cells were also detected in the isthmus of the antral glands, just above the base (Fig. S1D), with some co-expressing Ki67 and others co-expressing TFF2 (Fig. S1C, S1D). Notably, the expression of *Tff2* mRNA and TFF2 protein was less stratified in the shorter antral glands compared to the corpus (Fig. S1E), indicating reduced specificity of *Tff2-CreERT* in labeling distinct antral cell populations. In addition, tdTomato^+^ cells were identified in other tissues, including the pancreas,^23^ lung, trachea, and Brunner’s glands of the duodenum (Fig. S1F).

In short-term lineage tracing of Tff2^+^ cells in the corpus, by day 5 post-induction, Tff2^+^ progenitors had migrated downward and differentiated into lectin GS-II (GS-II)/TFF2 positive mucous neck cells (Fig. 1D, Fig. S1G) and ATP4B^+^ parietal cells in the neck region (Fig. 1E).

By day 7, these progenitors gave rise to GIF and GS-II double-positive intermediate cells^13–15^ in the lower third of the gland (Fig. 1F), distinct from metaplastic SPEM cells, as they were CD44v9^−^. By day 14, Tff2^+^ progenitors had differentiated into mature GIF^+^ GS-II^−^ chief cells at the gland base (Fig. 1G), indicating that Tff2^+^ progenitors give rise to multiple secretory lineages, including chief cells (Fig. 1H). In long-term lineage tracing, tdTomato^+^ cells expanded over time, peaking at 2 months post-induction and then gradually declining, with only a few traced chief cells persisting at 6 months (Fig. 1I, 1J). At 12 months, tdTomato^+^ cells were rare, with those remaining being isolated GIF^+^ chief cells located at the gland bases (Fig. S1H). This indicates that Tff2^+^ isthmus cells are TA progenitors that are short-lived and non-self-renewing under homeostatic conditions.

### *Tff2* ^+^ progenitors expand and develop into SPEM after acute injury

We investigated the role of Tff2^+^ progenitors in the regeneration of the gastric corpus following acute injury, characterized by the loss of parietal and chief cells, suggestive of gastric atrophy. *Tff2-CreERT; Rosa26-tdTomato* mice were subjected to high-dose tamoxifen (HDT) treatment (two doses of 6 mg tamoxifen), and corpus tissues were analyzed at selected time points (Fig. S1I). One day after HDT induction, there was a marked depletion of parietal and chief cells (Fig. 2A, 2B). SPEM cells that were GS-II^+^GIF^+^CD44v9^+^ emerged as part of the glandular repair response, peaking between days 3 and 7 (Fig. S2A-I). By two weeks post-treatment, cell morphology, numbers, and marker gene expression had returned to baseline (Fig. S2A-K). Compared to controls, HDT resulted in a more rapid lineage tracing from Tff2^+^ progenitors (Fig. 2C, 2D; Fig. S2L, S2M), which migrated downward faster (Fig. 2C; Fig. S2N), presumably to compensate for the loss of parietal and chief cells. A similar regenerative process was triggered by the acute injury-inducing agent DMP777, which also caused the depletion of parietal and chief cells and led to the expansion of Tff2^+^ progenitors (Fig. 2E; Fig. S3A-E). Downward migration of proliferating isthmus cells was confirmed via EdU labeling (Fig. 2F; Fig. S4A-C), indicating more rapid migration following HDT induction.

**Figure 2.**
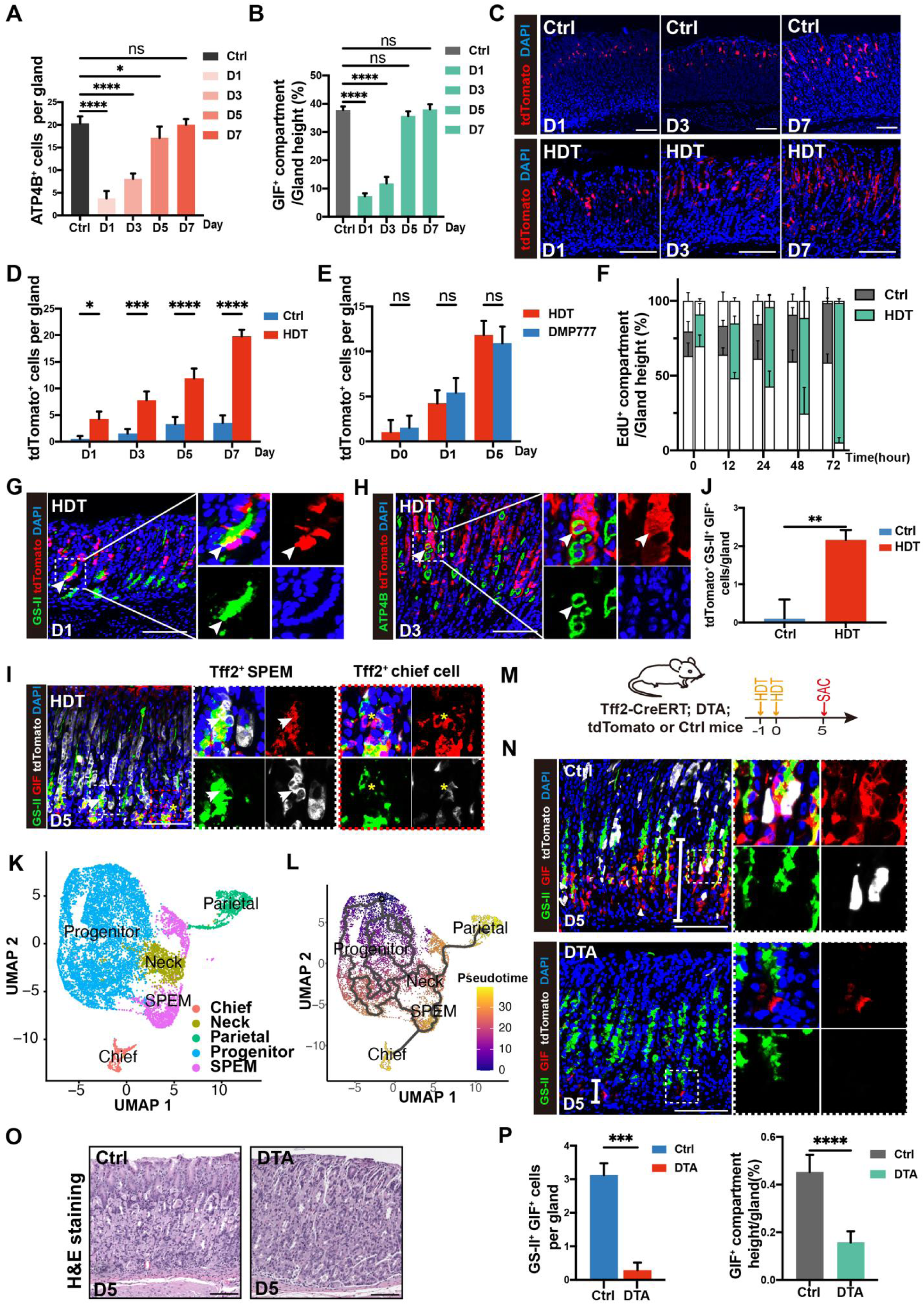
*Tff2* ^+^ progenitors expand and develop into SPEM after acute injury. (**A** and **B**) Quantification of the number of ATP4B^+^ parietal cells per gland (A) and the relative size of the GIF^+^ compartment to the gland height (B) from the corpus of *Tff2-CreERT2; R26-tdTomato* mice before (Ctrl) and after high-dose tamoxifen (HDT)-induced acute injury (D1, D3, D5, and D7). HDT, 300 mg/kg, once daily for 2 consecutive days. Data are shown as mean ± SD; n = 3 mice per time point per group. Statistical significance was assessed with one-way ANOVA. ns, not significant, *P < 0.05, ****P < 0.0001. (**C**) Immunofluorescence staining for tdTomato (red) and DAPI (blue) in the corpus glands of *Tff2-CreERT2; R26-tdTomato* mice after HDT or low-dose tamoxifen (Ctrl, 75 mg/kg, one dose) treatment at different time points (D1, D3, and D7). Scale bars, 100 μm. (**D**) Quantification of the number of tdTomato^+^ cells per gland with the same setting as (C) at different time points (D1, D3, D5 and D7). Data are shown as mean ± SD; n = 3 mice per time point per group. Statistical significance was assessed with two-way ANOVA. ns, not significant. *P < 0.05, ***P < 0.001, ****P < 0.0001. (**E**) Quantification of the number of tdTomato^+^ cells per gland from the corpus of *Tff2-CreERT2; R26-tdTomato* mice before (D0) and after HDT-or DMP777-(500 mg/kg, one dose) treatment (D1 and D5). All groups were treated with tamoxifen (75 mg/kg) to induce nuclear transfer of Cre recombinase 1 day before acute injury induction. Data are shown as mean ± SD; n = 3 mice per time point per group. Statistical significance was assessed with two-way ANOVA. ns, not significant. (**F**) Location and relative size of the EdU^+^ compartment to the gland height in the corpus gland of WT mice after HDT or low-dose tamoxifen (Ctrl, 75 mg/kg) treatment at different time points (H0, H12, H24, H48, and H72) as in (Fig. S4A-C). Data are shown as mean ± SD; n = 3 mice per time point per group. (**G**) Immunofluorescence staining for GS-II (green), tdTomato (red), and DAPI (blue) in the corpus glands of *Tff2-CreERT2; R26-tdTomato* mice at 1 day (D1) after HDT treatment. White arrow indicates GS-II^+^ tdTomato^+^ mucous neck cells. Scale bars, 100 μm. (**H**) Immunofluorescence staining for ATP4B (green), tdTomato (red), and DAPI (blue) in the corpus glands of *Tff2-CreERT2; R26-tdTomato* mice at 3 days (D3) after HDT treatment. White arrow indicates ATP4B^+^ tdTomato^+^ parietal cells. Scale bars, 100 μm. (**I**) Immunofluorescence staining for GS-II (green), GIF (red), tdTomato (white), and DAPI (blue) in the corpus glands of *Tff2-CreERT2; R26-tdTomato* mice at 5 days (D5) after HDT treatment. White arrow indicates GS-II^+^ GIF^+^ tdTomato^+^ SPEM cells. Yellow asterisk indicates GS-II^-^ GIF^+^ tdTomato^+^ mature chief cells. Scale bars, 100 μm. (**J**) Quantification of the number of GS-II^+^ GIF^+^ tdTomato^+^ cells per gland as in (I). Data are shown as mean ± SD; n = 3 mice per group. Statistical significance was assessed by unpaired Student’s t test. **P < 0.01. (**K**) scRNA-seq of Tff2^+^ corpus epithelial cells (tdTomato^+^EpCAM^+^) from HDT (n=6) and untreated control (n=6) of *Tff2-CreERT2; R26-tdTomato* mice. Cell clusters identified based on cell type-specific markers. (**L**) Monocle3 trajectory analysis shows progenitors serve as the origin of all other cell types, including chief cells and SPEMs. (**M**) Schematic representing the experimental design for (L-N). In brief, *Tff2-CreERT; R26-DTA; R26-tdTomato* (DTA) mice were treated with HDT that allowed for simultaneous tracing and ablation of Tff2^+^ progenitors after tamoxifen induction, and corpus tissues were sampled for histological analysis at 5 days (D5) after induction. *Tff2-CreERT; R26-tdTomato* mice were used as controls (Ctrl). (**N**) Immunofluorescence staining for GS-II (green), GIF (red), tdTomato (white), and DAPI (blue) in the corpus glands of DTA mice and matched Ctrl mice. The white line segment represents the height of the GIF^+^ compartment. Scale bars, 100 μm. (**O**) Hematoxylin and eosin (H&E) staining in the corpus glands of DTA mice and Ctrl mice. Scale bars, 100 μm. (**P**) Quantification of the number of GS-II^+^ GIF^+^ SPEM cells per gland (left) and the relative size of the GIF^+^ compartment to the gland height (right) as in (O). Data are shown as mean ± SD; n = 3 mice per group. Statistical significance was assessed by unpaired Student’s t test. ***P < 0.001, ****P < 0.0001.

After HDT, Tff2^+^ progenitors gave rise to differentiated cell types more quickly, producing mucous neck cells by day 1 (Fig. 2G) and parietal cells by day 3 (Fig. 2H), compared to day 5 under homeostatic conditions (Fig. 1D, 1E). These Tff2-derived cells also gave rise to GIF and GSII double-positive cells, some of which we identified as short-term SPEM cells based on CD44v9 staining (Fig. 2I; Fig. S2B, S2E). By day 5, some Tff2^+^ cells had already differentiated into mature chief cells (GIF^+^GSII^−^) (Fig. 2I), a process that typically occurs at day 14 under homeostatic conditions (Fig. 1G). Overall, HDT treatment resulted in a significant increase in the number of Tff2^+^GIF^+^GSII^+^ cells compared to controls (Fig. 2J).

To further validate the progeny of Tff2^+^ progenitors under homeostatic conditions or after acute injury, we specifically sorted Tff2^+^ progenitors and their progeny by flow cytometry and analyzed them by single-cell RNA sequencing (scRNA-seq). We sorted Tff2^+^ progenitors at day 1 (n=3) and their progeny at day 28 (n=3) from *Tff2-CreERT; Rosa26-tdTomato* mice treated with low-dose tamoxifen, using EpCAM^+^ and tdTomato^+^ markers as controls (Ctrl). Similarly, we sorted Tff2^+^ progenitors on day 1 (n=3) and their progeny on day 5 (n=3) from mice treated with high-dose tamoxifen (HDT) to model acute injury. After removing low-quality and mixed cell populations (Fig. S5A-J), we obtained 13628 cells (Ctrl, n=5469; HDT, n=8159) and classified them into 5 clusters according to marker genes: progenitor cells (*Mki67*, and *Top2a*); mucous neck cells (*Muc6*); chief cells (*Gif (Cblif)*); parietal cells (*Atp4b*) and SPEMs (*Muc6*; Tff2; and *Gif*) (Fig. 2K). The Monocle3 trajectory analysis predicted that the Tff2 mRNA^+^ progenitors serve as the origin of all other cell types among these 5 clusters, including chief cells and SPEMs (Fig. 2L).

To further investigate the critical role of Tff2^+^ progenitors in SPEM development and glandular repair, we generated *Tff2-CreERT; Rosa26-DTA; Rosa26-tdTomato* (DTA) mice (heterozygote), allowing simultaneous lineage tracing and ablation of Tff2^+^ progenitors following tamoxifen induction. Mice were analyzed 5 days post-HDT (Fig. 2M), and complete ablation of Tff2^+^ cells was confirmed (Fig. 2N). Ablation of Tff2^+^ cells resulted in impaired glandular repair following HDT (Fig. 2N, 2O). Quantification of SPEM cells (GIF^+^GSII^+^) revealed a significant reduction (>90%) in SPEM formation in DTA mice (Fig. 2P). Additionally, the number of newly formed chief cells was significantly reduced in DTA mice (Fig. 2P), confirming that Tff2^+^ progenitors are essential for the regeneration of lost chief cells.

### Genetic ablation of chief cells accelerates SPEM formation from Tff2^+^ progenitors

Given that HDT eliminates most but not all chief cells (Fig. 2B, Fig. S2B), and that chief cells have been proposed as a possible origin of SPEM, we sought to further clarify the role of chief cells in the development of SPEM. To determine whether more complete ablation of chief cells impacts SPEM formation during regeneration, we administered HDT alongside 1 µg diphtheria toxin (DT) to *Lgr5-DTR-eGFP* mice, achieving full ablation of chief cells (Fig. 3A). We then crossed *Tff2-CreERT* mice with *Lgr5-DTR-eGFP* mice and treated them with HDT and two doses of DT (with the second dose on day 4 to prevent late chief cell regeneration) (Fig. 3B). Notably, chief cell ablation did not inhibit SPEM formation or reduce the number of Tff2^+^ SPEM cells. In fact, we observed an increase in the number of Tff2^+^ SPEM cells in the Lgr5-DTR group compared to controls (Fig. 3C, 3D). These findings support previous observations^30–32^ suggesting that the loss of chief cells may enhance SPEM development from Tff2^+^ progenitors.

**Figure 3.**
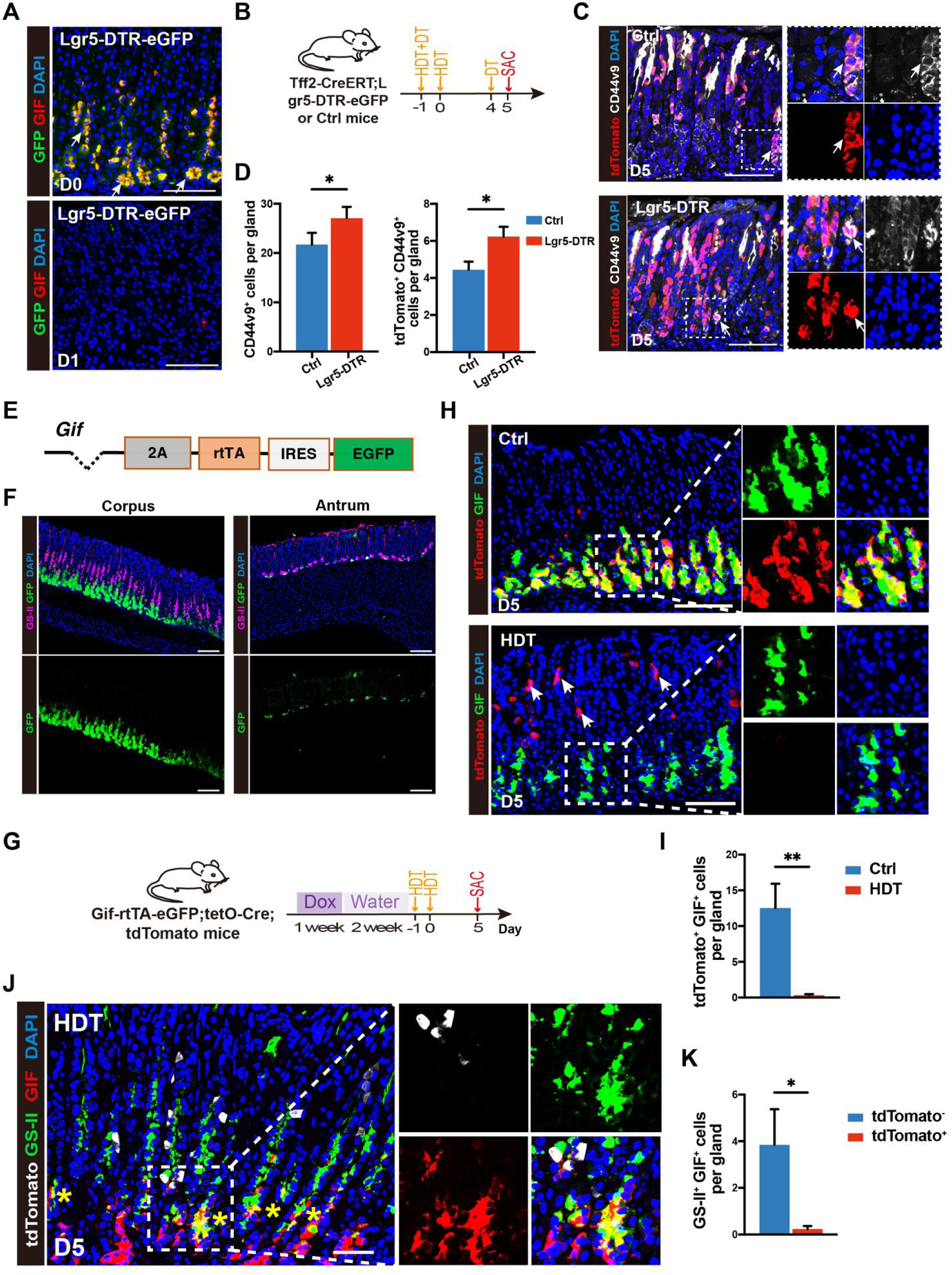
Chief cells are not the origin of SPEM and ablation does not impair SPEM formation. (**A**) Immunofluorescence staining for GFP (green), GIF (red), and DAPI (blue) in the corpus glands of *Lgr5-DTR-eGFP* mice before (D0) and 1 day (D1) after HDT combined with diphtheria toxin (DT, 50 μg/kg) treatment to induce Lgr5^+^ cell depletion. White arrow indicates GFP and GIF co-stained on the chief cells (yellow). Scale bars, 100 μm. (**B**) Schematic representing the experimental design for (C) and (D). In brief, *Tff2-CreERT; Lgr5-DTR-eGFP; R26-tdTomato* (Lgr5-DTR) mice were treated with HDT, as well as two doses of DT (with a second dose of DT was given on day 4 to prevent later regeneration of chief cells), and corpus tissues were sampled for histological analysis at 5 days (D5) after induction. *Tff2-CreERT; R26-tdTomato* mice were used as controls (Ctrl). SAC, sacrify. (**C**) Immunofluorescence staining for tdTomato (red), CD44v9 (white), and DAPI (blue) in the corpus glands of Lgr5-DTR or Ctrl mice 5 days (D5) after HDT as well as DT. White arrow indicates tdTomato^+^ CD44v9^+^ SPEM cells. Scale bars, 100 μm. (**D**) Quantification of the number of CD44v9^+^ cells per gland (left) and the number of tdTomato^+^ CD44v9^+^ SPEM cells per gland (right) as in (C). Data are shown as mean ± SD; n = 3 mice per group. Statistical significance was assessed by unpaired Student’s t test. *P < 0.05. (**E**) Schematic of the Gif knock-in mouse created for this study. (**F**) Immunofluorescence staining for GFP (green), GS-II (purple), and DAPI (blue) in the corpus (left) and antrum (right) glands of *Gif-rtTA-eGFP* mice. Scale bars, 100 μm. (**G**) Schematic representing the experimental design for (H-K). In brief, *Gif-rtTA-eGFP; tetO-Cre; R26-tdTomato* (Gif-rtTA) mice were treated with doxycycline for one week, after two weeks of doxycycline withdrawal, mice were treated with HDT or vehicle (Ctrl), and corpus tissues were sampled for histological analysis at 5 days (D5) after induction. Dox, doxycycline. (**H**) Immunofluorescence staining for GIF (green), tdTomato (red), and DAPI (blue) in the corpus glands of Gif-rtTA mice 5 days (D5) after HDT or Ctrl treatment. White arrow indicates tdTomato^+^ cells. Scale bars, 100 μm. (**I**) Quantification of the number of tdTomato^+^ GIF^+^ cells per gland as in (H). Data are shown as mean ± SD; n = 3 mice per group. Statistical significance was assessed by unpaired Student’s t test. **P < 0.01. (**J**) Immunofluorescence staining for GS-II (green), GIF (red), tdTomato (white) and DAPI (blue) in the corpus glands of Gif-rtTA mice 5 days (D5) after HDT. Yellow asterisk indicates tdTomato^-^ GS-II^+^ GIF^+^ SPEM cells. Scale bars, 100 μm. (**K**) Quantification of the number of GS-II^+^ GIF^+^ SPEM cells per gland derived from tdTomato^+^ or tdTomato^-^ cells as in (J). Data are shown as mean ± SD; n = 3 mice per group. Statistical significance was assessed by unpaired Student’s t test. *P < 0.05.

To further investigate the fate of chief cells during acute injury, we generated a new doxycycline-inducible Tet-ON construct driven by the expression of the chief cell marker *Gif* (*Gif-rtTA-eGFP*) (see Methods) (Fig. 3E). One week after doxycycline administration, Gif^GFP^ cells were localized entirely to the base of the corpus glands and could be clearly distinguished from GS-II^+^ mucous neck cells. A small population of intermediate cells (0.2/gland) co-expressed *Gif^GFP^* and GS-II (Fig. 3F), as previously reported^15^. Additionally, Gif^GFP^ was occasionally detected in some deep antral mucus cells (Fig. 3F). We next crossed *Gif-rtTA* mice with *tetO-Cre* and *Rosa26-tdTomato* mice, allowing us to trace existing Gif^GFP^ cells as tdTomato^+^ post-doxycycline treatment. After doxycycline withdrawal (tet-OFF) for two weeks, newly generated Gif^GFP^ cells were tdTomato^−^, enabling clear lineage tracing (Fig. 3G). Following HDT treatment on day 1, the vast majority of tdTomato^+^ chief cells were lost (Fig. 3H, 3I), and most SPEM cells were derived from tdTomato^−^ cells (Fig. 3J, 3K), confirming that chief cells are not the main source of SPEM. However, with HDT, it was notable that with HDT, tdTomato^+^ cells were observed on day 1 in the isthmus region. These new Gif-tdTomato progenitor cells showed co-localization with Ki67 and expanded downward over the next several days, migrating towards the glandular base (Fig. S6A). In summary, chief cells are lost with HDT but are regenerated from non-chief cell sources. In addition, HDT-associated injury results in upregulation of *Gif* in isthmus progenitors.

### H.p infection or Kras mutation induces Tff2*^+^* progenitors to develop metaplasia and dysplasia

To assess the impact of *H. pylori*-induced gastritis on Tff2^+^ cell lineage tracing, we administered a low dose of tamoxifen (75 mg/kg) to *Tff2-CreERT;Rosa26-tdTomato* mice to ensure synchronized labeling of Tff2^+^ progenitors prior to *H. pylori* infection (Fig. 4A). Two months post-infection (Fig. S6B), we observed a marked reduction in the number of chief cells, accompanied by a significant expansion of neck cells (Fig. 4B). Infected mice also exhibited chronic SPEM, characterized by co-localization of GS-II and GIF staining (Fig. 4B). Lineage tracing further demonstrated that *H. pylori* infection significantly increased both the number of Tff2 progenitor-derived tdTomato^+^ cells and the proportion of proliferating Ki-67^+^ cells compared to uninfected controls (Fig. 4C, 4D). Notably, we identified tdTomato/CD44v9 double positive cells (Fig. S6C), suggesting that *H. pylori* infection promotes the differentiation of Tff2^+^ progenitors into SPEM.

**Figure 4.**
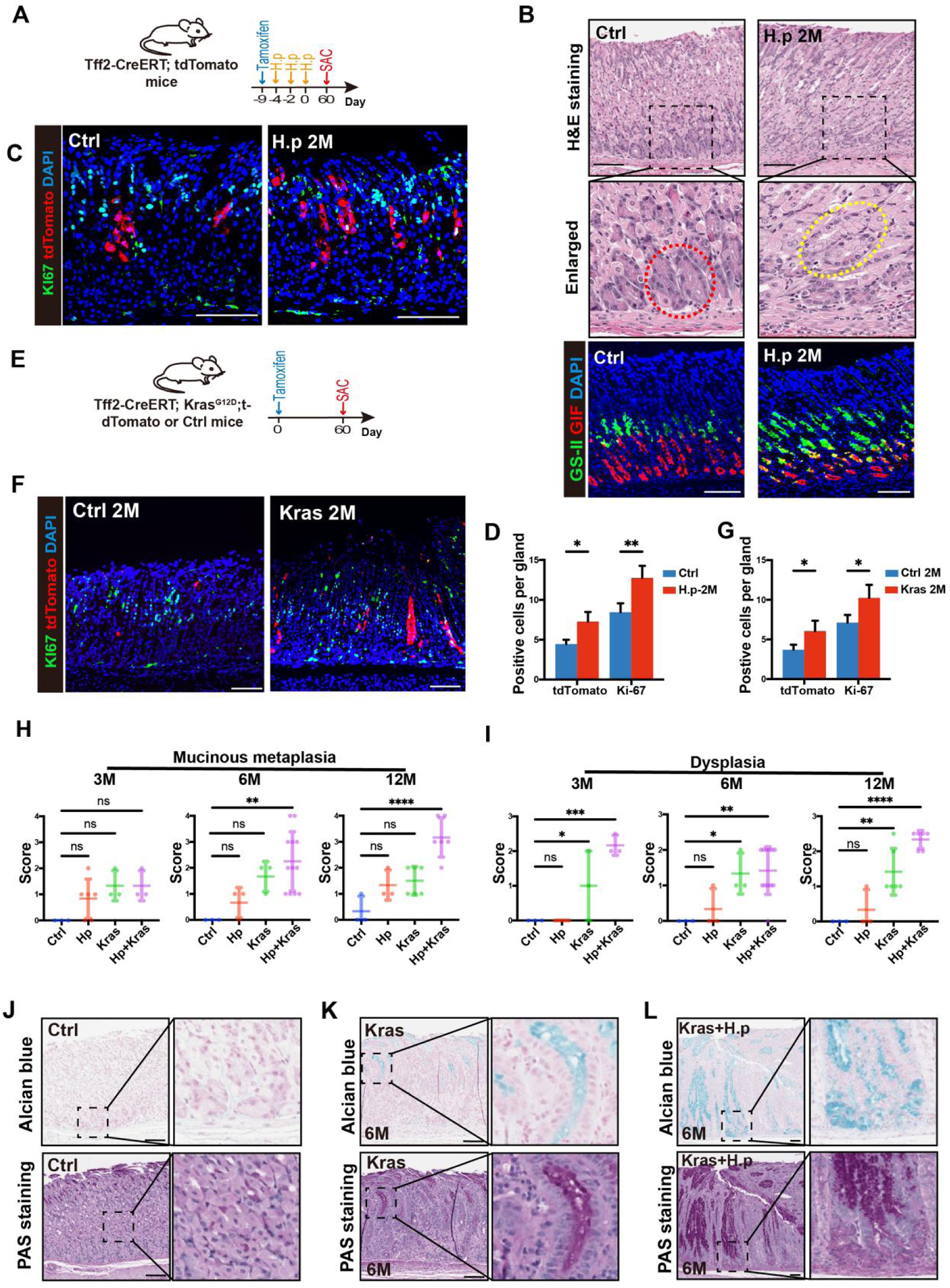
Hp infection or Kras mutation induces Tff2*^+^ progenitors* to develop metaplasia and dysplasia. (**A**) Schematic of the experimental design for the effect of H.p infection on the fate of Tff2^+^ progenitors. In brief, *Tff2-CreERT; R26-tdTomato* mice were treated with tamoxifen (75 mg/kg) to induce nuclear transfer of Cre recombinase, then the mice were orally gavaged with H. p (SS1, 2*109 CFU, every 2 days*3 doses) or vehicle 1 day later. Corpus tissues were then sampled for histological analysis 60 days after oral gavage (2M). SAC, scarify. (**B**) (Top and middle) H&E staining and (bottom) immunofluorescence staining for GS-II (green), GIF (red), and DAPI (blue) in the corpus glands of *Tff2-CreETR; tdTomato* mice 2 months after H.p infection (H.p 2M) or nontreated controls (Ctrl). Yellow dashed box indicates mucinous metaplasia, red dashed box indicates normal chief cell. Scale bars, 100 μm. (**C**) Immunofluorescence staining for Ki-67 (green), tdTomato (red), and DAPI (blue) as in (B). Scale bars, 100 μm. (**D**) Quantification of the number of positive cells per gland (tdTomato^+^ cells and Ki-67^+^ cells) as in (C). Data are shown as mean ± SD; n = 3 mice per group. Statistical significance was assessed by unpaired Student’s t test. *P < 0.05, **P < 0.01. (**E**) Schematic of the experimental design for the effect of Kras mutation on the fate of Tff2^+^ progenitors. In brief, *Tff2-CreERT; Kras^LSL-G12D/+^; R26-tdTomato* (Kras) mice or *Tff2-CreERT; Kras ^+/+^;R26-tdTomato* (Ctrl) mice were treated with tamoxifen (75 mg/kg) to induce Kras mutation and/or nuclear transfer of Cre recombinase. Corpus tissues were then sampled for histological analysis 60 days after induction (2M). (**F**) Immunofluorescence staining for Ki-67 (green), tdTomato (red), and DAPI (blue) in the corpus glands of Kras and Ctrl mice 2 months after induction. Scale bars, 100 μm. (**G**) Quantification of the number of positive cells per gland (tdTomato^+^ cells and Ki-67^+^ cells) as in (F). Data are shown as mean ± SD; n = 3 mice per group. Statistical significance was assessed by unpaired Student’s t test. *P < 0.05. (**H** and **I**) Histopathology score according to the criteria^49^ for mucinous metaplasia (H) and dysplasia (I) in corpus tissues of *Tff2-CreERT; Kras ^+/+^;R26-tdTomato* mice 6 months after H.p infection (H.p) or vehicle (Ctrl), or *Tff2-CreERT; Kras^LSL-G12D/+^; R26-tdTomato* mice 6 months after Kras mutation (Kras) or Kras mutation plus H.p infection (Kras+H.p). Schematic of the experimental design as shown in (Fig. S7G). Data are shown as mean ± SD. n = 3 to 12 mice per group. Statistical significance was assessed with one-way ANOVA. ns, not significant, *P < 0.05, **P < 0.01, ***P < 0.001, ****P < 0.0001. (**J**, **K**, **L**) (Top) Alcian blue staining for acidic mucins (blue) in the corpus glands of *Tff2-CreERT; Kras ^+/+^;R26-tdTomato* mice at 5 days after tamoxifen induction (Ctrl) (J), or *Tff2-CreERT; Kras^LSL-G12D/+^; R26-tdTomato* mice 6 months after Kras (K) and Kras+H.p (L). Cell nuclei were counterstained with hematoxylin (purplish blue). (Bottom) Periodic acid Schiff (PAS) staining neutral mucins (magenta) in the corresponding groups described in (top) panel. Cell nuclei were counterstained with hematoxylin (purplish blue). Scale bars, 100 μm.

To investigate an alternative model of chronic SPEM, we examined the effect of Kras mutation on Tff2^+^ progenitors by using *Tff2-CreERT; Kras^LSL-G12D/+^; Rosa26-tdTomato* (Kras) mice and inducing recombination with low-dose tamoxifen (Fig. 4E). Five days after induction, there was no significant increase in Tff2^+^ cells in the Kras mice compared to controls, indicating that constitutive Kras activation alone does not immediately expand Tff2^+^ progenitors in the absence of gastric injury/cell loss, including chief cells (Fig. S7A, S7B). However, in the context of substantial chief and parietal cell loss following HDT, Kras mutant Tff2^+^ progenitors rapidly generated more Tff2^+^ SPEM cells by day 5, with a significant increase in tdTomato^+^CD44v9^+^ cells compared with Kras^WT^ (Fig. S7C, S7D). At two months post-induction with low-dose tamoxifen, Kras mutation led to a marked increase in both Tff2^+^ cells and proliferating Ki-67^+^ cells (Fig. 4F, 4G). Furthermore, Kras mutation independently promoted the formation of SPEM from Tff2^+^ progenitors (Fig. S7E, S7F).

Given the slow progression of SPEM with Kras mutation alone, and the acceleration with HDT, one possibility is that SPEM development is triggered by the loss of chief cells. Therefore, we next evaluated lesion progression in four experimental groups: *H. pylori* infection (H.p), Kras mutation (Kras), Kras mutation combined with *H. pylori* infection (H.p +Kras), and control (Ctrl) mice. To label a larger population of Tff2^+^ cells prior to the experiment, we administered three low doses of tamoxifen given every other day (75 mg/kg) (Fig. S7G). Mice were analyzed at 3mo, 6mo and 12mo. Both *H. pylori* infection and Kras mutation alone induced some degree of mucinous metaplasia, while the combination H.p.+Kras mice showed the most pronounced metaplastic changes (Fig. 4H). *H. pylori* infection alone resulted in low-grade dysplasia, whereas Kras mutation resulted in moderate-to-severe dysplasia. Importantly, the H.p.+Kras combination accelerated the onset of moderate-to-severe dysplasia, which occurred within three months (Fig. 4I). At the earliest time points (3mo and 6mo), H. pylori infection effectively reduces the number of chief cells (Fig. S7H), whereas Kras mutation of Tff2^+^ progenitors do not, which may explain why the combination of H.p.+Kras accelerates the induction of Kras metaplasia/dysplasia. To confirm the presence of mucinous metaplasia, we performed histologic staining using periodic acid-Schiff (PAS) and Alcian blue.^33^ PAS- and Alcian blue-positive cells were detected in the gastric mucosa of mice from the H.p, Kras, and Hp+Kras groups, but not in controls at 6 months (Fig. 4J-L, Fig. S7I,S7J).

### Dysplasia originates directly from Tff2^+^ cells without transitioning through SPEM

Previous studies have identified Tff2 as a tumor suppressor,^5,6,34^ and its reduced expression has been associated with progression to intestinal-type gastric cancer.^34^ This prompted us to investigate whether dysplastic cells lacking Tff2 expression could originate directly from Tff2^+^ progenitors. To test this hypothesis, we treated *Tff2-CreERT; Kras^LSL-G12D/+^; Rosa26-tdTomato* mice with three low doses of tamoxifen (75 mg/kg) followed by *H. pylori* infection (Fig. S7G). Remarkably, at 6mo, 72.3% of dysplastic lesions (TROP2^+^) could be co-localized with tdTomato^+^ and thus traced back to Tff2^+^ progenitor cells in H.p.+Kras mice (Fig. 5A-C), suggesting that Tff2^+^ progenitors are a potential source of dysplasia.

**Figure 5.**
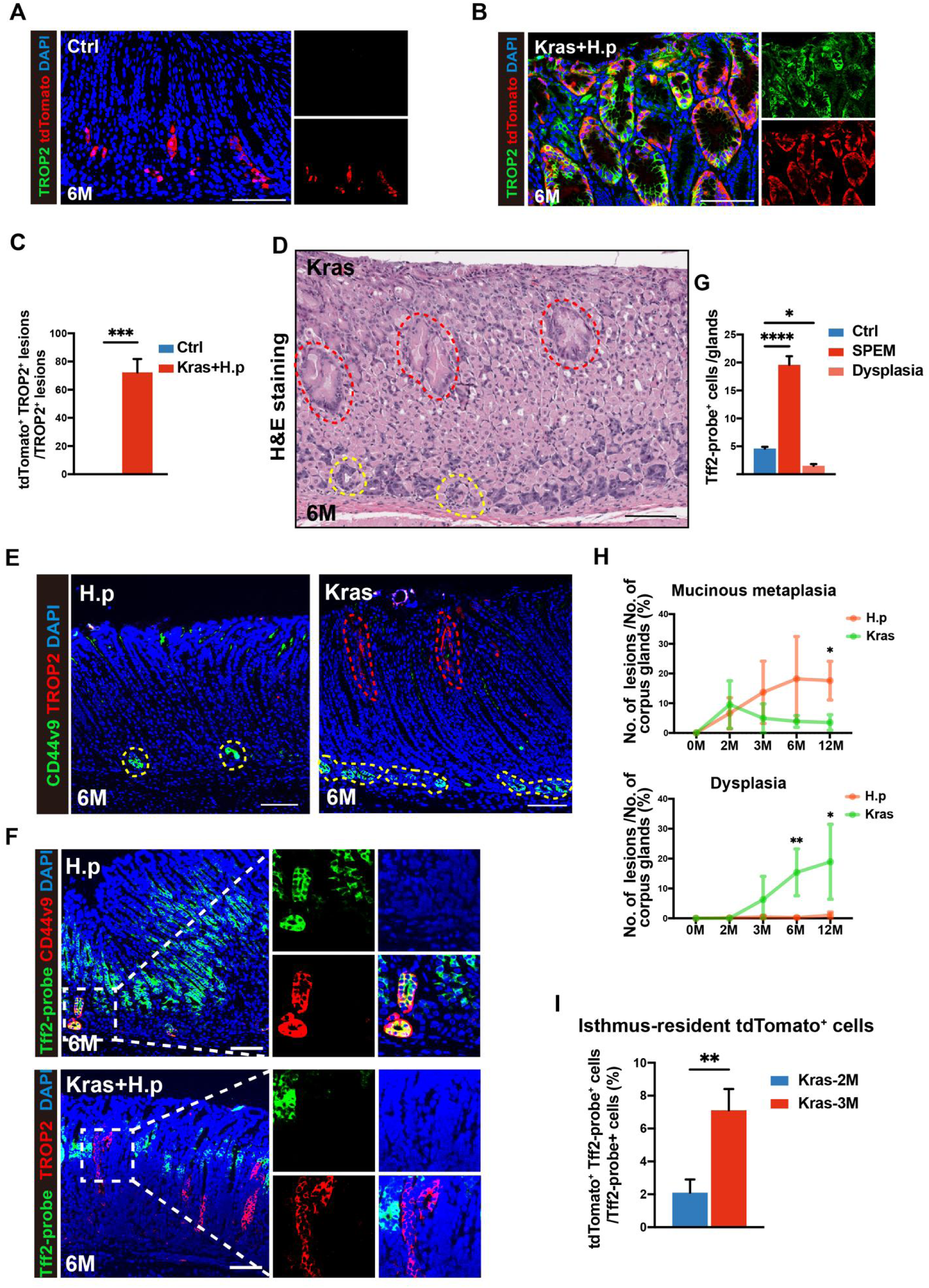
Dysplasia originates directly from Tff2+ cells without transitioning through SPEM. (**A** and **B**) Immunofluorescence staining for TROP2 (green), tdTomato (red), and DAPI (blue) in the corpus glands of *Tff2-CreETR; tdTomato* mice 6 months after tamoxifen induction (Ctrl) (A) or *Tff2-CreETR; Kras^LSL-G12D/+^; tdTomato* mice 6 months after Kras mutation plus H.p infection (Kras+ H.p) (B). Scale bars, 100 μm. (**C**) Quantification of the percentage of tdTomato^+^ TROP2^+^ lesions per TROP2^+^ lesion as in (A and B). Data are shown as mean ± SD; n = 3 mice per group. Statistical significance was assessed by unpaired Student’s t test. *P < 0.05. (**D**) H&E staining in the corpus glands of *Tff2-CreETR; Kras^LSL-G12D/+^; tdTomato* mice 6 months after Kras mutation. Yellow dashed box indicates metaplasia, red dashed box indicates dysplasia. Scale bars, 100 μm. (**E**) Immunofluorescence staining for CD44v9 (green), TROP2 (red), and DAPI (blue) in the corpus glands of *Tff2-CreETR; tdTomato* mice 6 months after H.p infection or *Tff2-CreETR; Kras^LSL-G12D/+^; tdTomato* mice 6 months after Kras mutation. Yellow dashed box indicates metaplasia, red dashed box indicates dysplasia. Scale bars, 100 μm. (**F**) (Top) In situ hybridization of *Tff2* (green) and immunofluorescence staining for CD44v9 (red), and DAPI (blue) in the corpus glands of *Tff2-CreETR; tdTomato* mice 6 months after H.p infection. Scale bars, 100 μm. (Bottom) In situ hybridization of *Tff2* (green) and immunofluorescence staining for TROP2 (red), and DAPI (blue) in the corpus glands of *Tff2-CreETR; Kras^LSL-G12D/+^; tdTomato* mice 6 months after Kras+ H.p. Scale bars, 100 μm. (**G**) Quantification of the number of Tff2-mRMA^+^ cells per gland as in (F). Data are shown as mean ± SD; n = 3 mice per group. Statistical significance was assessed with one-way ANOVA. *P < 0.05, ****P < 0.0001. (**H**) Quantification of the percentage of mucinous metaplasia (top) and dysplasia (bottom) to the total number of corpus glands of *Tff2-CreERT; R26-tdTomato* mice at different time points (0M, 2M, 3M, 6M, 12M) after H.p infection or *Tff2-CreERT; Kras^LSL-G12D/+^; R26-tdTomato* mice after Kras mutation. Schematic of the experimental design as shown in (Fig. S4H). Data are shown as mean ± SD. n = 3 to 12 mice per group. Statistical significance was assessed with multiple unpaired t tests. *P < 0.05, **P < 0.01. (**I**) Quantification of the percentage of tdTomato^+^ Tff2-mRMA^+^ cells to total Tff2-mRMA^+^ cells in the corpus glands of *Tff2-CreETR; Kras^LSL-G12D/+^; tdTomato* mice 2 and 3 months after Kras mutation. Data are shown as mean ± SD; n = 3 mice per group. Statistical significance was assessed by unpaired Student’s t test. **P < 0.01.

Next, we explored whether Tff2^+^ progenitors could give rise to dysplasia independent of SPEM. While H.p.+Kras mice recapitulate clinical features of gastric dysplasia, the frequent co-occurrence of inflammation, metaplasia and dysplasia in this model complicates the interpretation of their sequential relationships (Fig. S8A, S8B). In contrast, in Kras mutant mice with mild chronic inflammation, reduced parietal cells but minimal chief cell loss (Fig. S7H), dysplasia often developed directly in the isthmus region of the glands (Fig. 5D), bypassing SPEM altogether. Importantly, this dysplasia was spatially separated from metaplastic lesions by normal glands (Fig. 5D). Staining for metaplasia (CD44v9) and dysplasia (TROP2) markers confirmed the lack of continuity between these lesions (Fig. 5E), supporting the notion that dysplasia could potentially arise either independently or in conjunction with metaplasia.

We further examined *Tff2* mRNA expression in isthmus, metaplastic, and dysplastic regions using ISH. Six months post *H. pylori* infection, a significant increase in Tff2-mRNA^+^ cells was observed in glands with metaplasia (CD44v9^+^)(Fig. 5F, 5G), particularly in the isthmus region. Conversely, Tff2-mRNA^+^ cells were notably diminished in dysplastic glands (TROP2^+^) in H.p+Kras mice (Fig. 5F, 5G), with a similar reduction in the isthmus. Furthermore, we confirmed that TROP2^+^ dysplastic cells lacking Tff2 mRNA expression were derived from Tff2^+^ progenitors (Fig. S8C). This differential expression pattern suggests that the developmental fate of isthmus cells plays a crucial role in determining cell fate, with downregulation of Tff2 strongly associated with progression to dysplasia.

To investigate the developmental trajectories of SPEM and dysplasia, we quantified the prevalence of metaplastic and dysplastic lesions relative to the total number of corpus glands in Kras and H.p mice (Fig. 5H). Metaplastic lesions peaked within the first two months post-induction, after which their frequency declined, whereas dysplastic lesions progressively increased over time in Kras mice (Fig. 5H). Intriguingly, the peak of metaplasia in Kras mice (e.g. 2 mo) coincided with the peak of Tff2^+^ lineage tracing under homeostatic conditions (Fig. 1I, 1J), consistent with the notion that metaplasia originates from Tff2^+^ progenitors migrating downwards. In contrast, dysplasia in Kras mice may arise from Tff2^+^ progenitors that remain in the isthmus region, possibly acquiring self-renewal and longevity.

We confirmed this hypothesis by analyzing the fate of tdTomato^+^ cells in corpus tissues two and three months after Kras mutation (Fig. 5I, Fig. S8D). At two months, tdTomato^+^ cells had migrated downward into SPEM and other differentiated cell types, leaving only a few tdTomato^+^ cells resident in the isthmus. However, by three months, the number of tdTomato^+^ cells in the isthmus were significantly increased in dysplastic tissues (Fig. 5I, Fig. S8D). These results suggest that while SPEM represents a stable, differentiated state derived from Tff2^+^ progenitors, dysplasia appears to originate from Tff2^+^ cells that remain arrested in a progenitor state, unable to differentiate further and thus bypassing SPEM.

### Tff2^+^ progenitor acquire stem-like properties before progressing into dysplasia

To demonstrate that Tff2^+^ progenitors may give rise to injury-repair lineages and contribute to dysplasia, we analyzed the Tff2^+^ cell lineage scRNA-seq data. CytoTRACE analysis revealed that Tff2^+^ progenitors exhibited the highest differentiation potential, followed by SPEM cells, with chief cells showing the lowest potential (Fig. S5H). Notably, the differentiation capacity of Tff2^+^ progenitors was further enhanced post-injury (Fig. S5I), accompanied by increased expression of proliferation and stem cell-associated genes, including *Cd44*, *Sox2*, and *Yap1* (Fig. S5J). To explore whether the contribution to injury repair lineage and dysplasia is conserved in other isthmus stem/progenitor cell lineages, we re-analyzed scRNA-seq data^35^ from mouse gastric corpus models of acute and chronic injury (Fig. S9A-S9J). The results also revealed that isthmus cells had the highest differentiation potential, which was further enhanced after injury, accompanied by increased expression of proliferation and stem cell-related genes (Figure S9G).

To further validate the in vitro experiments, we isolated Tff2^+^ progenitors from DMP777-induced acute injury and control mice for organoid culture (Fig. 6A). Organoids derived from Tff2^+^ progenitors from DMP777-treated mice displayed a larger growth area (Fig. 6B, Fig. S10A), a higher proportion of Ki-67^+^ cells (Fig. 6C, 6D), and a reduced rate of apoptosis compared to Tff2^+^ progenitors control (untreated) mice (Fig. 6E, 6F). Both organoid types differentiated into GS-II/GIF double-positive cells (SPEM in DMP777-treated mice, intermediate cells in controls) (Fig. S10B), highlighting the pivotal role of isthmus cells in post-injury repair and their enhanced differentiation and proliferative capacity.

**Figure 6.**
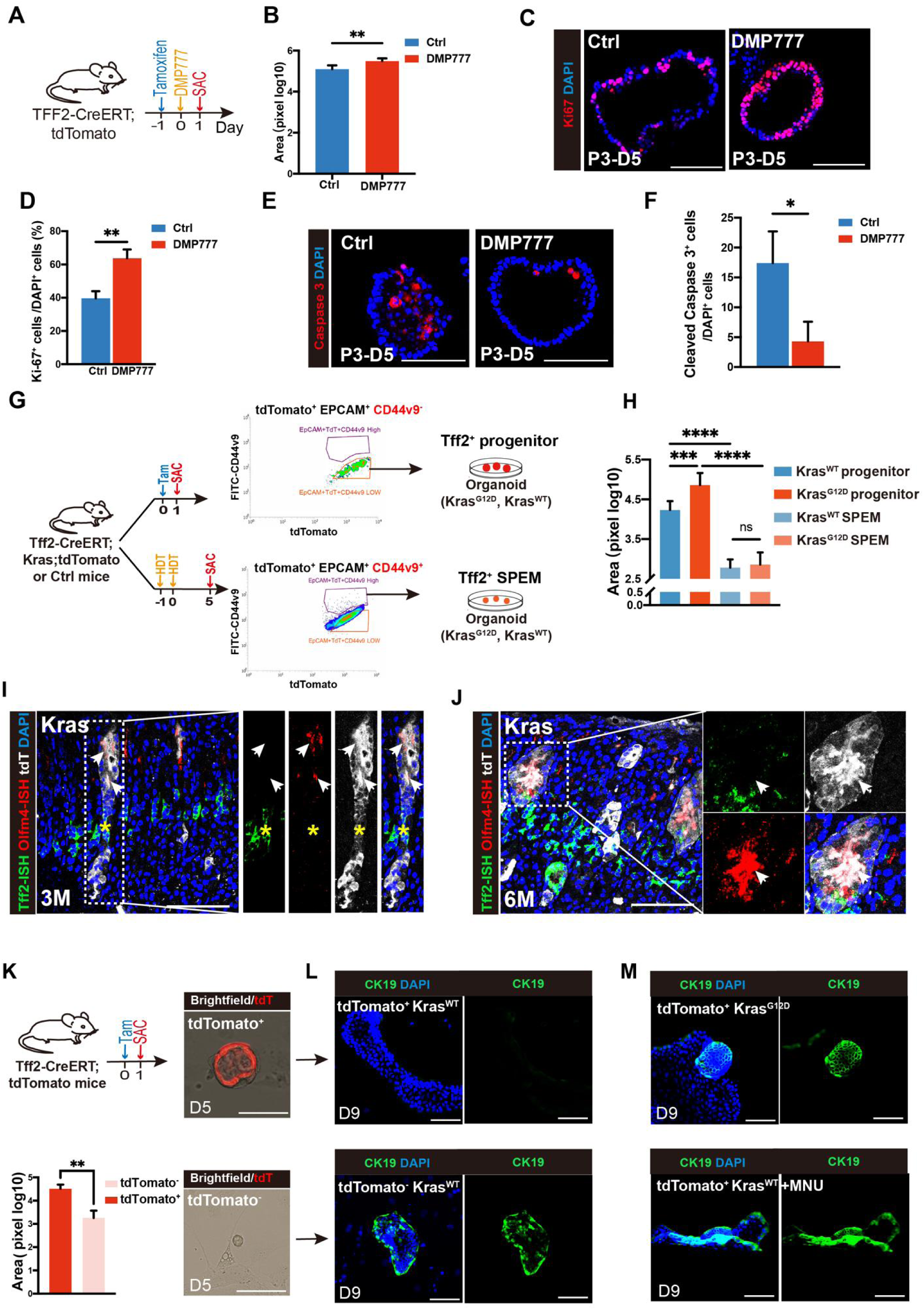
Tff2^+^ progenitor acquire stem-like properties before progressing into dysplasia. **(A)** Schematic representing the experimental design for screening of Tff2^+^ progenitors for organoid culture after DMP777-induced acute injury or nontreated controls (Ctrl). In brief, *Tff2-CreERT; R26-tdTomato* mice were treated with tamoxifen (75 mg/kg) to induce Cre recombinase 1 day before DMP777 treatment or nontreated. Flow cytometric selecting of tdTomato^+^ EpCAM^+^ cells as Tff2^+^ progenitors 1 day after DMP777 treatment or Ctrl and tested them for 3D organoid growth. (**B**) Organoid growth quantification evaluated as grown area (log10 scale) as in (A). Data are shown as mean ± SD. n = 3 mice per group. Statistical significance was assessed by unpaired Student’s t test. **P < 0.01. (**C**) Immunofluorescence staining for Ki67 (red) and DAPI (blue) 5 days after the third passage (P3-D5) of organoids comparing the proliferation capacity of Tff2^+^ progenitors from DMP777 and Ctrl mice as in (A). Scale bars, 100 μm. (**D**) Quantification of the percentage of Ki-67^+^ cells to total DAPI^+^ cells in organoids as in (C). Data are shown as mean ± SD; n = 3 mice per group. Statistical significance was assessed by unpaired Student’s t test. **P < 0.01. (**E**) Immunofluorescence staining for Cleaved Caspase 3 (red) and DAPI (blue) 5 days after the third passage (P3-D5) of organoids comparing the anti-apoptotic capacity of Tff2^+^ progenitors from DMP777 and Ctrl mice as in (A). Scale bars, 100 μm. (**F**) Quantification of the percentage of Cleaved Caspase 3^+^ cells to total DAPI^+^ cells in organoids as in (E). Data are shown as mean ± SD; n = 3 mice per group. Statistical significance was assessed by unpaired Student’s t test. *P < 0.05. (**G**) Schematic representing the experimental design for screening and organoid culture of Tff2^+^ progenitor or SPEM cells. In brief, *Tff2-CreERT; Kras^LSL-G12D/+^; R26-tdTomato mice* (Kras^G12D^) or *Tff2-CreERT; Kras ^+/+^;R26-tdTomato* mice (Kras^WT^) were used. Flow cytometric selecting of CD44v9^-^ tdTomato^+^ EpCAM^+^ cells as Tff2^+^ Kras^G12D^ or Tff2^+^ Kras^WT^ progenitors 1 day after low-dose tamoxifen induction, selecting of CD44v9^+^ tdTomato^+^ EpCAM^+^ cells as Tff2^+^ Kras^G12D^ or Tff2^+^ Kras^WT^ SPEM cells 5 days after HDT treatment, and tested them for 3D organoid growth. (**H**) Organoid growth quantification evaluated as grown area (log10 scale) with the same setting as in (G). Data are shown as mean ± SD. n = 3 mice per group. Statistical significance was assessed with one-way ANOVA. ns, not significant, ***P < 0.001, ****P < 0.0001. (**I** and **J**) In situ hybridization of Tff2 (green), Olfm (red), immunofluorescence staining for tdTomato (white), and DAPI (blue) in the corpus glands of *Tff2-CreETR; Kras^LSL-G12D/+^; tdTomato* (Kras) mice 3 (I) or 6 (J) months after Kras mutation. White arrow indicates Olfm-mRMA^+^ cells derived from tdTomato^+^ progenitors. Yellow asterisk indicates tdTomato^+^ Tff2-mRMA^+^ progenitors. Scale bars, 100 μm. (**K**) (Top, left) Schematic representing the experimental design for screening of tdTomato^+^ progenitors and tdTomato^−^ epithelial cells for 2D cell culture. In brief, *Tff2-CreERT; Kras^LSL-G12D/+^; tdTomato* (Kras^G12D^) and *Tff2-CreERT; Kras ^+/+^; tdTomato* (Kras^WT^) mice were treated with tamoxifen (75 mg/kg) to induce Cre recombinase for 1 day. Flow cytometric selecting of tdTomato^+^ EpCAM^+^ cells as Tff2^+^ progenitors and tdTomato^-^ EpCAM^+^ cells as other epithelial cells 1 day after tamoxifen-induced and tested them for 2D cell growth. (Bottom, left) 2D cell growth quantification evaluated as grown area (log10 scale) with the same setting as in (top, left). Data are shown as mean ± SD. n = 3 mice per group. Statistical significance was assessed by unpaired Student’s t test. **P < 0.01. (Right) Bright field and fluorescence images of epithelial 2D cell cultures at 5 days (D5) for tdTomato^+^ progenitors and tdTomato^−^ epithelial cells as in (top, left). Scale bars, 100 μm. (**L**) Immunofluorescence staining for Cytokeratin 19 (CK19) (green) and DAPI (blue) at 9 days (D9) for tdTomato^+^ Kras^WT^ progenitors and tdTomato^−^ Kras^WT^ epithelial cells as in (K). Scale bars, 100 μm. (**M**) Immunofluorescence staining for CK19 (green) and DAPI (blue) at day 9 (D9) for both tdTomato^+^ Kras^G12D^ progenitors (top) and tdTomato^+^ Kras^WT^ progenitors treated with N-Methyl-N-nitrosourea (MNU, 20 nmol/L) for 24hours starting at day 7 (bottom). Scale bars, 100 μm.

To explore whether Kras mutation alters the differentiation potential of Tff2^+^ progenitors or SPEM cells, we sorted Tff2^+^ progenitors at day 1 from *Tff2-CreERT; Kras^LSL-G12D/+^; Rosa26-tdTomato* (Kras^G12D^) and *Tff2-CreERT; Kras^+/+^;Rosa26-tdTomato* (Kras^WT^) mice treated with low-dose tamoxifen using EpCAM^+^ and tdTomato^+^ markers (Fig. 6G). In order to isolate SPEM cells for comparison, tdTomato^+^ cells were sorted 5 days after HDT induction using EpCAM^+^, tdTomato^+^, and CD44v9^+^ markers from the same mice (Fig. 6G). Organoid cultures revealed that Tff2^+^ Kras^G12D^ progenitors exhibited a significantly larger growth area compared to Kras^WT^, whereas SPEM cells showed reduced growth and an inability to expand in culture, even in the presence of the Kras^G12D^ (Fig. 6H, Fig. S10C, S10D). Additionally, in *Gif-rtTA-eGFP; tetO-Kras^G12D^* mice (Fig. S11A), Kras mutation led to a loss of Gif^+^ chief cells 7 days after doxycycline treatment, accompanied by an increase in Ki-67^+^ proliferating cells and GS-II^+^ mucous neck cells (Fig. S11B-C). Prolonged activation of the tetO-Kras system (90 days) resulted in continued loss of Gif^+^ chief cells without progression to dysplasia (Fig. S11D). These findings indicate that Tff2^+^ progenitors, but not SPEM or chief cells, are the primary candidates for harboring Kras mutations.

To further confirm that Tff2^+^ progenitors acquire stem-like properties following Kras mutation and could serve as precursor cells for intestinal-type gastric cancer, we investigated the expression of the intestinal stem cell marker OLFM4 (Fig. S11E). While OLFM4 is absent in normal gastric stem cells (Fig. S11F), it is expressed in metaplastic stem cells and intestinal-type gastric cancer.^36–38^ ISH combined with IF staining stomach sections from *Tff2-CreERT; Kras^LSL-G12D/+^; Rosa26-tdTomato* mice revealed that Olfm4 mRNA was predominantly present in the isthmus of the tdTomato^+^ gastric corpus glands, above the Tff2-mRNA^+^ cells, as part of the Tff2 lineage, three months post-induction of Kras mutation (Fig. 6I). tdTomato^+^ Olfm4-mRNA^+^ cells exhibited the ability to differentiate throughout the gland (Fig. 6I). As dysplasia developed, *Olfm4* expression was further upregulated in tdTomato^+^ dysplastic lesions (Fig. 6J). Importantly, Olfm4-mRNA^+^ cells remained confined to the region above Tff2-mRNA^+^ cells and did not appear within SPEM lesions. This supports a model in which Tff2^+^ progenitors undergo a direct transformation into stem-like cells and dysplasia, independent of SPEM.

The enhanced stem-like properties of Tff2^+^ progenitors were further evidenced by their increased differentiation potential. While Tff2^+^ progenitors do not normally give rise to pit cells under homeostatic conditions, lineage tracing revealed that following Kras mutation, Tff2^+^ cells were able to differentiate into pit cells, consistent with an increase in their stemness (Fig. S11G). To confirm this, we used a 2D culture system suitable for pit cell growth^39^ and sorted tdTomato^+^ progenitors and tdTomato^−^ epithelial cells (including stem and mature epithelial cells) from *Tff2-CreERT; Kras^LSL-G12D/+^; Rosa26-tdTomato* (Kras^G12D^) and *Tff2-CreERT; Kras^+/+^; Rosa26-tdTomato* (Kras^WT^) mice one day after low-dose tamoxifen induction (Fig. 6K). tdTomato^+^ cells displayed a significantly larger growth area compared to tdTomato^−^ cells (Fig. 6K). IF staining showed that tdTomato^+^ Kras^WT^ cells failed to differentiate into cells expressing CK19, a pit cell marker,^21^ whereas tdTomato^+^ Kras^G12D^ cells and tdTomato^+^ Kras^WT^ cells treated with the carcinogen (N-Methyl-N-nitrosourea, MNU) successfully differentiated into pit cells (Fig. 6L, 6M). In summary, these findings demonstrate that Tff2^+^ progenitors acquire stem cell-like properties and enhanced differentiation potential following Kras mutation or carcinogen exposure, exhibiting a differentiation arrest and driving the development of dysplasia without involvement of SPEM.

### SPEM and gastric cancer arise from the isthmus but follow distinct differentiation paths in humans

To determine whether the differentiation of isthmus progenitor cells into SPEM and gastric cancer (GC) is conserved in humans, we performed scRNA-seq and spatial transcriptomics analyses. First, we examined the expression pattern of *TFF2* mRNA in human gastric tissue and found that its distribution mirrors that seen in mice (Fig. 7A). A representative spatial transcriptomics sample revealed a clear boundary between *TFF2* and *MUC6* mRNA expression in normal gastric corpus mucosa. Specifically, *TFF2* mRNA was primarily localized to the isthmus and pit cell regions, while in metaplastic areas, *TFF2* co-expressed with *MUC6*. ISH further confirmed *TFF2* mRNA expression in the isthmus and in mucinous metaplasia of normal human gastric corpus, with a subsequent decrease in dysplasia (Fig. 7B, Fig.S12A, S12B). These findings suggest that the *TFF2* mRNA expression pattern in isthmus, SPEM, and dysplasia observed in mice are also applicable to humans.

**Figure 7.**
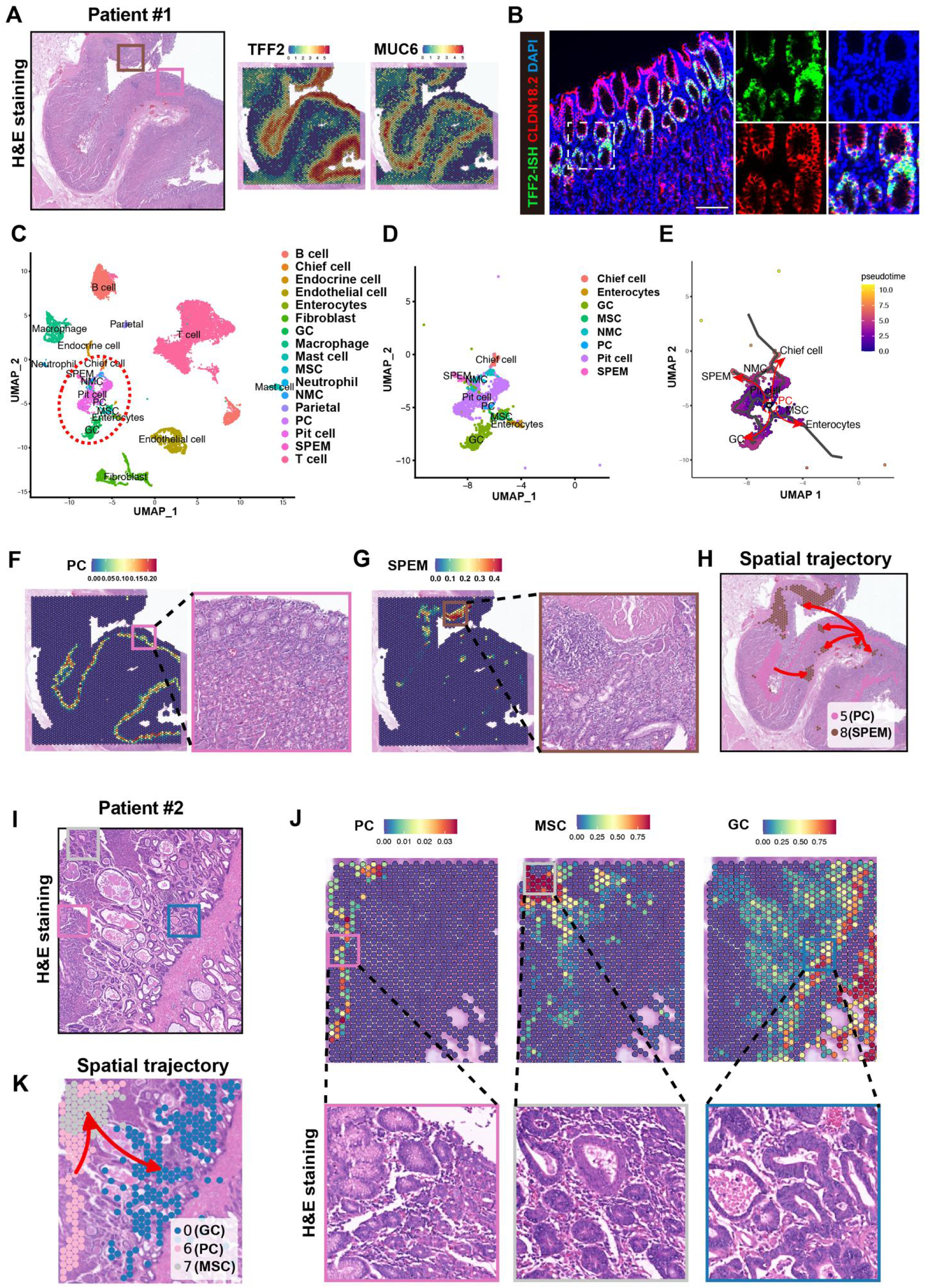
SPEM and gastric cancer arise from the isthmus but follow distinct differentiation paths in humans. **(A)** (Left) H&E staining of human gastric corpus section used for spatial transcriptomics (Patient #1). *TFF2* (middle) and *MUC6* (right) mRNA expression levels and localization in corpus tissues in spatial transcriptomics (Patient #1). Each spot represents 2-10 cells on the corresponding HE section. Color-coded values (from blue to red) reflect low to high levels of expression. Brown box indicates SPEM area, pink box indicates normal gland. (**B**) In situ hybridization of *TFF2* (green) and immunofluorescence staining for CLDN18.2 (red), and DAPI (blue) in the human normal corpus glands. White dashed box indicates *TFF2* mRNA expressed in the isthmus region. Scale bars, 100 μm. (**C** and **D**) scRNA-seq of gastric cancer in corpus (n=3) and adjacent mucosal tissue (more than 5 cm from the cancer) (n=2) from human patients. (C) After distinguishing normal epithelial cells from cancer cells, cell clusters identified based on cell type-specific markers. Red dashed box indicates epithelial clusters (without endocrine or parietal cells), and subset as shown in (D). (**E**) Monocle3 trajectory analysis. PC differentiate into normal epithelium (pit cell, NMC, chief cell), SPEM, enterocytes (mature IM cells) and GC in four directions. MSC transformed from PC and are the origin of enterocytes and GC. PC, normal proliferative cell cluster; MSC, metaplastic stem-like cells; GC, gastric cancer; NMC, mucous neck cell. (**F** and **G**) (Left) The spatial distribution of PC (F) and SPEM (G) defined by integration result by transferring the scRNA-seq dataset to the spatial transcriptomics dataset (Patient #1). (Right) High magnification details of the H&E images for validation of results in (left) panels. Color-coded values (from blue to red) reflect low to high predicted values. (F) Pink box indicates PC is mapped to the isthmus of human gastric coprpus and confirmed by H&E. (G) Brown box indicates normal corpus gland replaced by SPEM. (**H**) Spatio-temporal trajectory of SPEM progression based on the spatial changes between different cell states within a corpus tissue (Patient #1). The cluster 5 and 8 in spatial transcriptomics can be defined as PC and SPEM, respectively, by integrating information from H&E, scRNA-seq dataset, clustering distribution, and lineage specific gene expression. As predicted by stLearn, it runs from the PC (cluster 5) to the SPEM (cluster 8). (**I**) H&E staining of human gastric corpus cancer section used for spatial transcriptomics (Patient #2). (**J**) (Top) The spatial distribution of PC (left), MSC (middle), and GC (right) defined by integration result by transferring the scRNA-seq dataset to the spatial transcriptomics dataset (Patient #2). (Bottom) High magnification details of the H&E images for validation of results in (top) panels. Color-coded values (from blue to red) reflect low to high predicted values. Pink box indicates PC is mapped to the isthmus of human gastric corpus and confirmed by H&E. Gray box indicates MSC also map to the isthmus, but H&E shows that the cellular morphology is different from that of the PC. Blue box indicates defined GC clusters and confirmed by H&E. (**K**) Spatio-temporal trajectory of gastric cancer progression based on the spatial changes between different cell states within a cancerous tissue (Patient #2). The cluster 0, 6 and 7 in spatial transcriptomics can be defined as GC, PC and MSC, respectively, by integrating information from H&E, scRNA-seq dataset, clustering distribution, and lineage specific gene expression. As predicted by stLearn, it runs from the PC (cluster 6) through the MSC (cluster 7) and then the GC (cluster 0).

To investigate cell differentiation trajectories, we analyzed scRNA-seq datasets from five gastric corpus cancer cases and their adjacent tissues (Fig. 7C, Fig.S12C, S12D). Using cell origin, copy number variation (CNV), and k-means clustering (Fig.S13A-C), we distinguished GC from normal epithelial cells and identified these epithelial cell types based on marker genes (Fig.S13D). For the differentiation trajectory of metaplasia and tumors, we focused on key cell types: gastric cancer cells (GCs), proliferating cells (PCs) representing isthmus progenitor cells, metaplastic stem-like cells (MSCs), mucous neck cells (MNCs), pit cells, chief cells, SPEM, and enterocytes (mature intestinal metaplasia cells or IM) (Fig. 7D). Monocle3 trajectory analysis predicted that proliferating cells (PCs) serve as the origin of all these cell types, differentiating along distinct pathways into chief cells, SPEM, enterocytes, and GCs (Fig. 7E). Notably, PCs first transition into MSCs before developing into IM or GCs (Fig. 7E).

To map the spatiotemporal trajectories of cell types in situ, we integrated 10× Genomics Visium Spatial Gene Expression data with human scRNA-seq deconvolution results. The spatial distribution of PCs, SPEM, MSCs, and GCs corresponded well with mapped H&E images and marker gene expression (Fig.7F-J, Fig.S13E-G). Unlike scRNA-seq, where each spot represents a single cell, spatial transcriptomics captures data from spots containing 2–10 cells, meaning that clusters in spatial transcriptomics with mixed cell types. To determine the spatial distribution of cell types of interest, we made a comprehensive decision based on Louvain clustering (Fig.S13E-G, Fig.S14A-E), cell morphology observed in H&E images (Fig.7F-J), and marker gene expression patterns (Fig.7A).

To confirm the differentiation trajectory of SPEM, we selected a gastric tissue section (Patient #1) containing PCs, SPEM, and chief cells. PCs were predominantly localized to spatial cluster 5, SPEM to cluster 8, and chief cells to cluster 0 (Fig.7A, 7F-G, Fig. S13F, S13G, and Fig.S14A, S14B). Pseudo-spatiotemporal trajectory analysis using stLearn predicted that SPEM originates from PCs (cluster 5) and not from chief cells (cluster 0) (Fig.7H). To trace the differentiation trajectory of GC, we examined a tissue section containing PCs, MSCs, SPEM, and GC. PCs were localized to spatial cluster 6, MSCs to cluster 7, GCs to cluster 0, and SPEM to cluster 5 (Fig. 7I, 7J, Fig. S13E, S13F, and Fig.S14C-E). The predicted spatiotemporal trajectory of GC followed a path from PCs (cluster 6) to MSCs (cluster 7) and then to GCs (cluster 0), bypassing SPEM (cluster 5) (Fig.7K). In sum, both SPEM and GC appear to arise from the isthmus in the human gastric corpus, but follow distinct differentiation trajectories.

## Discussion

Our study identifies Tff2+ corpus isthmus cells as TA progenitors and the cellular origin of SPEM and dysplasia in the gastric corpus. Using a *Tff2-T2A-CreERT2* knock-in mouse model, we showed that Tff2+ progenitors were highly proliferative and under normal homeostatic conditions give rise to multiple glandular secretory cells (e.g. neck, chief and parietal) but lacked long-term self-renewal. However, upon Kras mutation, Tff2^+^ progenitors exhibit some plasticity and acquire stem-like properties and contribute to the development of dysplasia. Notably, the more differentiated SPEM cells exhibited limited proliferative potential and failed to progress to dysplasia even in the presence of oncogenic Kras, challenging our understanding of the prevailing stepwise model of inflammation-metaplasia-dysplasia progression. These findings highlight the pivotal role of isthmus stem/progenitor cells in linking injury, metaplasia and carcinogenesis and provide a clearer understanding of the cellular transitions that occur during progression from injury to carcinogenesis in the gastric epithelium.

Previous studies have suggested that chief cells dedifferentiate, regain stem cell-like properties, and initiate the expression of SPEM markers such as TFF2 and MUC6.^14,16,24–26^ This hypothesis, supported by acute injury models involving chemically induced parietal cell loss, posits that chief cells have the potential to transdifferentiate into SPEM. However, our results provide compelling evidence that Tff2+ progenitors, rather than chief cells, are the primary contributors to SPEM formation following both acute and chronic gastric injury. Notably, complete ablation of chief cells did not interfere with SPEM development or the glandular repair process. Furthermore, genetic tracing of Gif+ chief cells reconfirmed their fate of loss following HDT-induced acute injury or Kras mutation.^13^

Consistent with oncogenic transformation following Kras activation of isthmus stem cells,^27^ our data indicate that Tff2^+^ progenitors are also susceptible to oncogenic progression, as evidenced by their direct contribution to dysplasia following Kras activation. While Kras mutation alone was sufficient to induce dysplasia in Tff2^+^ progenitors, a process accelerated by chief cell loss, SPEM cells remained largely arrested even in the presence of oncogenic Kras. This suggests that the progression from inflammation to dysplasia may involve direct oncogenic transformation of stem-like progenitors, bypassing intermediate metaplastic stages. These findings are consistent with emerging evidence that stem/progenitor cells within the gastric isthmus are the predominant source of dysplastic lesions, particularly under conditions of *H. pylori*-induced inflammation.^4,27,40,41^

Our results some modification to the classical stepwise model of gastric carcinogenesis, which assumes a linear progression from chronic inflammation to metaplasia and finally tumorigenesis.^1,2^ While correct at the whole tissue level, we propose that dysplastic cells arise directly from isthmus progenitors, with SPEM representing an adaptive but largely non-proliferative and terminally differentiated lineage.^42^ This has significant implications for understanding the early stages of gastric cancer initiation, suggesting that identifying better markers for these early stem/progenitor cells within the isthmus, rather than focusing just on metaplasia, may be a more effective strategy to screen for patients at high risk for progression to cancer. Furthermore, the limited proliferative capacity of SPEM cells and their expression of TFF2 peptide, shown to be a tumor suppressor, suggests that they represent a protective response to injury rather than a high-risk, pre-neoplastic state. Thus, while SPEM often coexist with dysplastic lesions, our data support the view that it is the stem/progenitor cells, rather than SPEM itself, that drive neoplastic progression.

Although our study provides new insights into the cellular origins of SPEM and dysplasia, certain limitations must be acknowledged. First, although we demonstrate that Tff2^+^ isthmus cells are TA progenitors that can acquire stem-like properties and progress to dysplasia following Kras mutation, the precise molecular mechanisms driving this transformation remain to be elucidated. In addition, our studies indicate that Tff2^+^ progenitors are likely downstream of gastric stem cells, which could be the primary site of cancer initiation. Future studies should focus on identifying the key signalling pathways that regulate this transition, particularly in the context of chronic inflammation. Furthermore, while we have shown that Tff2^+^ progenitors are the immediate source of SPEM and dysplasia in our mouse models, further validation in human tissues is essential to confirm the clinical relevance of these findings. Advances in single-cell RNA sequencing and spatial transcriptomics provide promising tools for tracing lineage trajectories in human gastric stem/progenitor cells, which may help elucidate conserved mechanisms of gastric carcinogenesis.

In conclusion, our findings identify Tff2^+^ isthmus progenitors as key players in both the repair of gastric injury through SPEM formation and the generation of dysplasia through oncogenic transformation. These findings challenge the conventional view of SPEM as a precursor to dysplasia^5,8–11^ and suggest that future therapeutic approaches should focus on targeting stem/progenitor cells within the gastric isthmus to prevent cancer initiation. Further investigation of the molecular regulation of these stem/progenitor cells will be crucial to fully understand their role in gastric carcinogenesis and to develop effective preventive and therapeutic strategies.

### Limitations of the study

Our study identifies Tff2^+^ progenitors as an important cellular origin of SPEM and dysplasia, but several limitations remain. The precise molecular mechanisms driving the transformation of Tff2^+^ progenitors into dysplasia are not fully characterised. Although our findings are supported by human single-cell data, mouse models may not fully recapitulate human gastric pathology. Further validation in larger human cohorts and diverse genetic contexts is required to establish clinical relevance.

## ACKNOWLEDGMENTS

This research was funded by the NIH grants (to TCW) including grant numbers R35CA210088 and R01CA272901. Additional funded in part by the NIH/NCI Cancer Center Support Grant (P30CA013696), National Natural Science Foundation of China (to RT) (82403015) and utilized the resources of the Herbert Irving Comprehensive Cancer Center, including the Flow Cytometry Shared Resources, Molecular Pathology/MPSR, Genomics and High Throughput Screening, and the Genetically Modified Mouse Model Shared Resource (GMMMSR). The research was also supported from the Columbia University Digestive and Liver Disease Research Center (CU-DLDRC) grant 1P30DK132710 and its Bioimaging and Bioinformatics cores.

## AUTHOR CONTRIBUTIONS

Conceptualization, R.T. and T.C.W.; methodology, R.T. and T.C.W.; software, B.Z.; formal analysis, R.T., H.Z., B.Z., and Q.Z.; investigation, R.T., H.Z.,B.Z., Q.Z., J.Q., F.W., T.S., Y.O., H.K., Q.T.W., L.B.Z., S.T., S.M., Y.H., and T.C.W.; resources, C.H., and P.L.; writing-original draft, R.T.; writing-review & editing, R.T., H.Z.,B.Z., Y.H., and T.C.W.; supervision, T.C.W.; funding acquisition, R.T., and T.C.W.

## DECLARATION OF INTERESTS

The authors declare no competing interests.

## STAR★METHODS

### KEY RESOURCES TABLE

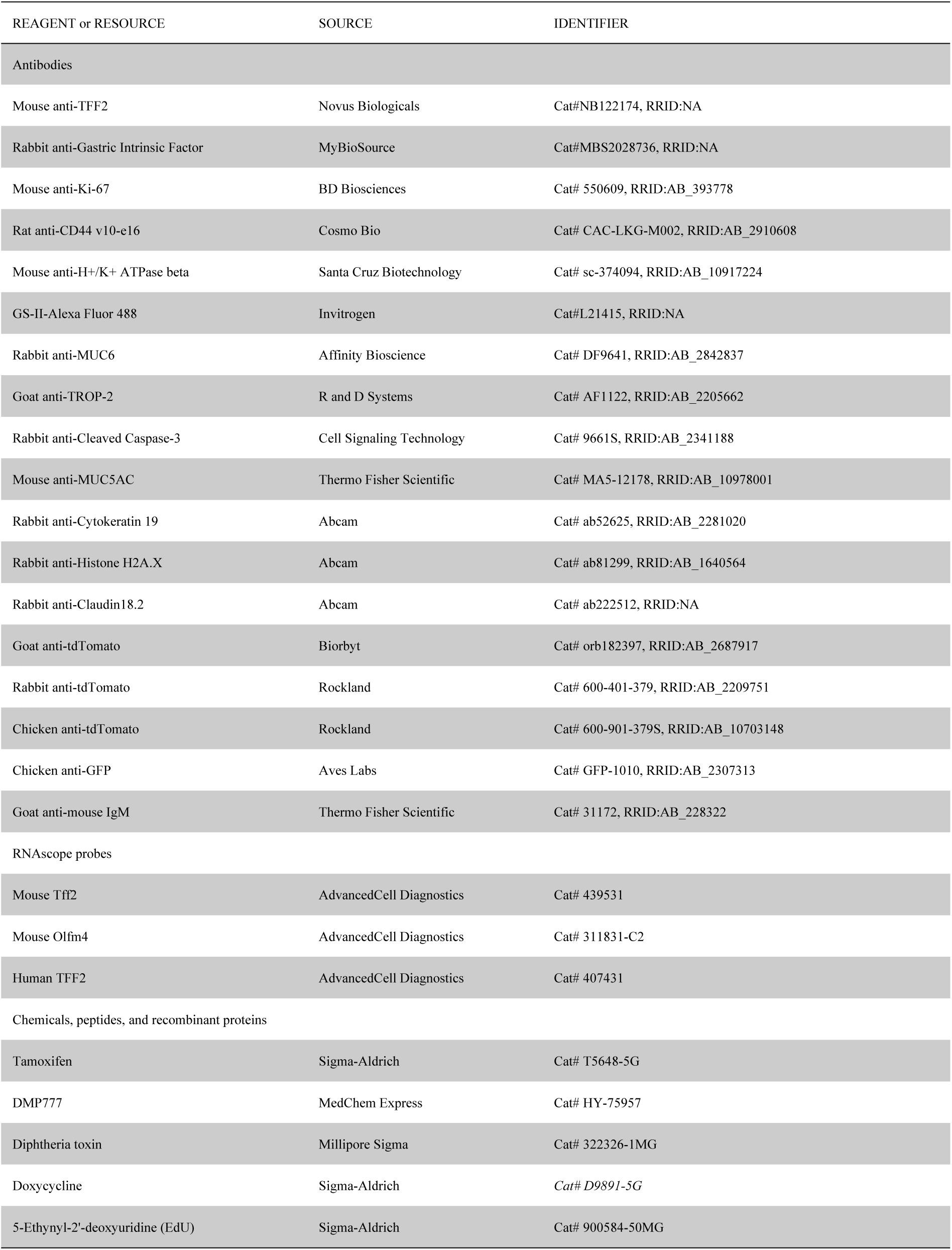
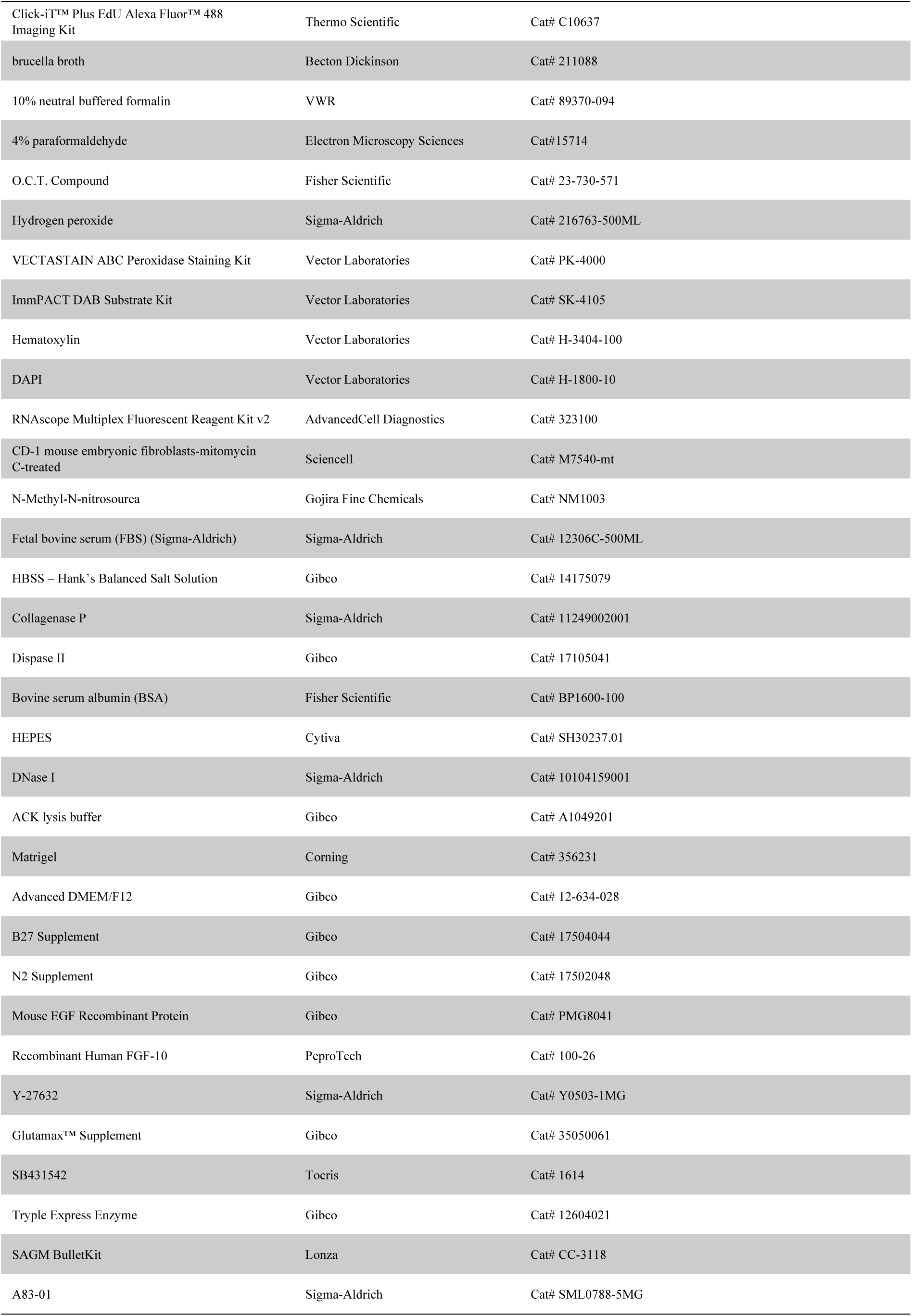
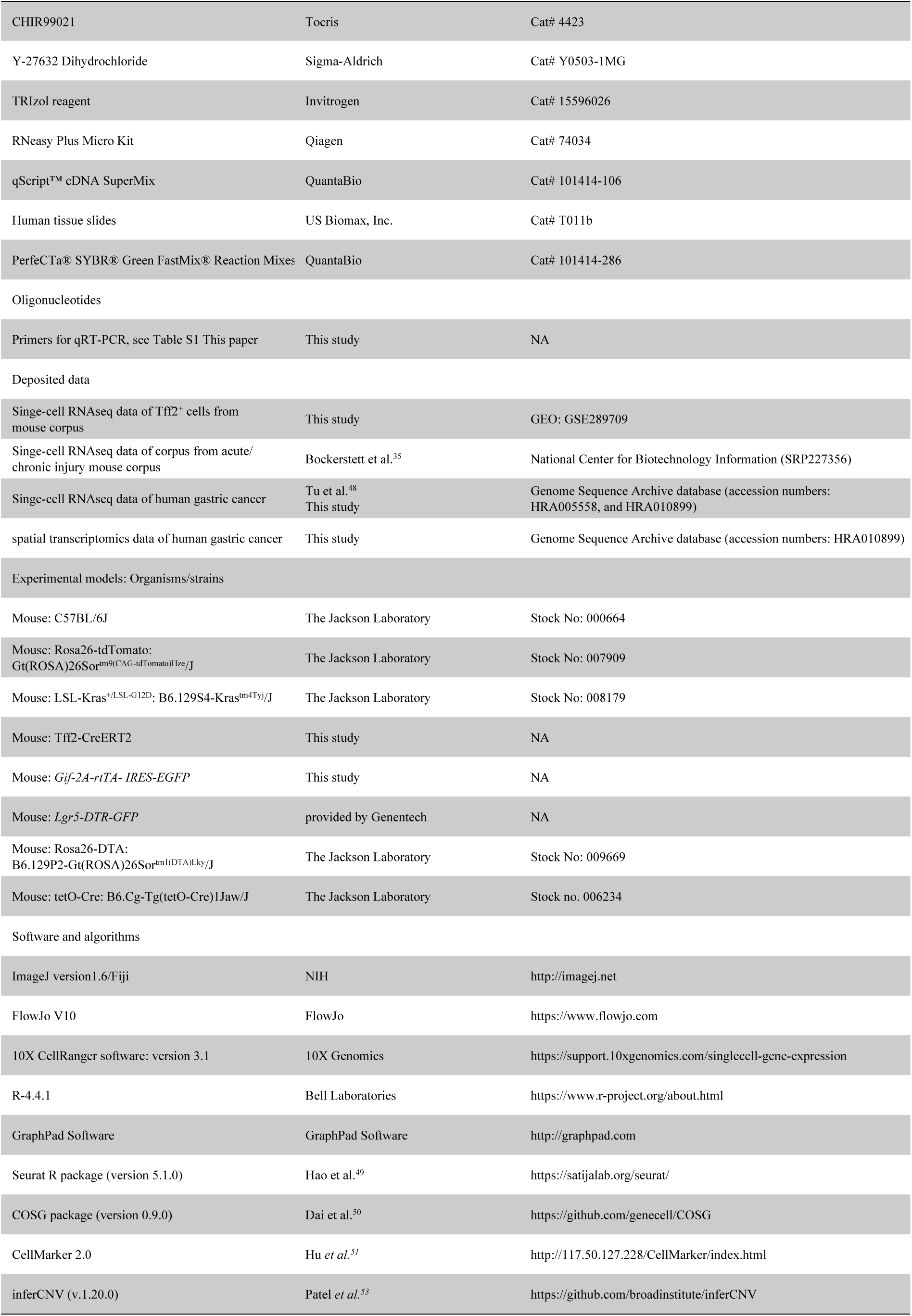
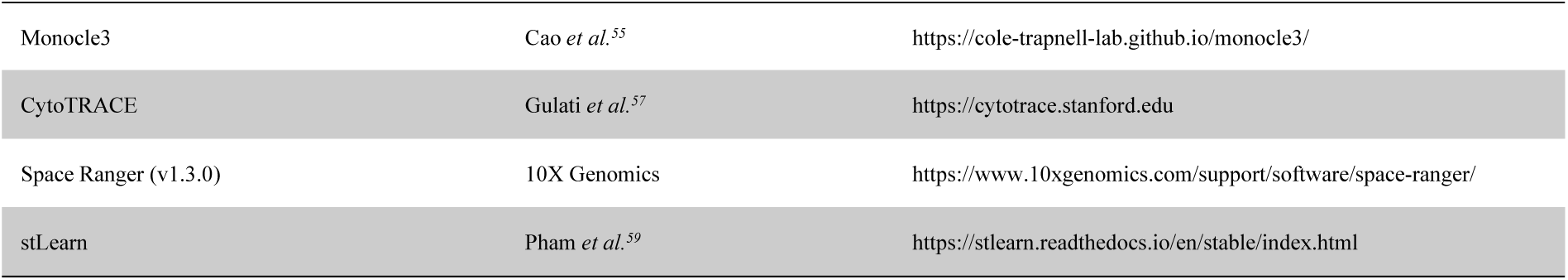

## RESOURCE AVAILABILITY

### Lead contact

Requests for resources and reagents should be directed to and will be fulfilled by the lead contact, Timothy C. Wang.

### Materials availability

Requests for mouse lines generated in this study should be directed to the lead contact and will be fulfilled. This study did not generate new unique reagents.

### Data and code availability

- The scRNA-seq dataset generated during this study is available in the Gene Expression Omnibus and Genome Sequence Archive database, and the accession number is listed in the key resources table.
- This paper does not report original code.
- Any additional information required to reanalyze the data reported in this paper is available from the lead contact upon request.

## EXPERIMENTAL MODEL AND SUBJECT DETAILS

### Animal models

All animal studies and procedures were reviewed and approved by the Columbia University Medical Center Institutional Animal Care and Use Committee (IACUC)-approved protocol (AC-AABV0664), and all mice were housed in a specific pathogen free (SPF) conditions. Male and/or female animals were used for experiments.

*Tff2-T2A-CreERT2* knock-in mice were generated at the Columbia University Genetically Modified Mouse Models Core facility. RP24-335C10 Bacterial Artificial Chromosome (BAC) clone carrying the *Tff2* gene was purchased from BACPAC Resource (BACPAC Genomics, Inc., Emeryville, California, USA.). This *Tff2* BAC clone was transferred into SW106 bacteria strain and modified into the *Tff2-T2A-CreERT2-FrtNeoFrt* BAC by BAC recombineering method. The *pMCS-Tff2-T2A-CreERT2-DTA* gene targeting vector was constructed by retrieving 1991 bp left arm-^43^-3277 bp right arm from the *Tff2-T2A-CreERT2-FrtNeoFrt* BAC into *pMCS-DTA* plasmid to generate the *pMCS-Tff2-T2A-CreERT2-DTA* gene targeting vector carrying a *DTA* (Diphtheria Toxin Alpha chain) negative selection marker. The FNF cassette confers G418 resistance in KV1 (129B6 hybrid) ES cells and the DTA cassette provides an autonomous negative selection to reduce the random integration event during gene targeting. Two targeted ES cell clones were identified. These two targeted ES cells were injected into C57BL/6J blastocysts to generate chimeric mice. Male chimeras should be bred to female *ACTB-Flpe* delete (*B6.Cg-Tg(ACTFLPe)9205Dym/J*, https://www.jax.org/strain/005703) to remove *Neo* cassette (A 73 bp sequence containing a Frt site is left behind) (Fig.1A). Positive founders were identified and backcrossed to C57BL/6J mice.

*Gif-2A-rtTA-IRES-EGFP* (referred to as Gif-rtTA) mice were generated at the Laboratory Animal Resource Center in Transborder Medical Research Center of University of Tsukuba. We selected a sequence (5’-AAA CCA TCC TTT TGT CAC AT-3’) containing the termination codon of the Gif as the CRISPR target. We purchased synthetic crRNA containing this target in sequence from IDT (Iowa, US). In the pGif-EGFP-rtTA donor DNA, we placed the IRES-EGFP-GSGP2A-rtTA Advance-rabbit beta-globin polyadenylation signal sequence between the 5’ and 3’ homology arms. The 5’-homology arm is the genomic region from 1,262 bp upstream to 20 bp downstream of the Gif termination codon, and the 3’-homology arm is the genomic region from 21 bp downstream to 1,377 bp downstream of the Gif termination codon. The CRISPR-Cas9 ribonucleoprotein complex and each donor DNA were microinjected into zygotes of C57BL/6J mice (Jackson Laboratory Japan, Kanagawa, Japan) according to our previous report.^44^ Subsequently, microinjected zygotes were transferred into oviducts in pseudo-pregnant ICR female (Jackson Laboratory Japan, Kanagawa, Japan) and newborns were obtained. To confirm the designed knock-in allele, the genomic DNA were purified from the tail with PI-200 (KURABO INDUSTREIS LTD, Osaka, Japan) according to manufactureŕs protocol. Genomic PCR was performed with KOD-Fx (TOYOBO, Osaka, Japan). The primers (Gif screening 5F: 5’-CGC TGA AAC TTC TTC CCA CTT AAA GAG C-3’ and Gif screening 3Rv: 5’-CTT TGG ATT CTG CAC CCT CCT GTT TAT T-3’) were used for detecting the correct knock-in allele. In addition, we checked random integration of donor DNA by PCR with donor vector backbone detecting primer (Donor detect F2: 5’-TGT GGA ATT GTG AGC GGA TA-3’, and Gif donor detect R: 5’-TGG CAC ATA GGA ACC ATT CA-3’).

*Lgr5-DTR-GFP* mice, which have been previously reported,^45–47^ were provided by Genentech. *B6.129P2-Gt(ROSA)26Sor^tm^*^1^*^(DTA)Lky^/J* (stock no. 009669, referred to as DTA) mice*, B6.129S4-Kras^tm4Tyj^/J* (stock no. 008179, referred to as Kras^G12D^) mice*, B6.Cg-Gt(ROSA)26Sor^tm^*^9^*^(CAG-tdTomato)Hze^/J* (stock no. 007909, referred to as ROSA26-tdTomato) mice, *B6.Cg-Tg(tetO-Cre)1Jaw/J* (stock no. 006234, referred to as tetO-Cre) mice, C57BL/6J (stock no. 000664, referred to as WT) mice were purchased from The Jackson Laboratory.

*Tff2-CreERT* mice were treated with tamoxifen (75 mg/kg, intraperitoneal injection (IP), diluted in 200 μl corn oil, Sigma-Aldrich, Cat# T5648-5G) to induce nuclear transfer of Cre recombinase. *Gif-rtTA;tetO-Cre* mice were induced by administering doxycycline (Sigma-Aldrich, Cat# D9891-5G) in drinking water at a concentration of 0.2 mg/mL for one week, followed by a switch to doxycycline-free water for two weeks to ensure that the tet system was completely inactivated before subsequent treatments.*Gif-rtTA;tetO-Kras^G12D^* mice were fed with doxycycline throughout the experiment. Mice were examined daily for health conditions and euthanized at the indicated time points by CO2 asphyxiation and cervical dislocation. For all experiments, mice were randomly allocated to experimental groups. Unless otherwise noted, experiments were performed in 3 biological replicates per condition.

### Human samples

5 samples from 4 patients of fresh gastric tissues for single cell RNA-seq, 24 samples of gastric cancer FFPE section for spatial transcriptomics, and 188 pairs of tissue microarrays (TMA1-4) consisting of gastric adenocarcinoma specimens and their corresponding non-cancerous tissues were obtained from patients who underwent gastrectomy at Fujian Medical University Union Hospital (FJMUUH) between January 2010 and June 2012. Written informed consent was obtained from all participants. The design and reporting procedures of this study were approved by the Institutional Review Board of Fujian Medical University Union Hospital (IRB number:2021KY042-03). Human tissue slides containing cases of gastric cancer and normal stomach tissue were also purchased from US Biomax, Inc. (Cat# T011b).

## METHOD DETAILS

### In vivo experiments

*Induction of acute injury.* Age and sex matched 6-to 8-week-old mice were received 2 injections of high-dose tamoxifen (HDT, 300mg/Kg, IP once a day for 2 days, diluted in 200 μl corn oil) or one injection of DMP777 (500 mg/Kg, oral gavage, diluted in 200 μl corn oil, MedChem Express, Cat# HY-75957). Control mice were injected with vehicle.

*Lgr5-DTR-GFP* mice or *Tff2-CreERT; Lgr5-DTR-GFP; tdTomato* mice were treated with two doses of HDT as well as two doses of diphtheria toxin (DT, 50 μg/kg, IP, Millipore Sigma, Cat# 322326-1MG), which allowed simultaneous induction of acute injury and ablation of Lgr5^+^ chief cells. The first dose of tamoxifen and the first dose of DT were administered on the same day, with a second dose of DT administered on day 4 to prevent later regeneration of chief cells.

*Tff2-CreERT2; DTA; tdTomato* mice were treated with two doses of HDT, allowing simultaneous tracing and ablation of Tff2^+^ progenitors after tamoxifen induction.

*5-Ethynyl-2’-deoxyuridine (EdU) labeling tracing.* 6-to 8-week-old WT mice were treated with EdU (50 mg/kg, IP, Sigma-Aldrich, Cat# 900584-50MG) to label proliferating cell 2 hours before last dose of tamoxifen. Mice were sacrificed and examines tissue sections at the indicated time points, stained according to Click-iT™ Plus EdU Alexa Fluor™ 488 Imaging Kit (Thermo Scientific, Cat# C10637) following the manufacturer’s protocol.

*H. pylori infection.* 6-to 8-week-old *Tff2-CreERT; Kras^LSL-G12D^; ROSA26-tdTomato* mice or *Tff2-CreERT; ROSA26-tdTomato* mice were infected by oral gavage with *H. pylori* (SS1, 2×10^9^ CFU, every 2 days×3 doses, from) in 0.2ml brucella broth (Becton Dickinson, Cat# 211088).

### Histopathology, immunohistochemistry, immunofluorescence, and in situ hybridization

*Histopathology.* At the time of harvest, the stomach was opened along the greater curvature and laid flat. For the preparation of paraffin sections, tissue specimens were fixed overnight in 10% neutral buffered formalin (VWR, Cat# 89370-094), transferred to 70% ethanol, and then embedded in paraffin. For the preparation of frozen sections, tissue specimens were fixed in 4% paraformaldehyde (Electron Microscopy Sciences, Cat#15714) for 24 hours, infiltrated by 30% sucrose and embedded in O.C.T. Compound (Fisher Scientific, Cat# 23-730-571). The blocks of samples were then serially sectioned on edge at 4 μm thickness, and one section from each series was stained with hematoxylin and eosin (H&E), and H&E images were captured using a Leica Aperio AT2 DX System by the Columbia Molecular Pathology/MPSR (HICCC) core. The remaining sections were kept for immunofluorescence, immunohistochemistry, or in situ hybridization analysis.

*For immunohistochemistry staining*, endogenous peroxidases were blocked by incubation with 3% hydrogen peroxide (Sigma-Aldrich, Cat# 216763-500ML) and non-specific signals were blocked using 4% BSA, 2% donkey serum, and 0.15% Triton. Primary antibodies were applied and incubated overnight at 4°C. ImmPress HRP IgGs (Vector Laboratories) were used as secondary antibodies, followed by VECTASTAIN ABC Peroxidase Staining Kit (Vector Laboratories, Cat# PK-4000) and ImmPACT DAB Substrate Kit (Vector Laboratories, Cat# SK-4105) for detection. The nuclei were counterstained with hematoxylin (Vector Laboratories, Cat# H-3404-100). Images were scanned with a Leica Aperio AT2 DX System.

*For immunofluorescence staining*, non-specific signals were blocked using 4% BSA, 2% donkey serum, and 0.15% Triton. Primary antibodies were applied and incubated overnight at 4°C. Secondary antibodies conjugated with Alexa Fluor-488, Alexa Fluor-555 or Alexa Fluor-647 (Thermo Fisher Scientific) were used. The nuclei were then counterstained with DAPI (Antifade Mounting Medium With DAPI, Vector Laboratories, Cat# H-1800-10). Images were captured with a SPOT Pursuit Color digital camera, using a Nikon Eclipse TE2000 Inverted Microscope.

*Primary antibodies*: TFF2 (Novus Biologicals, Cat#NB122174, RRID:NA), Gastric Intrinsic Factor (GIF, MyBioSource, Cat#MBS2028736, RRID:NA), Ki-67 (BD Biosciences, Cat# 550609, RRID:AB_393778), CD44 v10-e16, ortholog of human v9 (Cosmo Bio, Cat# CAC-LKG-M002, RRID:AB_2910608), H+/K+ ATPase beta (ATP4B, Santa Cruz Biotechnology, Cat# sc-374094, RRID:AB_10917224), GS-II-Alexa Fluor 488 (Invitrogen, Cat#L21415, RRID:NA), MUC6 (Affinity Biosciences, Cat# DF9641, RRID:AB_2842837), TROP-2(R and D Systems, Cat# AF1122, RRID:AB_2205662), Cleaved Caspase-3 (Cell Signaling Technology, Cat# 9661S, RRID:AB_2341188), MUC5AC (Thermo Fisher Scientific, Cat# MA5-12178, RRID:AB_10978001), Cytokeratin 19 (Abcam, Cat# ab52625, RRID:AB_2281020), Histone H2A.X (Abcam, Cat# ab81299, RRID:AB_1640564), Claudin18.2 (Abcam, Cat# ab222512, RRID:NA), tdTomato (Biorbyt, Cat# orb182397, RRID:AB_2687917), tdTomato (Rockland, Cat# 600-401-379, RRID:AB_2209751), tdTomato (Rockland, Cat# 600-901-379S, RRID:AB_10703148), GFP (Aves Labs, Cat# GFP-1010, RRID:AB_2307313). For TFF2 detection, goat anti-mouse IgM (Thermo Fisher Scientific, Cat# 31172, RRID:AB_228322) was used to combine with TFF2 antibody (IgM), then a third antibody conjugated with Alexa Fluor-488, Alexa Fluor-555 or Alexa Fluor-647 (Thermo Fisher Scientific) was used.

*Periodic acid-Schiff staining and Alcian blue (pH 2.5) staining* were performed on paraffin sections by the Columbia Molecular Pathology/MPSR (HICCC) Core. Images were captured using a Leica Aperio AT2 DX system.

Steiner staining (Abcam) was used to detect *H. pylori* infection on paraffin sections. Cell nuclei were then stained with hematoxylin (Vector). Images were captured using a Leica Aperio AT2 DX system.

*In situ hybridization,* RNA in situ detection using an RNAscope Multiplex Fluorescent Reagent Kit v2 (AdvancedCell Diagnostics, Cat# 323100) according to the manufacturer’s instructions. RNAscope probes against mouse Tff2 (AdvancedCell Diagnostics, Cat# 439531), mouse Olfm4 (AdvancedCell Diagnostics, Cat# 311831-C2), and human TFF2 (AdvancedCell Diagnostics, Cat# 407431) were used; positive and negative control probes were run in parallel and used according to the manufacturer’s instructions. In combination with in situ hybridization and immunofluorescence, a solution containing 4% BSA, 2% donkey serum, and 0.15% Triton was used to block non-specific signals after the C1 probe channel, or both the C1 and C2 probe channels, had been developed. Primary antibodies were then applied. Images were captured with a SPOT Pursuit Color digital camera, using a Nikon Eclipse TE2000 Inverted Microscope.

### In vitro experiments

#### Gastric gland isolation and single-cell sorting

The corpus tissues were dissected and washed repeatedly in cold PBS with 1% antibiotic-antimycotic (Gibco), then transferred to 1.5 mL microcentrifuge tubes and minced into small pieces. The tissues were incubated on a shaker for 20 minutes at 37°C in 20 mL Hanks’ balanced salt solution (HBSS) (Gibco) containing 1 mg/mL collagenase P (Sigma-Aldrich), 1 mg/mL dispase II (Gibco), 10 mg/mL bovine serum albumin (BSA) (Fisher Scientific), 10 mmol/L HEPES (Cytiva), and 2 U/mL DNase I (Sigma-Aldrich). The incubation was neutralized with 20 mL cold HBSS containing 5% fetal bovine serum (FBS) (Sigma-Aldrich), and the cell suspension was passed through a 40 µm filter and centrifuged at 500g for 5 minutes at 4°C. Red blood cells were lysed using ACK lysis buffer (Gibco). After several washes with cold PBS containing 2% FBS, the cells were transferred to a polystyrene tube with a cell strainer cap (Corning). The cells were then stained with EpCAM, CD44v9, or CD45, and DAPI (4’,6-diamidino-2-phenylindole, dihydrochloride) (BD) was used to exclude dead cells. Cell sorting was performed on a BD Influx cell sorter.

### 3D organoid cultures

A total of 2,000 single cells (see above) were mixed with 20 μL Matrigel (Corning) per well and plated in 24-well plates. After polymerization of the Matrigel, organoid cultures were maintained as previously described^39,48^ with some modifications. The basal culture medium consisted of 50% conditioned medium containing Wnt3A, R-spondin-1, and Noggin (WRN, ATCC) and 50% advanced DMEM/F12 (Gibco) supplemented with 20% FBS, B27 (1×, Gibco), N2 (1×, Gibco), and 1% antibiotic-antimycotic. In addition, the medium was supplemented with 50 ng/mL EGF (Gibco), 100 ng/mL FGF10 (Peprotech), 10 µmol/L Y-27632 (Sigma-Aldrich), 2 mmol/L GlutaMAX (Gibco), 10 mmol/L HEPES, and 10 µmol/L SB431542 (Tocris). Culture medium was replaced every other day, and organoids were passaged every 6 days at a 1:3 ratio by dissolving the Matrigel with TrypLE (Gibco) and mechanically disrupting the organoids by pipetting.

### 2D gastric epithelial culture

After sorting, a total of 2,000 single cells per flask were plated and cultured as previously described^39^ with some modifications. First, T25 flasks were coated with poly-lysine (Sigma-Aldrich) to enhance cell adhesion to plasticware. Next, 2×10⁵ CD-1 mouse embryonic fibroblasts-mitomycin C-treated (Sciencell, Cat# M7540-mt) were plated and cultured overnight in DMEM (Gibco) supplemented with 10% FBS. Gastric epithelial cells were plated on these T25 flasks the next day. Both mouse embryonic fibroblasts and gastric epithelial cells were maintained in 2D culture medium prepared with 30% 3D organoid basal culture medium and 70% SABM medium (Lonza) supplemented with 1 μmol/L A83-01 (Sigma-Aldrich), 1 μmol/L CHIR99021 (Tocris), and 10 μmol/L Y-27632 (Sigma-Aldrich). The culture medium was replaced every other day, and gastric epithelial cells were passaged at a ratio of 1:5 when they reached 70% to 80% confluence.

For pit cell differentiation experiments, sorted tdTomato^+^ or tdTomato^-^ epithelial cells were planted directly (see above) onto Lab-Tek® II Chambered Coverglass (Biolyst Scientific). After 7 days of growth, when distinct cell clusters were evident, the MNU group was treated with 20 nmol/L N-Methyl-N-nitrosourea (MNU, Gojira Fine Chemicals, Cat# NM1003) for 24 hours, followed by medium change to fresh culture medium for another 24 hours. The other three groups (tdTomato^+^ progenitor, tdTomato^+^ progenitor with Kras, and tdTomato^-^ epithelial cells) were left untreated. Immunofluorescence staining was performed on day 9.

### RNA extraction and real-time quantitative PCR (RT-qPCR)

Total RNA was isolated from corpus tissues or sorted single cells (see above) using TRIzol reagent (Invitrogen) and the RNeasy kit (Qiagen) according to the manufacturer’s protocol. Purified RNA was then reverse transcribed into cDNA using a qScript™ cDNA SuperMix (QuantaBio). RT-qPCR analyses were performed using PerfeCTa® SYBR® Green FastMix® Reaction Mixes (Quantabio) on a QuantStudio 3 Real-Time PCR System (Thermo Fisher Scientific). Relative mRNA expression was calculated using the 2(-ΔΔCt) method and normalized to Gapdh. The primer sequences used are listed in Supplementary Table 1.

### Processing of Single-cell RNA-sequencing data

#### Human scRNA-seq date

Within one hour of gastrectomy, small pieces of the tumor and adjacent tissues (more than 5 cm from tumor) were immediately harvested for labeling and processing as our previously described.^49^ Samples were digested to obtain single-cell suspensions, which were then loaded into the wells of a microfluidic chip to generate a cDNA library using the 10x Genomics droplet-based sequencing platform. Single-cell transcriptome amplification and library preparation were performed according to the manufacturer’s instructions using Chromium Single Cell 3’ v2 chemistry (10x Genomics). Libraries were then pooled and sequenced on six lanes on an Illumina HiSeq X-10 system.

The raw sequencing data were aligned and quantified against the human reference genome (hg19) using CellRanger Single Cell Software Suite (version 3.1). The Seurat R package (version 5.1.0)^50^ was used to import and process the gene expression matrices. Initial preprocessing thresholds included genes detected in at least three cells and cells with at least 200 genes. Quality filtered using the following criteria: nGene > 500 and < 6000, nUMI > 600 and < 20,000, and mitochondrial genome < 25%. This filtering resulted in 14,464 cells left to analyze. The normalized data function was used to normalize the data. Principal component analysis (PCA) was performed on the top 2000 highly variable genes, and clustering analysis was performed on the top 30 PCA. Marker genes were identified using the COSG package (version 0.9.0)^51^ and then manually annotated based on differentially expressed genes (DEGs) references from previous literature and CellMarker 2.0.^52^ We defined these clusters into ten different cell types: epithelium (EpCAM, KRT18, and KRT8), parietal cell (ATP4A), endocrine cell (CHGA and CHGB), fibroblast (ACTA2 and COL1A2), endothelial cell (ENG and VWF), B cell (MS4A1 and CD79A), mast cell (CPA3), T cell (CD2, CD3D, and CD3E), macrophage (CD14, C1QA, and C1QB), and neutrophil (FCGR3B).

The identification of malignant and non-malignant epithelial cells was based on a previous scRNA-seq study of human gastric cancer,^53^ which used a k-means clustering algorithm. First, all epithelial cells were categorized as tumor or normal epithelial cells based on whether they were derived from tumor tissue or adjacent mucosa. The DEGs between the two groups were then calculated. The scores derived from the k-means clustering algorithm were used to redefine the putative malignant and non-malignant epithelial cells, followed by recalculation of the DEGs. This process was repeated iteratively until the classification results stabilized.

We also used inferCNV (v.1.20.0)^54^ to infer large-scale CNVs in each epithelial cell, using fibroblasts and endothelial cells as reference controls. Gene expression for each cell was re-standardized, with values constrained between −1 and 1. The CNV score for each cell was then calculated as the quadratic sum of CNVs.^55^

After defining the gastric cancer (GC) subcluster, other epithelial cells were easily defined based on marker genes and previous studies, such as pit cell (MUC5AC and TFF1), chief cell (PGA4 and PGA3), mucous neck cell (MUC6), SPEM (MUC6 and TFF2), proliferating cell (PC) representing isthmus cell (MKI67 and TOP2A), metaplastic stem-like cell (MSC) (OLFM4 and REG4^36–38^), and enterocyte as a mature intestinal metaplastic cell (FABP1^36,37^).

Trajectory analysis was performed on the dataset using Monocle3.^56^ The default parameters were used. The embeddings of the Monocle3 cell dataset object were replaced with the Seurat UMAP embeddings for consistency. The root cells of the cell dataset object were manually set as PC.

### Mouse scRNA-seq date

The Seurat R package (version 5.1.0) was used for the processing with analysis pipelines similar to those described in the human scRNA seq. Quality filtered using the following criteria: nGene > 250 and < 6000, nUMI > 500 and < 20,000, and mitochondrial genome < 25%. We sorted Tff2^+^ progenitors at day 1 (n=3) and their progeny at day 28 (n=3) from *Tff2-CreERT; Rosa26-tdTomato* mice treated with low-dose tamoxifen, using EpCAM^+^ and tdTomato^+^ markers as controls (Ctrl). Similarly, we sorted Tff2^+^ progenitors on day 1 (n=3) and their progeny on day 5 (n=3) from mice treated with high-dose tamoxifen to model acute injury (HDT). After removing low-quality and mixed cell populations, we obtained 13628 cells (Ctrl, n=5469; HDT, n=8159) and classified them into 5 clusters according to marker genes: progenitor cells (*Mki67*, and *Top2a*); neck mucous cells (*Muc6*); chief cells (*Gif(Cblif)*); parietal cells (*Atp4b*) and SPEMs (*Muc6*; Tff2; and *Gif*). The other mouse gastric corpus single-cell data were obtained from a public database.^35^ This is a sc-RNA seq analysis containing only corpus glands from acute (high dose tamoxifen induced acute gastric injury in BALB/c mice, n=2), chronic injury (autoimmune gastric metaplasia in the TxA23 mice, n=3) and nontreated WT mice (BALB/c mice, n=2). This filtering resulted in 21,118 cells left to analyze. We defined these clusters into 14 different cell types: isthmus cell (Mki67 and Top2a), chief cell (Gif), parietal cell (Atp4a), mucous neck cell (Muc6), SPEM (Muc6, Gkn3 and Tff2), pit cell (Muc5ac and Tff1), tuft cell (Dclk1), enteroendocrine cell (Chga and Chgb), transitional basal cell (Krt5^57^), fibroblast (Acta2 and Col1a2), endothelial cell (Eng and Vwf), B cell (Ms4a1 and Cd79a), T cell (Cd2, Cd3d, and Cd3e), and macrophage (Cd14, C1qa, and C1qb).

Monocle3 trajectory analysis predicted that the progenitors serve as the origin of all other cell types, including chief cells and SPEMs. CytoTRACE^58^ has been used with default parameters for the inference of cellular differentiation states. To investigate whether isthmus cells change after acute and chronic injury compared to the wild-type (WT) group, we used Seurat’s FindMarkers function to identify differentially expressed genes (DEGs) from our scRNA-seq data. We then performed Gene Set Enrichment Analysis (GSEA) on these DEGs using the hallmark gene sets (’mh.all.v2023.2.Mm.symbols.gmt’) obtained from the Molecular Signatures Database (MSigDB) to explore the associated biological processes.

### Processing of spatial transcriptomics data

Five consecutive 5 μm paraffin sections of gastric corpus tissue were selected per patient, with one section used for spatial transcriptomics sequencing analysis, one section for H&E staining and scanned using a Leica Aperio Versa 8 whole slide scanner (brightfield imaging settings, 20× magnification), and several sections kept in reserve. Section preparation and deparaffinization were performed according to the 10× Genomics protocol, followed by the addition of a human whole transcriptome probe set to the tissue. The probes generated a spatial transcriptomics library that was sequenced on the Illumina NovaSeq 6000 system (performed by Novogene Co., Ltd.). Raw FASTQ files and histological images were processed using the short-read probe alignment algorithm for the FFPE “Count” method in Space Ranger (v1.3.0) from 10× Genomics, with probe reads aligned to the human reference genome (GRCh38). Gene expression normalization, dimensionality reduction, spot clustering, and differential expression analysis were performed using the Seurat package (version 5.1.0). To define the spatial distribution of cell types identified by human scRNA-seq (described above), cell clusters were transferred from scRNA to spatial clusters using the TransferData function. Anchors were identified using the FindTransferAnchors function, and all variable features for transfer.^59^

To study the spatio-temporal trajectories of cells in situ, we integrated the raw spatial data with the deconvolution results (described above) in Python and performed further analyses following the protocol outlined in stLearn.^60,61^ After performing Louvain clustering, we performed pseudo-spatio-temporal trajectory analysis to reconstruct trajectories based on changes in transcriptional state across key clusters of interest, including PCs, MSCs, SPEM, and GC. In contrast to scRNA-seq, where each spot corresponds to a single cell, spatial transcriptomics captures data from 2-10 cells per spot. As a result, clusters in spatial transcriptomics can contain multiple cell types. The selection of clusters of interested cell type was guided by a combination of 1) cell type distributions identified by deconvolution within the Louvain clusters, 2) cell morphology observed in mapped H&E images, and 3) associated marker gene expression. For analyses of PC development and metaplasia or tumor progression, we manually set PCs as root cells.

## QUANTIFICATION AND STATISTICAL ANALYSIS

### Quantification

*For quantification of the number or ratio of single, double, and triple positive cells in IHC or IF staining,*^62,63^ images of corpus glands were taken at 10× or 20× magnification for each mouse. The total number of corpus glands and DAPI-positive glandular cells were counted using Fiji.^63^ We then calculated the staining intensity for each color channel and applied a uniform threshold to distinguish positive from negative cells, allowing us to count the number of positive individual cells. Regions of interest (ROIs) were created for each channel, encompassing all foreground pixels. Using the ‘AND’ option in the ROI manager, a new ROI was created containing only the overlapping regions of the selected ROIs, allowing us to count the number of double or triple positive cells. All vertically sectioned glands from longitudinal sections of the mouse stomach were included in the analysis. The investigator performing the image analysis was blinded to the experimental conditions.

*For quantification of the size and location of the tdTomato and EdU-positive gland compartments as previously described,*^30^ each gland was measured for its total height, the distance from the gland base to the first positive cell, the height of the positive gland compartment, and the distance from the most apical positive cell to the top of the gland. The ratio between these measurements was defined as the relative size of each gland compartment.

*For murine gastric histopathology scoring,* gastric corpus lesions (mucinous metaplasia, dysplasia) were scored on an ascending scale from 0 to 4 by a pathologist blinded to treatment group according to the criteria.^64^ The total number of corpus glands and lesions were also calculated, with the lesion rate defined as the ratio of lesions to the total number of glands.

*For quantification of organoid size,* 4× or 10×-magnification fields were analyzed with Fiji,^63^ expressing organoid areas as pixels. Images used for quantification were captured with a SPOT Pursuit Color digital camera, using a Nikon Eclipse TE2000 Inverted Microscope.

### Statistical analyses

Statistical analyses were performed with GraphPad Prism software. Data are presented as mean ± SD and significance was assessed using Student’s t-test for two group comparisons or one-way or two-way ANOVA with post hoc tests (such as Tukey’s or Dunnett’s) for multiple group comparisons. Differences with a p-value < 0.05 were considered statistically significant. All analyses included at least three independent biological replicates, unless otherwise stated.

## Notes

### Competing Interest Statement

The authors have declared no competing interest.

