## Supplemental Figures for "*Tff2* marks gastric corpus progenitors that give rise to pyloric metaplasia/SPEM following injury"

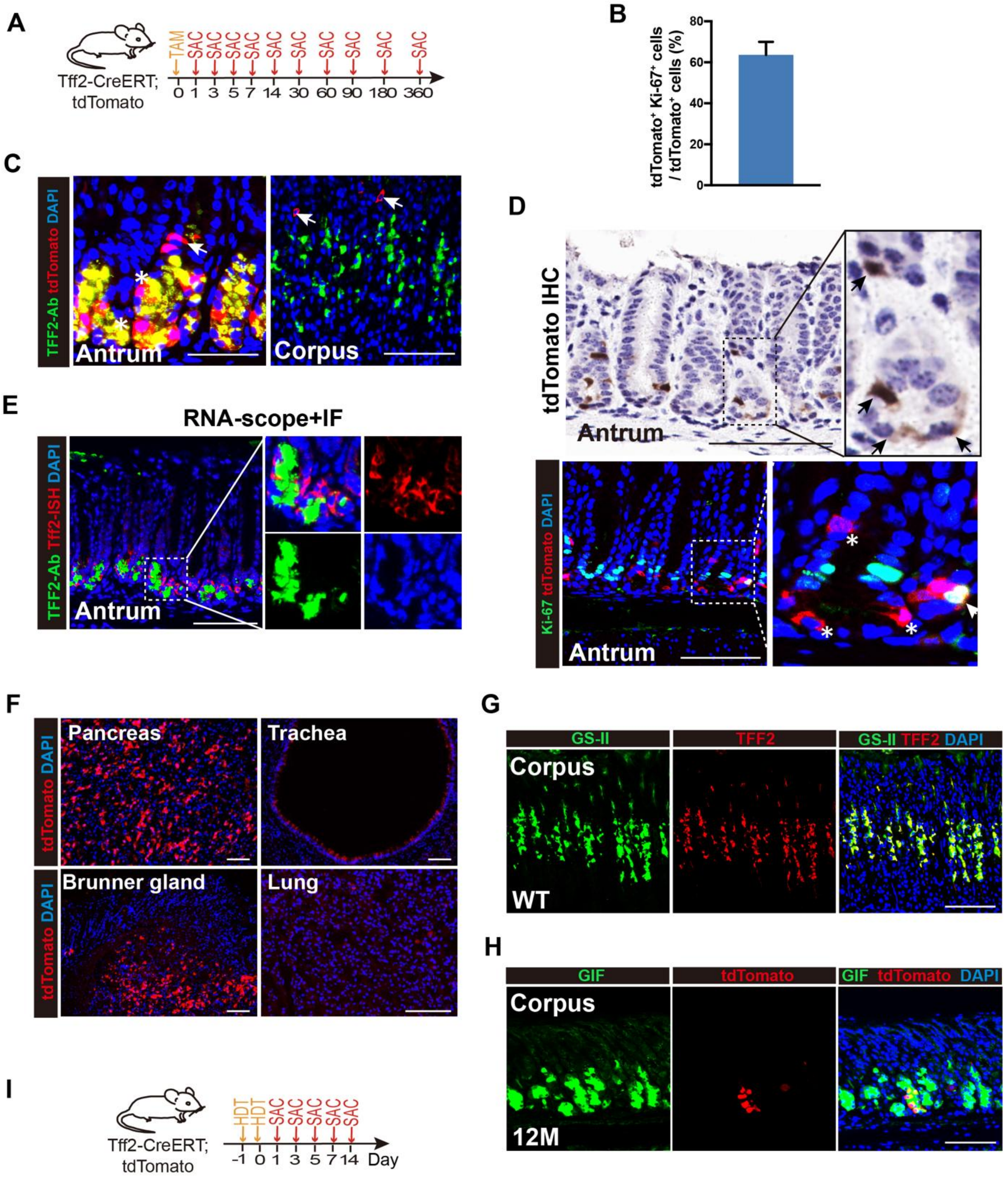


**Figure S1.** Tff2-CreERT⁺ cells also expressed in the antrum and other organs.

(**A**) Schematic of the experimental design for the short- and long-term lineage tracing of Tff2-CreERT⁺ progenitors in homeostasis. In brief, *Tff2-CreERT; R26-tdTomato* mice were treated with low-dose tamoxifen (75 mg/kg, one dose), and corpus tissues were sampled for histological analysis at different time points after induction. TAM, tamoxifen, SAC, sacrify. (**B**) Quantification of the percentage of tdTomato⁺ Ki-67⁺

cells to total tdTomato⁺ cells as in (Fig. 1B). Data are shown as mean ± SD; n = 3 mice per group. (**C**) Immunofluorescence staining for TFF2 (green), tdTomato (red), and DAPI (blue) in the antrum (left) and corpus (right) glands of *Tff2-CreERT2; R26-tdTomato* mice at 24 hours after tamoxifen (75 mg/Kg) induction. White arrow indicates tdTomato$^+$ cells. White asterisk indicates tdTomato$^+$ TFF2$^+$ cells. Scale bars, 100 μm. (**D**) (Top) Immunohistochemical staining for tdTomato (brown) in the antrum glands of *Tff2-CreERT2; R26-tdTomato* mice at 24 hours after tamoxifen (75 mg/Kg) induction. Black arrow indicates tdTomato$^+$ cell in the isthmus and base region. Scale bars, 100 μm. (Bottom) Immunofluorescence staining for Ki-67 (green), tdTomato (red), and DAPI (blue) in the antrum glands as in (top) panel. White arrow indicates tdTomato$^+$ Ki-67$^+$ cells in the isthmus region. White asterisk indicates tdTomato$^+$ cells in the isthmus and base region. Scale bars, 100 μm. (**E**) In situ hybridization of *Tff2* (green) and immunofluorescence staining for tdTomato (red), and DAPI (blue) in the antrum glands of *Tff2-CreERT2; R26-tdTomato* mice at 24 hours after tamoxifen (75 mg/Kg) induction. Scale bars, 100 μm. (**F**) Immunofluorescence staining for tdTomato (red) and DAPI (blue) in the pancreas (left top), trachea (right top), duodenal Brunner gland (left bottom), and lung (right bottom) of *Tff2-CreERT2; R26-tdTomato* mice at 24 hours after tamoxifen (75 mg/Kg) induction. Scale bars, 100 μm. (**G**) Immunofluorescence staining for GS-II (green), TFF2 (red) and DAPI (blue) in the corpus gland of nontreated wild-type (WT) mice, and showed complete overlap of both staining. Scale bars, 100 μm. (**H**) Immunofluorescence staining for GIF (green), tdTomato (red), and DAPI (blue) in the corpus glands of *Tff2-CreERT2; R26-tdTomato mice* at 12 months (12M) after tamoxifen (75 mg/Kg) induction. Scale bars, 100 μm. (**I**) Schematic of the experimental design for the short-term lineage tracing of Tff2$^+$ progenitors after high-dose tamoxifen (HDT)- induced acute injury. In brief, *Tff2-CreERT; R26-tdTomato* mice were treated with HDT (300 mg/kg, once daily for 2 consecutive days), and corpus tissues were sampled for histological analysis at different time points after induction.

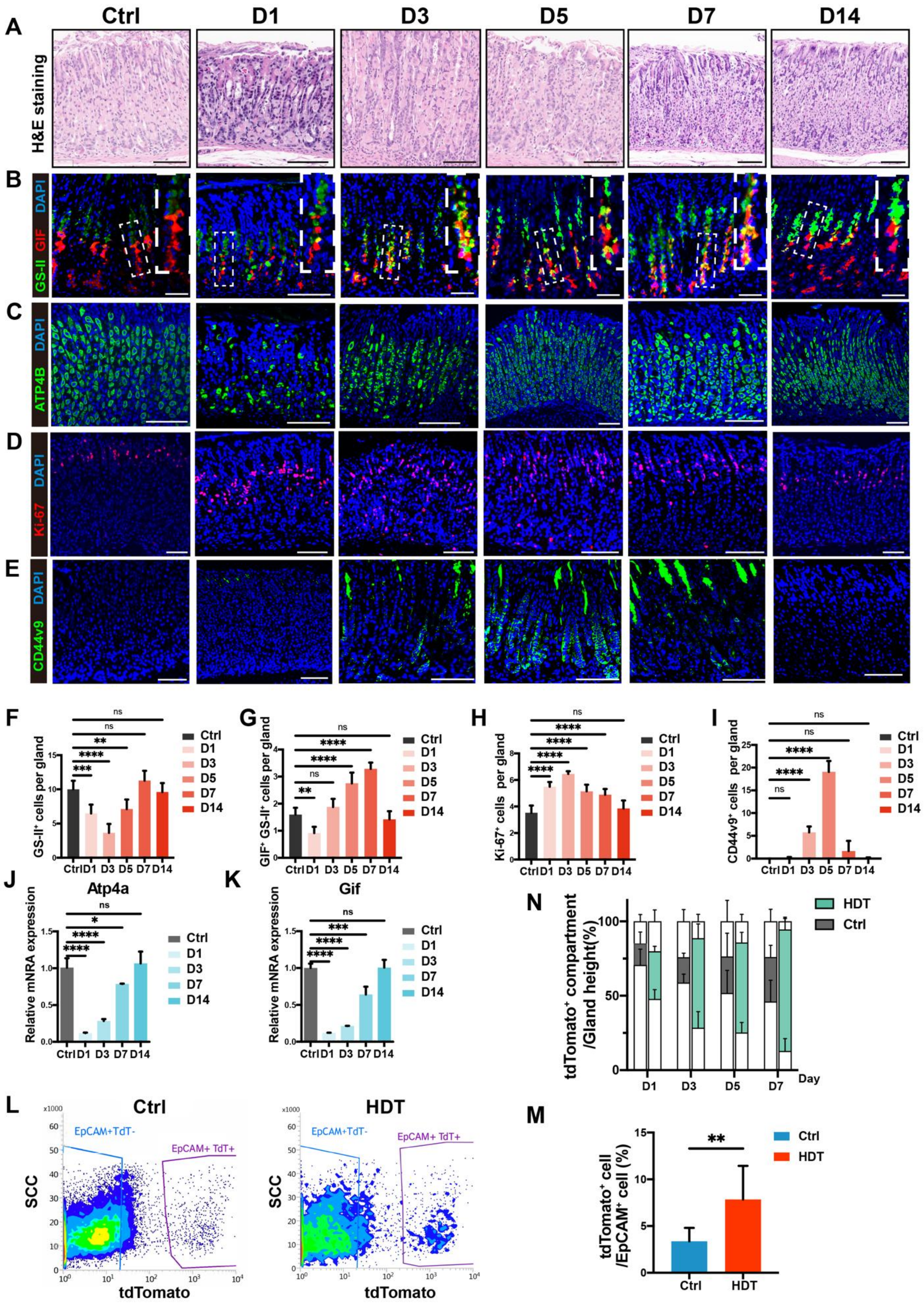


**Figure S2.** Acute corpus gland injury and repair processes induced by HDT.

(**A**) H&E staining in the corpus glands of WT mice before (Ctrl) and after HDT-induced acute injury (HDT, 300 mg/kg*2, once daily for 2 consecutive days) at different time points (D1, D3, D5, D7 and D14). Scale bars, 100 μm. (**B**) Immunofluorescence staining for GS-II (green), GIF (red), and DAPI (blue) in the corresponding groups described in (A). Insets indicates high magnification details of the white dashed box. Scale bars, 100 μm. (**C**) Immunofluorescence staining for ATP4B (green) and DAPI (blue) in the corresponding groups described in (A). Scale bars, 100 μm. (**D**) Immunofluorescence staining for Ki-67 (red) and DAPI (blue) in the corresponding groups described in (A). Scale bars, 100 μm. (**E**) Immunofluorescence staining for CD44v9 (green) and DAPI (blue) in the corresponding groups described in (A). Scale bars, 100 μm. (**F** and **G**) Quantification of the number of GS-$II^+$ cells per gland (F) and GS-$II^+$ $GIF^+$ cells per gland (G) as in (B). Data are shown as mean ± SD; n = 3 mice per time point per group. Statistical significance was assessed with one-way ANOVA. ns, not significant, **P < 0.01, ***P < 0.001, ****P < 0.0001. (**H**) Quantification of the number of Ki-$67^+$ cells per gland as in (D). Data are shown as mean ± SD; n = 3 mice per time point per group. Statistical significance was assessed with one-way ANOVA. ns, not significant, ****P < 0.0001. (**I**) Quantification of the number of $CD44v9^+$ cells per gland as in (E). Data are shown as mean ± SD; n = 3 mice per time point per group. Statistical significance was assessed with one-way ANOVA. ns, not significant, ****P < 0.0001. (**J** and **K**) Gene expression of parietal cell marker (*Atp4a*) (J) and chief cell marker (*Gif*) (K) in the corresponding groups described in (A). Expression is shown relative to before HDT induced group (Ctrl) and normalized to Gapdh. Data are shown as mean ± SD; n = 3 mice per time point per group. Statistical significance was assessed with one-way ANOVA. ns, not significant, *P < 0.05, ***P < 0.001, ****P < 0.0001. (**L** and **M**) Representative flow cytometry analysis (L) and quantification (M) of $tdTomato^+$ cells in corpus epithelium gated on $tdTomato^+EpCAM^+$ $DAPI^-$ cells from *Tff2-CreERT2; R26-tdTomato* mice at 24 hours after HDT or low-dose tamoxifen treatment (Ctrl, 75 mg/kg). SCC, side scatter. Data are shown as mean ± SD; n = 7 mice per group. Statistical significance is assessed with unpaired Student's t test. **P < 0.01. (**N**) Location and relative size of the $tdTomato^+$ compartment to the gland height over time as in (Fig. 2C). Data are shown as mean ± SD; n = 3 mice per time point per group.

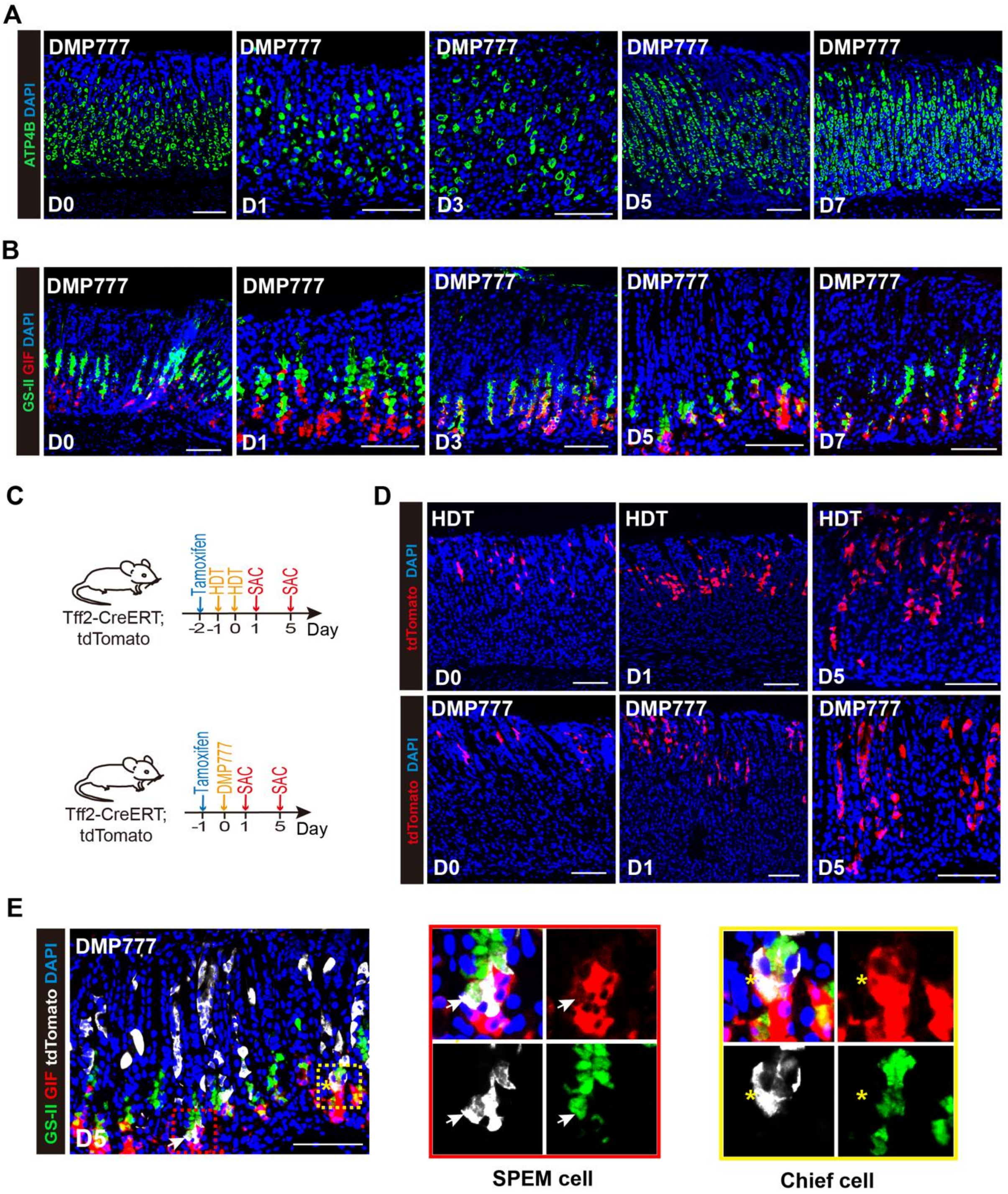


**Figure S3.** Acute corpus gland injury and repair processes induced by DMP777.

(**A**) Immunofluorescence staining for ATP4B (green) and DAPI (blue) in the corpus glands of WT mice before (D0) and after DMP777- induced acute injury (500 mg/kg, one dose) at different time points (D1, D3, D5, and D7). Scale bars, 100 μm. (**B**) Immunofluorescence staining for GS-II (green), GIF (red), and DAPI (blue) in the corresponding groups described in (A). Scale bars, 100 μm. (**C**) Schematic of the experimental design for comparing the effects of HDT-(top) and DMP777-(bottom) induced acute injury on lineage tracing of Tff2$^{+}$ progenitors. In brief, *Tff2-CreERT; R26-tdTomato* mice were treated with tamoxifen (75 mg/kg) to induce Cre recombinase 1 day before acute injury induction. Subsequently, mice were treated with HDT (300 mg/kg, once daily for 2 consecutive days) or DMP777(500 mg/kg, one dose), and corpus tissues were sampled for histological analysis at different time points after induction. SAC, scarify. (**D**) Immunofluorescence staining for tdTomato (red) and DAPI (blue) in the corpus glands of *Tff2-CreERT2; R26-tdTomato* mice before (D0) and after HDT (top) or DMP777 (bottom) treatment at different time points (D1 and D5). Scale bars, 100 μm. (**E**) (left) Immunofluorescence staining for GS-II (green), GIF (red), tdTomato (white), and DAPI (blue) in the corpus glands of *Tff2-CreERT2; R26-tdTomato* mice at 5 days (D5) after DMP777 treatment. White arrow indicates tdTomato$^{+}$ GS-II$^{+}$ GIF$^{+}$ SPEM cells. Yellow asterisk indicates tdTomato$^{+}$ GS-II$^{-}$ GIF$^{+}$ mature chief cells. Scale bars, 100 μm. (middle) High magnification details of the red dashed box in (left) to show SPEM cells. (right) High magnification details of the yellow dashed box in (left) to show chief cells.

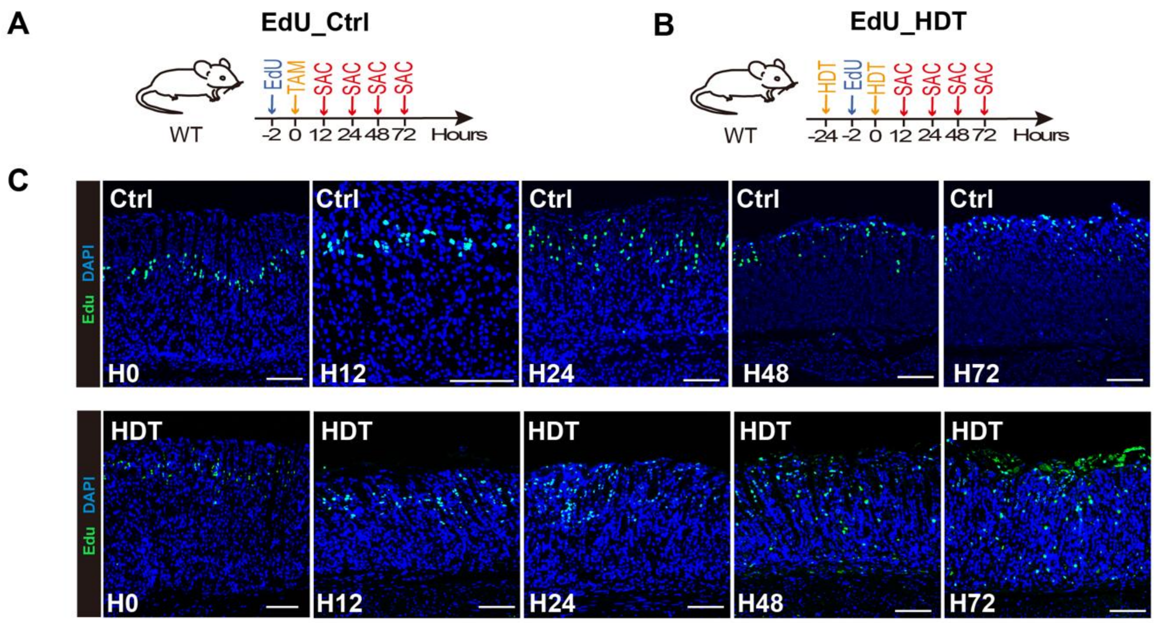


**Figure S4. Effect of H. pylori infection/Kras mutation on the fate of Tff2$^{+}$ progenitors**

(**A** and **B**) Schematic of the experimental design for 5-Ethynyl-2'-deoxyuridine (EdU) labeling proliferating cells migration as shown in (C, and Fig. 2F). In brief, WT mice were treated with low-dose tamoxifen (Ctrl) (A) or HDT (B). All groups were treated with EdU (50 mg/kg) to label proliferating cells 2 hours before the last dose of tamoxifen (H0). Corpus tissues were then sampled for histological analysis at different time points after tamoxifen induction (H12, H24, H48 and H72). SAC, scarify. (**C**) Immunofluorescence staining for EdU (green) and DAPI (blue) in the corpus glands of WT mice before (H0), and after HDT or Ctrl treatment at different time points (H0, H12, H24, H48 and H72). Scale bars, 100 μm.

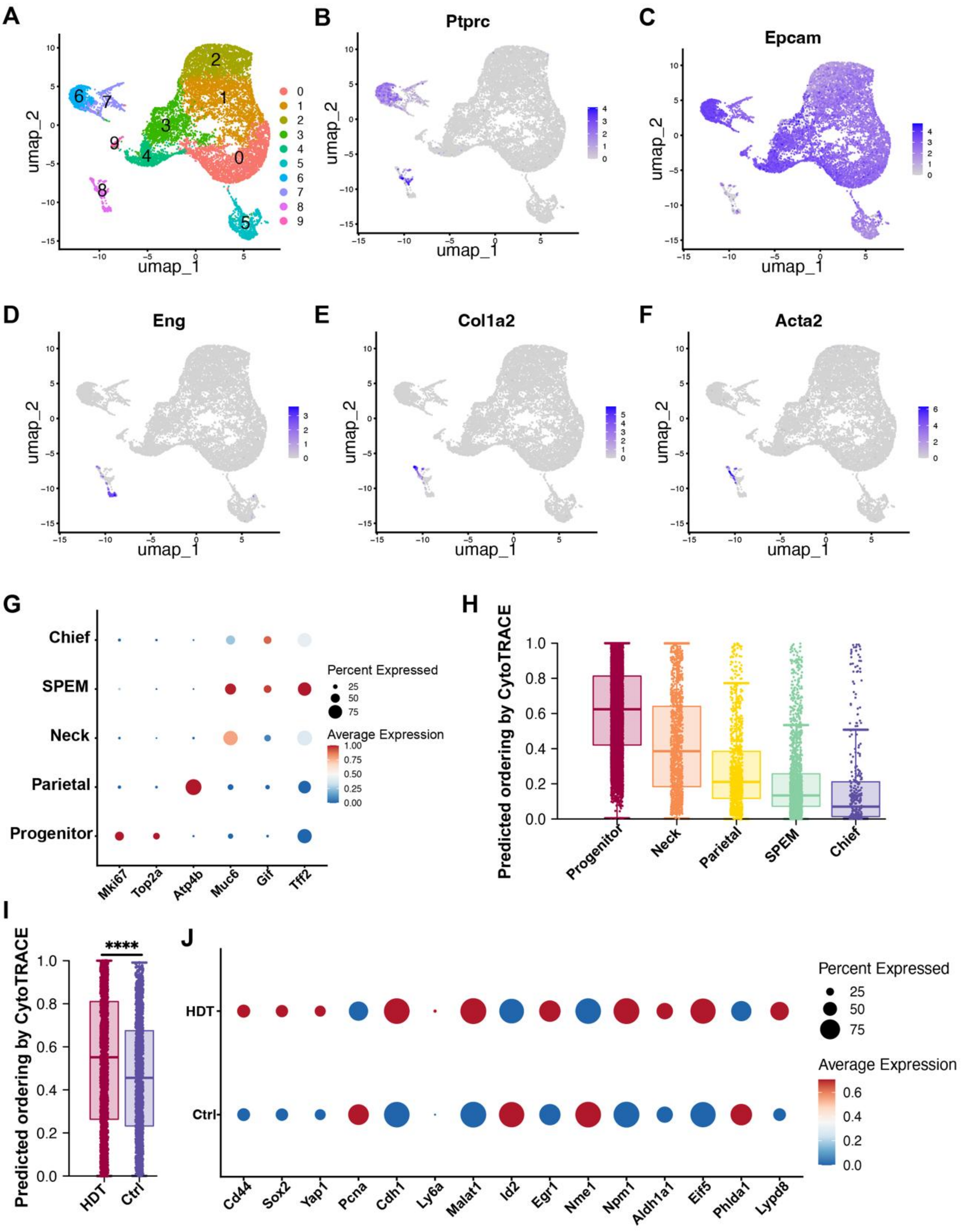


**Figure S5. Single-cell RNA-seq analysis of Tff2⁺ mouse corpus cells under homeostatic conditions or after acute injury**

(**A**) UMAP plots showing all clusters before removing low-quality and mixed cell populations in response to acute injury compared to control mice. Combining (B-F), we finally chose cluster 0-5 and cluster 9 as candidates for epithelial cell clusters. n=6. (**B-F**) UMAP plots showing the expression levels of canonical marker genes for different cell types. The color key from gray to purplish blue indicates low to high expression levels, respectively. (**G**) Dot plot showing the marker genes of 5 different cell types. The color key from blue to red indicates low to high expression levels, respectively. The circle size indicates the percentage of cells expressing a certain gene. (**H**) Comparison of CytoTRACE scores between epithelial clusters indicates progenitor clusters have the highest cell differentiation potential, while chief cells have the lowest. Boxes indicate the median ± interquartile range. (**I**) Comparison of CytoTRACE scores between progenitor clusters in response to HDT compared to Ctrl mice. Boxes indicate the median ± interquartile range. Statistical significance was assessed by two-sided Wilcoxon rank-sum test with Benjamini-Hochberg correction. ****$P < 0.0001$. (**J**) Dot plot showing the different expression of proliferation and stem cell-associated genes in progenitor cluster in response to HDT compared to Ctrl mice. The color key from blue to red indicates low to high expression levels, respectively. The circle size indicates the percentage of cells expressing a certain gene.

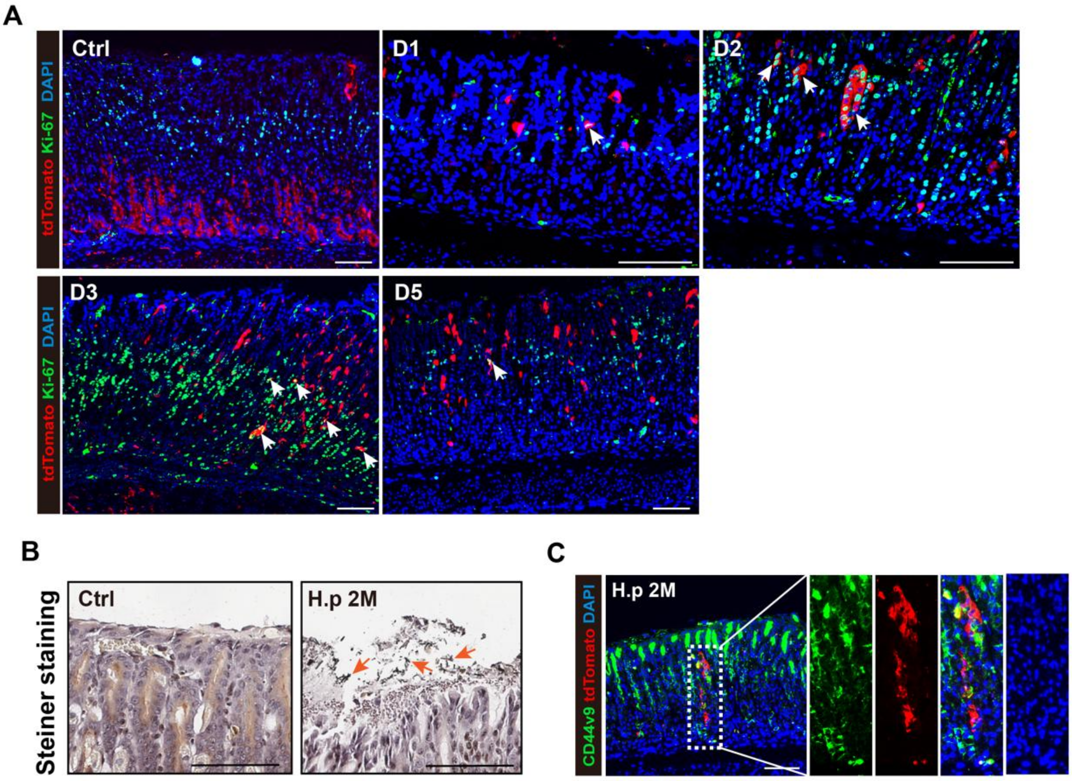


**Figure S6. Gif- labelled chief cell lineage tracing after acute injury**

(**A**) Immunofluorescence staining for Ki-67 (green), tdTomato (red), and DAPI (blue) in the corresponding *Gif-rtTA-eGFP; tetO-Cre; R26-tdTomato* (Gif-rtTA) mice 1d, 2d, 3d and 5d or control (Ctrl) after treatment as described in (Fig 3D). Scale bars, 100 μm. (**B**) Steiner staining for H. pylori (black to brown) in the corpus glands of *Tff2-CreETR; R26-tdTomato* mice 2 months (2M) after H. pylori infection (H.p) or control (Ctrl) as described in (Fig. 4A). Cell nuclei were counterstained with hematoxylin (purplish blue). Orange arrow indicates H. pylori in the superficial mucus of corpus glands. Scale bars, 100 μm. (**C**) Immunofluorescence staining for CD44v9 (green), tdTomato (red), and DAPI (blue) in the corresponding H. p mice 2 months (2M) after infection as described in (D). Scale bars, 100 μm.

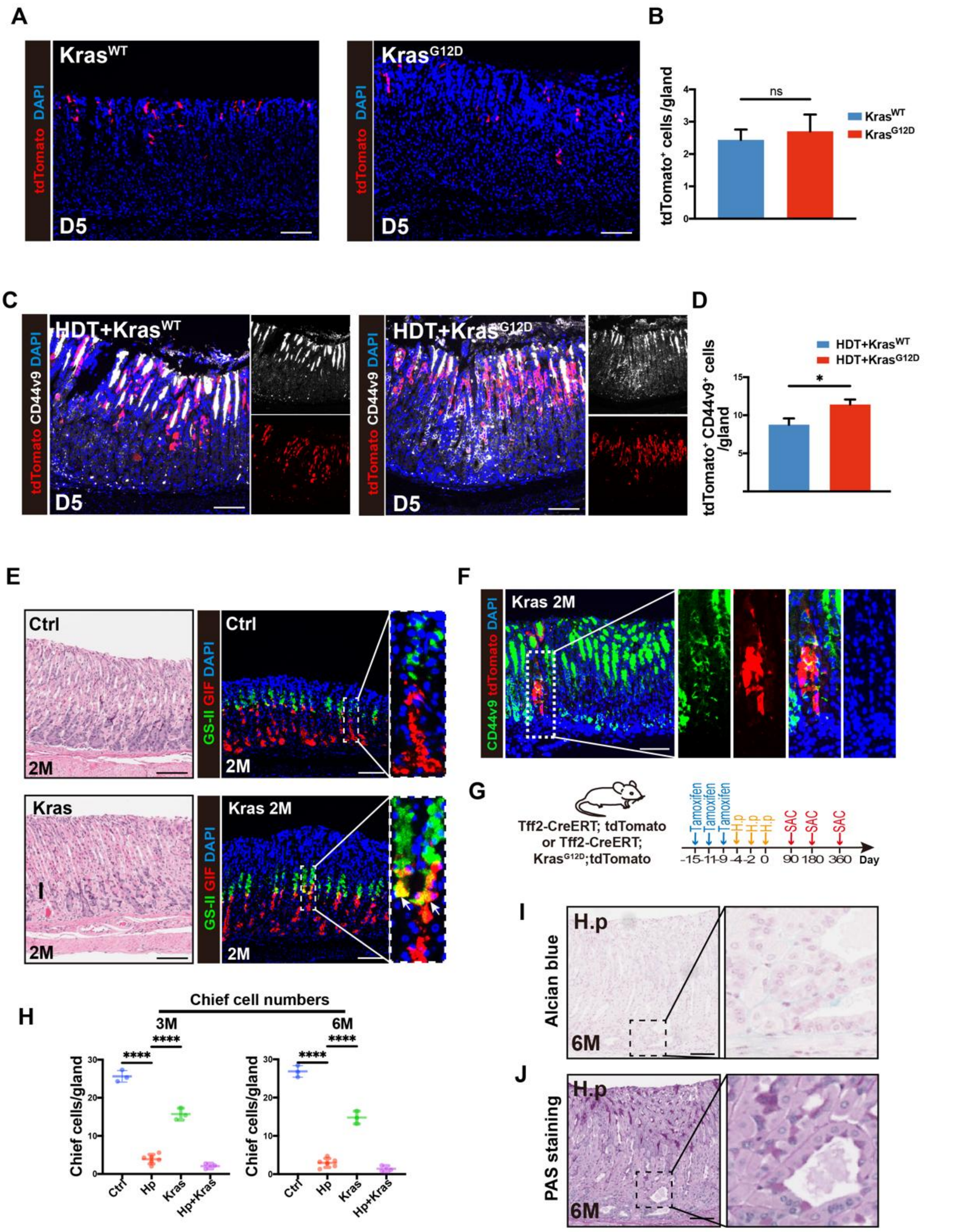


**Figure S7. Kras mutation in Tff2+ progenitor led to SPEM**

(**A**) Immunofluorescence staining for tdTomato (red) and DAPI (blue) in the corpus glands of *Tff2-CreERT; Kras$^{LSL\text{-}G12D/+}$; R26-tdTomato* (Kras$^{G12D}$) mice and *Tff2-CreERT; Kras $^{+/+}$;R26-tdTomato* (Kras$^{WT}$) mice 5 days after low dose tamoxifen (75 mg/kg). Scale bars, 100 µm. (**B**) Quantification of the number of tdTomato$^{+}$ cells per gland as in (A). Data are shown as mean ± SD; n = 3 mice per group. Statistical significance was assessed by unpaired Student's t test. ns, not significant. (**C**)

Immunofluorescence staining for tdTomato (red), CD44v9 (white), and DAPI (blue) in the corpus glands of $Kras^{G12D}$ mice and $Kras^{WT}$ mice 5 days after HDT treatment. Scale bars, 100 µm. (**D**) Quantification of the number of $tdTomato^{+}$ $CD44v9^{+}$ cells per gland as in (C). Data are shown as mean ± SD; n = 3 mice per group. Statistical significance was assessed by unpaired Student's t test. *P < 0.05. (**E**) (left) H&E staining and (right) immunofluorescence staining for GS-II (green), GIF (red), and DAPI (blue) in the corpus glands of *Tff2-CreETR; $Kras^{LSL\text{-}G12D/+}$; R26-tdTomato* (Kras) mice or *Tff2-CreETR; $Kras^{+/+}$;R26-tdTomato* (Ctrl) mice 2 months (2M) after tamoxifen induction. Scale bars, 100 µm. (**F**) Immunofluorescence staining for CD44v9 (green), tdTomato (red), and DAPI (blue) in the corresponding Kras mice 2 months (2M) after tamoxifen induction as described in (E). Scale bars, 100 µm. (**G**) Schematic of the experimental design for the long-term effect of H. pylori infection and (or) Kras mutation on the fate of $Tff2^{+}$ progenitors. In brief, *Tff2-CreERT; $Kras^{LSL\text{-}G12D/+}$; R26-tdTomato* mice or *Tff2-CreERT; $Kras^{+/+}$;R26-tdTomato* mice were treated with 3 doses of tamoxifen (75 mg/kg) to induce Kras mutation and/or nuclear transfer of Cre recombinase, then the mice were orally gavaged with H. pylori (SS1, $2*10^{9}$ CFU, every 2 days*3 doses) or vehicle 1 day later. Corpus tissues were then sampled for histological analysis at different time points after oral gavage (3 M, 6M and 12M). (**H**) Quantification of the number of chief cells per gland in corpus tissues of *Tff2-CreERT; $Kras^{+/+}$;R26-tdTomato* mice 3mo and 6mo after H.p infection (H.p) or vehicle (Ctrl), or *Tff2-CreERT; $Kras^{LSL\text{-}G12D/+}$; R26-tdTomato* mice 3mo and 6mo after Kras mutation (Kras) or Kras mutation plus H.p infection (Kras+H.p). Schematic of the experimental design as shown in (G). Data are shown as mean ± SD. n = 3 to 12 mice per group. Statistical significance was assessed with one-way ANOVA. ns, not significant, *P < 0.05, **P < 0.01, ***P < 0.001, ****P < 0.0001. (**I**) Alcian blue staining for acidic mucins (blue) in the corpus glands of *Tff2-CreERT; R26-tdTomato* mice 6 months after H.p infection (H.p). Cell nuclei were counterstained with hematoxylin (purplish blue). Scale bars, 100 µm. (**J**) Periodic acid Schiff (PAS) staining neutral mucins (magenta) in the corresponding groups described in (I). Cell nuclei were counterstained with hematoxylin (purplish blue). Scale bars, 100 µm.

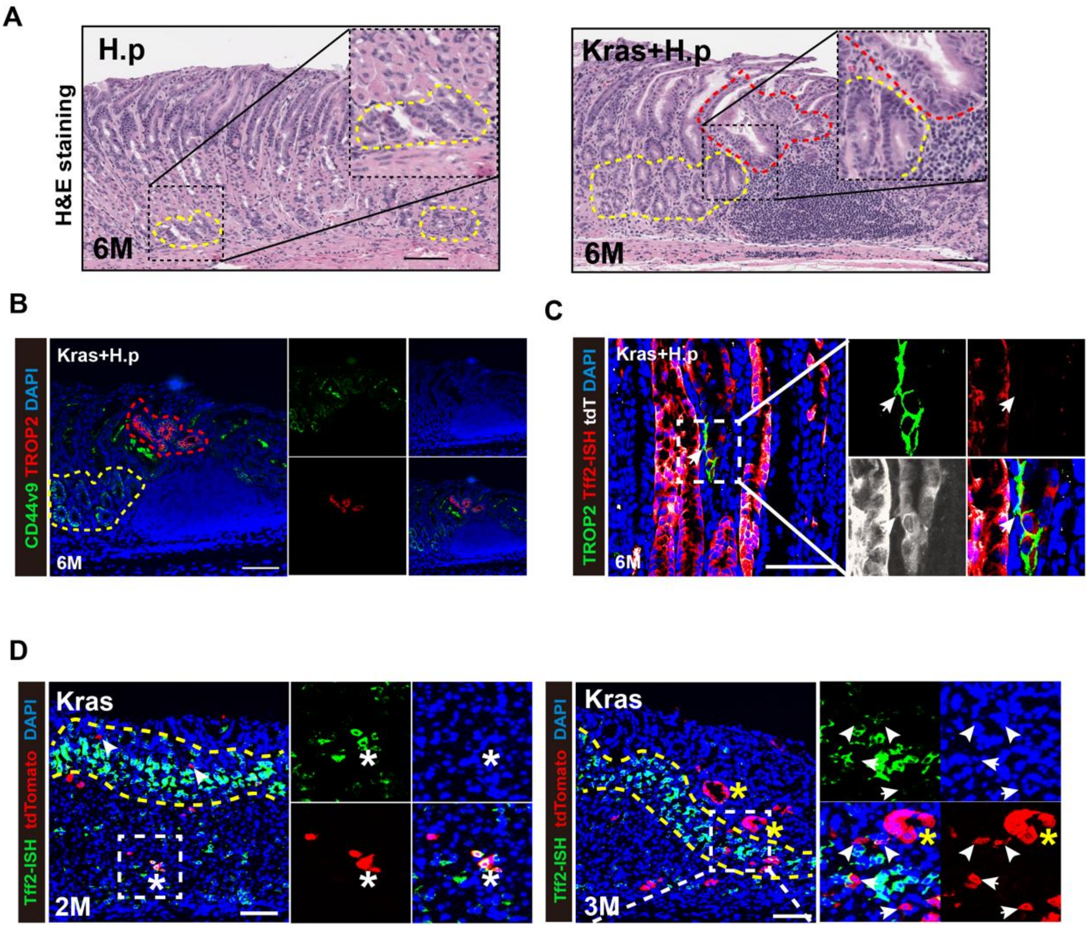


**Figure S8. Tff2$^{+}$ progenitors progress to metaplasia and dysplasia**

(**A**) H&E staining in the corpus glands of *Tff2-CreERT; Kras $^{+/+}$;R26-tdTomato* mice 6 months after H.pylori infection (H.p) (left), or *Tff2-CreETR; Kras$^{LSL\text{-}G12D/+}$; R26-tdTomato* mice 6 months after tamoxifen-induced Kras mutation plus H.p infection (Kras+ H.p) (right). Yellow dashed box indicates metaplasia, red dashed box indicates dysplasia. Scale bars, 100 µm. (**B**) Immunofluorescence staining for CD44v9 (green), TROP2 (red), and DAPI (blue) in the corpus glands of *Tff2-CreERT; Kras $^{+/+}$;R26-tdTomato* mice 6 months after H.p, or *Tff2-CreETR; Kras$^{LSL\text{-}G12D/+}$; R26-tdTomato* mice 6 months after Kras+ H.p. Yellow dashed box indicates metaplasia, red dashed box indicates dysplasia. Scale bars, 100 µm. (**C**) In situ hybridization of *Tff2* (red) and immunofluorescence staining for TROP2(green), tdTomato (white), and DAPI (blue) in the corpus glands of *Tff2-CreETR; Kras$^{LSL\text{-}G12D/+}$; R26-tdTomato* mice 6 months after Kras+ H.p. white asterisk indicates TROP2$^{+}$ dysplastic cells lacking *Tff2* mRNA expression were derived from Tff2$^{+}$ progenitors. Scale bars, 100 μm. (**D**) In situ hybridization of *Tff2* (green) and immunofluorescence staining for tdTomato (red), and DAPI (blue) in the corpus glands of *Tff2-CreETR; Kras$^{LSL\text{-}G12D/+}$; R26-tdTomato* mice 2 (left) and 3 (right) months after Kras mutation. Yellow dashed box indicates Tff2-mRMA$^{+}$ isthmus region, white arrow indicates isthmus-resident tdTomato$^{+}$ cells, white asterisk indicates tdTomato$^{+}$ Tff2-mRMA$^{+}$ SPEM cells, yellow asterisk indicates tdTomato$^{+}$ dysplasia. Scale bars, 100 μm.

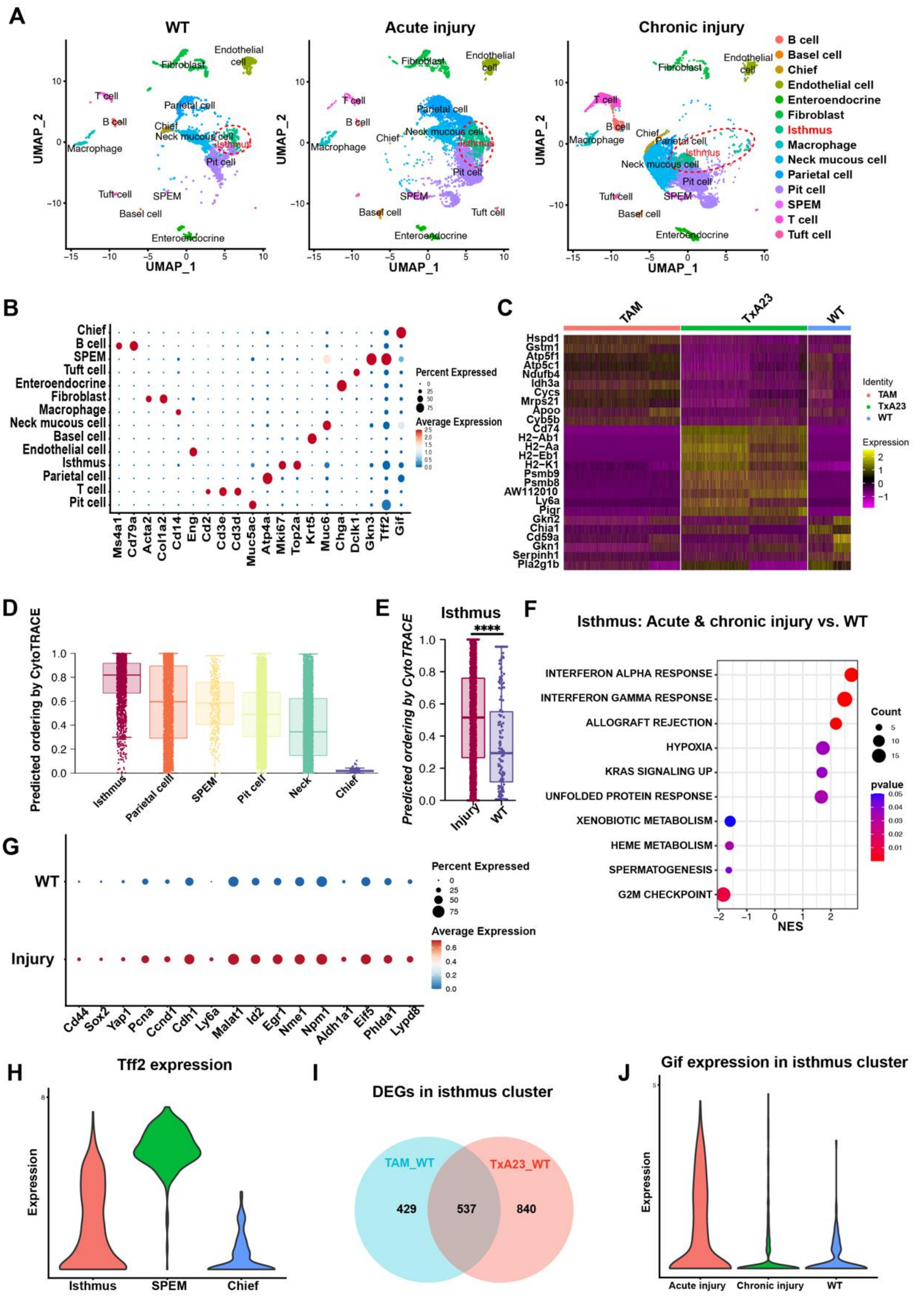


**Figure S9. Single-cell RNA-seq analysis of mouse corpus cells from a published dataset**

(**A**) Single-cell RNA-seq of corpus tissues from WT (left), acute-injury (middle) and chronic-injury (right) mice with a published data set[35]. Clusters identified based on cell type-specific markers revealed transcriptomic changes after acute- or chronic-injury mainly in epithelial cells. Specifically in the isthmus clusters (red dashed box), which differ in a transcriptional state in response to acute- or chronic-injury compared to WT mice. WT, nontreated BALB/c mice, n=2. Acute injury, high dose tamoxifen induced acute gastric injury in BALB/c mice, n=2. Chronic injury, autoimmune gastric metaplasia in the TxA23 mice, n=3. (**B**) Dot plot showing the marker genes of 14 different cell types. The color key from blue to red indicates low to high expression levels, respectively. The circle size indicates the percentage of cells expressing a certain gene. (**C**) Heatmap showing differentially expressed genes (DEGs) of corpus cells from WT, acute injury (TAM) and chronic injury (TxA23) mice. The color key from purple to yellow indicates low to high expression levels, respectively. (**D**) Comparison of CytoTRACE scores between epithelial clusters indicates isthmus clusters have the highest cell differentiation potential, while chief cells have the lowest. Boxes indicate the median ± interquartile range. (**E**) Comparison of CytoTRACE scores between epithelial clusters. Boxes indicate the median ± interquartile range. Statistical significance was assessed by two-sided Wilcoxon rank-sum test with Benjamini-Hochberg correction. ****P < 0.0001. (**F**) Dot plot for Gene Set Enrichment Analysis (GSEA) comparing the expression profile of the isthmus cluster from acute- and chronic-injury mice and WT mice. NES, normalized enrichment score. Statistical significance was assessed by a global FDR adjusted for multiple testing according to the method of Benjamini and Hochberg. (**G**) Dot plot showing the different expression of proliferation and stem cell-associated genes in isthmus cluster from injury or WT mice. The color key from blue to red indicates low to high expression levels, respectively. The circle size indicates the percentage of cells expressing a certain gene. (**H**) Expression of *Tff2* in different clusters (isthmus, SPEM and chief cell). (**I**) Venn diagram showing the overlap between DEGs of isthmus clusters in the HDT-induced acute injury group relative to the WT group (TAM_WT) and DEGs of isthmus clusters in the TxA23-induced chronic injury group relative to the WT group (TxA23_WT). (**J**) Expression of *Gif* in isthmus cluster across different groups (acute injury, chronic injury and WT).

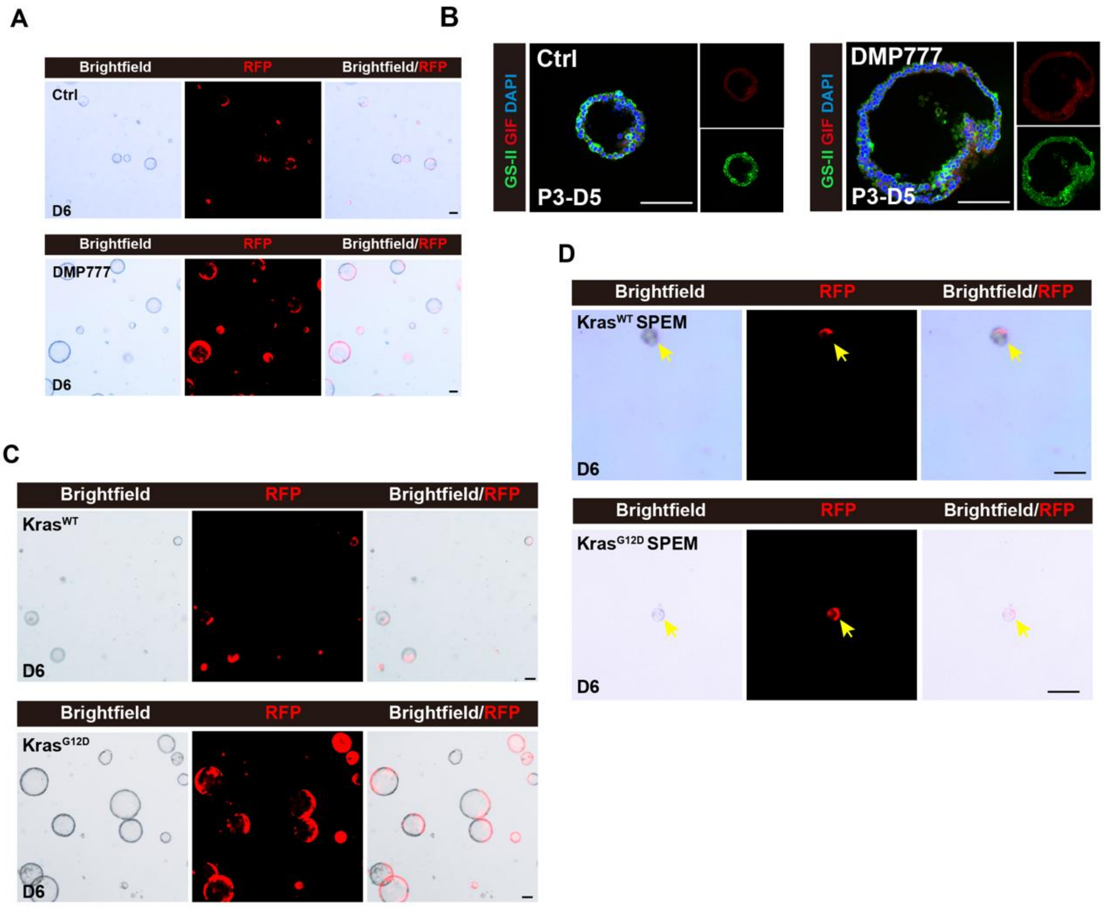


**Figure S10. Organoid cultures for Tff2$^+$ progenitor and SPEM cells**

(**A**) Bright field and fluorescence images of primary organoid cultures (P0) at 6 days (D6) for Tff2$^+$ progenitor in Ctrl or DMP777 groups as described in (Fig. 6A). Scale bars, 100 μm. (**B**) Immunofluorescence staining for GS-II (green), GIF (red) and DAPI (blue) 5 days after the third passage (P3-D5) of organoids for Tff2$^+$ progenitors as described in (Fig. 6A). Scale bars, 100 μm. (**C** and **D**) Bright field and fluorescence images of primary organoid cultures (P0) at 6 days (D6) for Tff2$^+$ progenitor (Kras$^{G12D}$ or Kras$^{WT}$) (C) and Tff2$^+$ SPEM (Kras$^{G12D}$ or Kras$^{WT}$) (D) as described in (Fig. 6G, 6H). Yellow arrow indicates tdTomato$^+$ SPEM organoid. Scale bars, 100 μm.

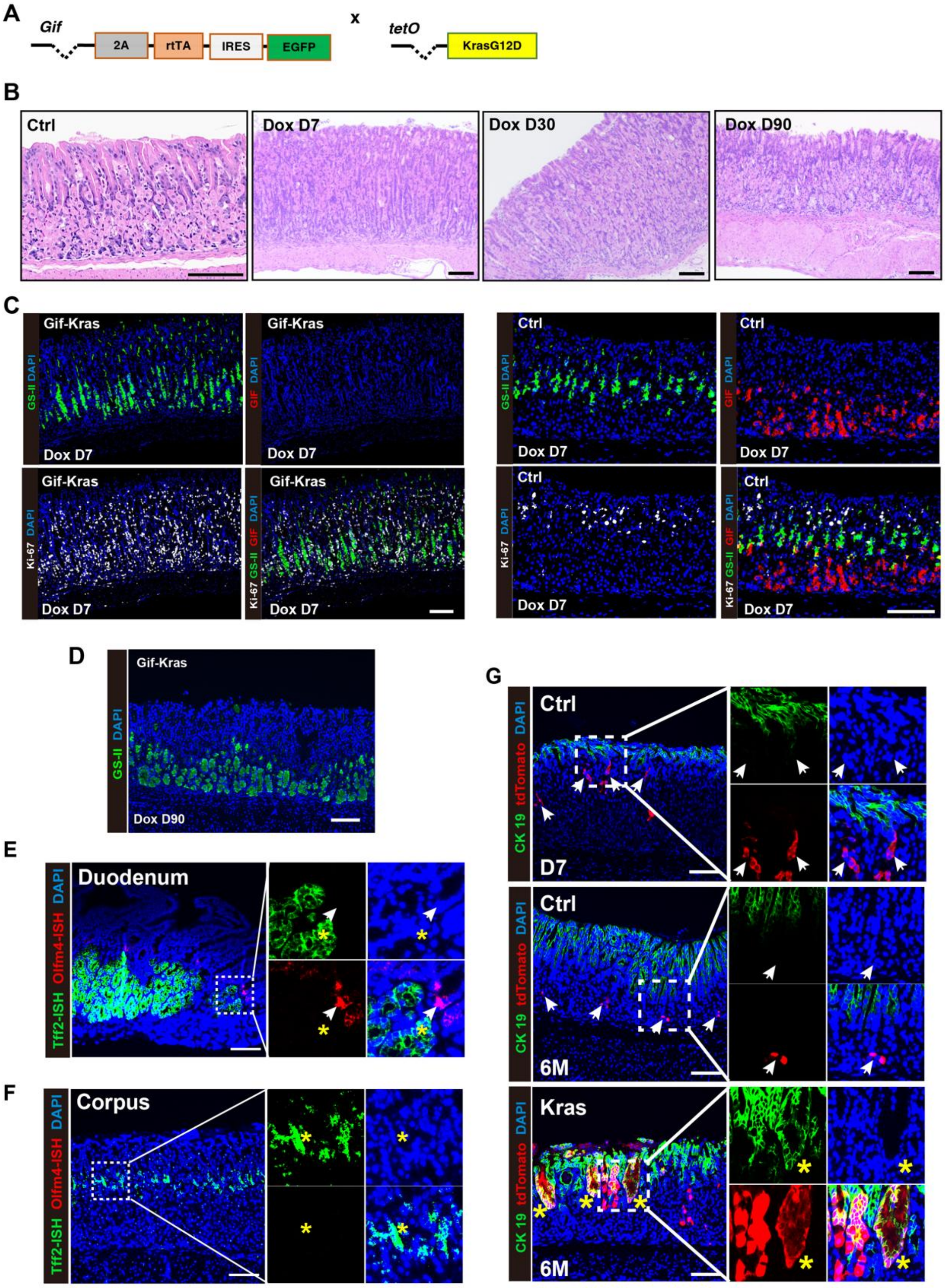


**Figure S11. Kras mutation in Gif⁺ chief cell led to loss**

(**A**) Schematic representing the experimental design for (B-C). In brief, *Gif-rtTA-eGFP; tetO-Kras$^{G12D}$* (Gif-Kras) mice were treated with doxycycline throughout the process, and corpus tissues were sampled for histological analysis at different time points (D7, D30, D90) after induction. *tetO-Kras$^{G12D}$* mice were used as controls (Ctrl).

(**B**) H&E staining 7, 30, 90 days (D7, D30, D90) after doxycycline treatment in the corpus glands of Gif-Kras mice and Ctrl mice as described in (A). Scale bars, 100 µm. (**C**) Immunofluorescence staining for GS-II (green), GIF (red), Ki-67 (white) and DAPI (blue) 7 days after doxycycline treatment in corpus glands of Gif-Kras mice and Ctrl mice as described in (A). Scale bars, 100 µm. (**D**) Immunofluorescence staining for GS-II (green) and DAPI (blue) 90 days after doxycycline treatment in corpus glands of Gif-Kras mice as described in (A). Scale bars, 100 µm. (**E** and **F**) In situ hybridization of *Tff2* (green), *Olfm* (red), and DAPI (blue) in the duodenal Brunner glands (E) and in the corpus glands (F) of nontreated WT mice. White arrow indicates Olfm-mRMA$^{+}$ intestinal stem cells. Yellow asterisk indicates Tff2-mRMA$^{+}$ cells in Brunner glands (E) or corpus isthmus region. Scale bars, 100 µm. (**G**) Immunofluorescence staining for CK19 (green), tdTomato(red), and DAPI (blue) in the corpus glands of *Tff2-CreETR; Kras* $^{+/+}$*;R26-tdTomato* mice 7days (D7) or 6 months (6M) after tamoxifen treatment (Ctrl) or *Tff2-CreETR; Kras*$^{LSL\text{-}G12D/+}$*; R26-tdTomato* mice 6 months after Kras mutation (Kras). White arrow indicates tdTomato$^{+}$ CK19$^{-}$ cells, yellow asterisk indicates tdTomato$^{+}$ CK19$^{+}$ pit cells derived from Tff2$^{+}$ progenitors. Scale bars, 100 µm.

A

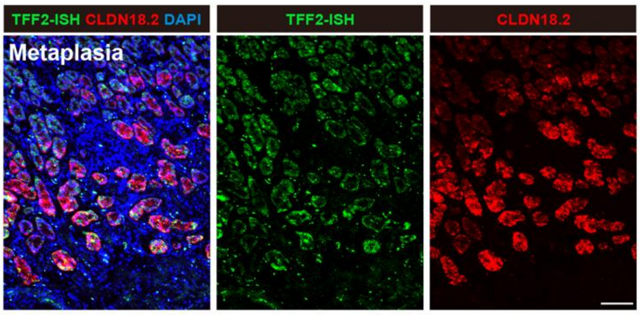


B

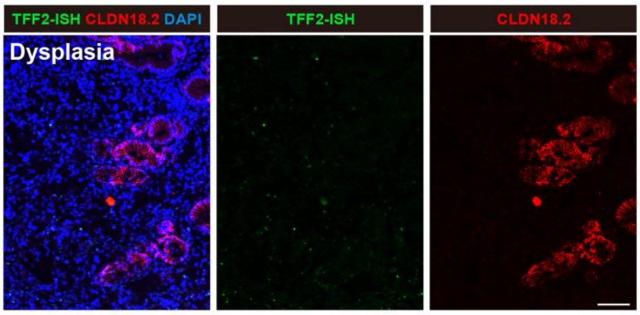


C

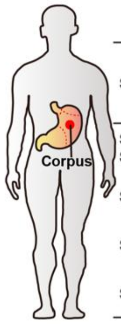


| Sample | Age | Sex | Histopathological diagnosis | Site of origin | Lauren's classificatio n | Pathologic al Staging | Sampling site |
|---|---|---|---|---|---|---|---|
| Sample01, Sample02 | 62 | F | Poorly differentiated adenocarcinoma | Corpus | Diffuse | pT4aN1M0 | Tumor, Adjacent mucosa |
| Sample03 | 65 | M | Moderately differentiated adenocarcinoma | Corpus | Intestinal | pT2N0M0 | Tumor |
| Sample04 | 69 | M | Moderately differentiated adenocarcinoma | Corpus | Intestinal | pT2N1M0 | Tumor |
| Sample05 | 37 | F | Poorly differentiated adenocarcinoma | Corpus | Intestinal | pT4N1M0 | Adjacent mucosa |

D

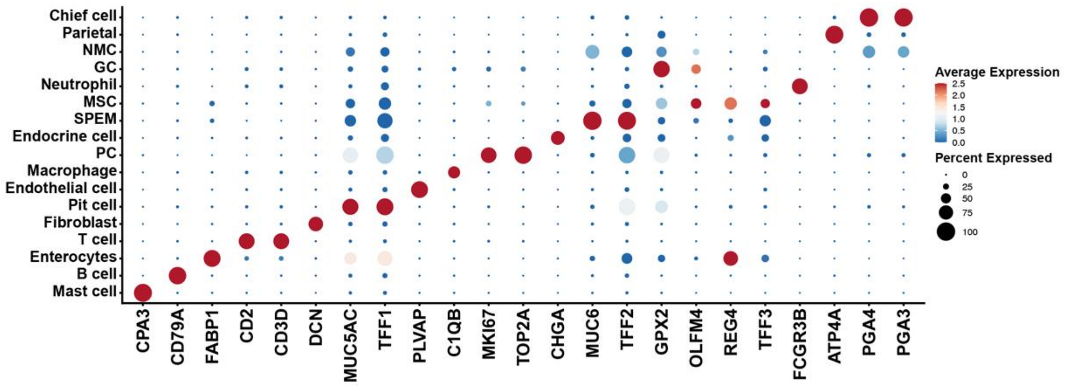


**Figure S12. In situ hybridization of TFF2 in human tissues**

(**A** and **B**) In situ hybridization of *TFF2* (green) and immunofluorescence staining for CLDN18.2 (red), and DAPI (blue) in the human mucinous metaplasia (A) and dysplasia (B) lesions. Scale bars, 100 μm. (**C**) Clinicopathologic characteristics of five corpus samples used for single-cell RNA-seq analysis as in Fig. 6A. (**D**) Dot plot showing the marker genes of 17 different cell types. The color key from blue to red indicates low to high expression levels, respectively. The circle size indicates the percentage of cells expressing a certain gene.

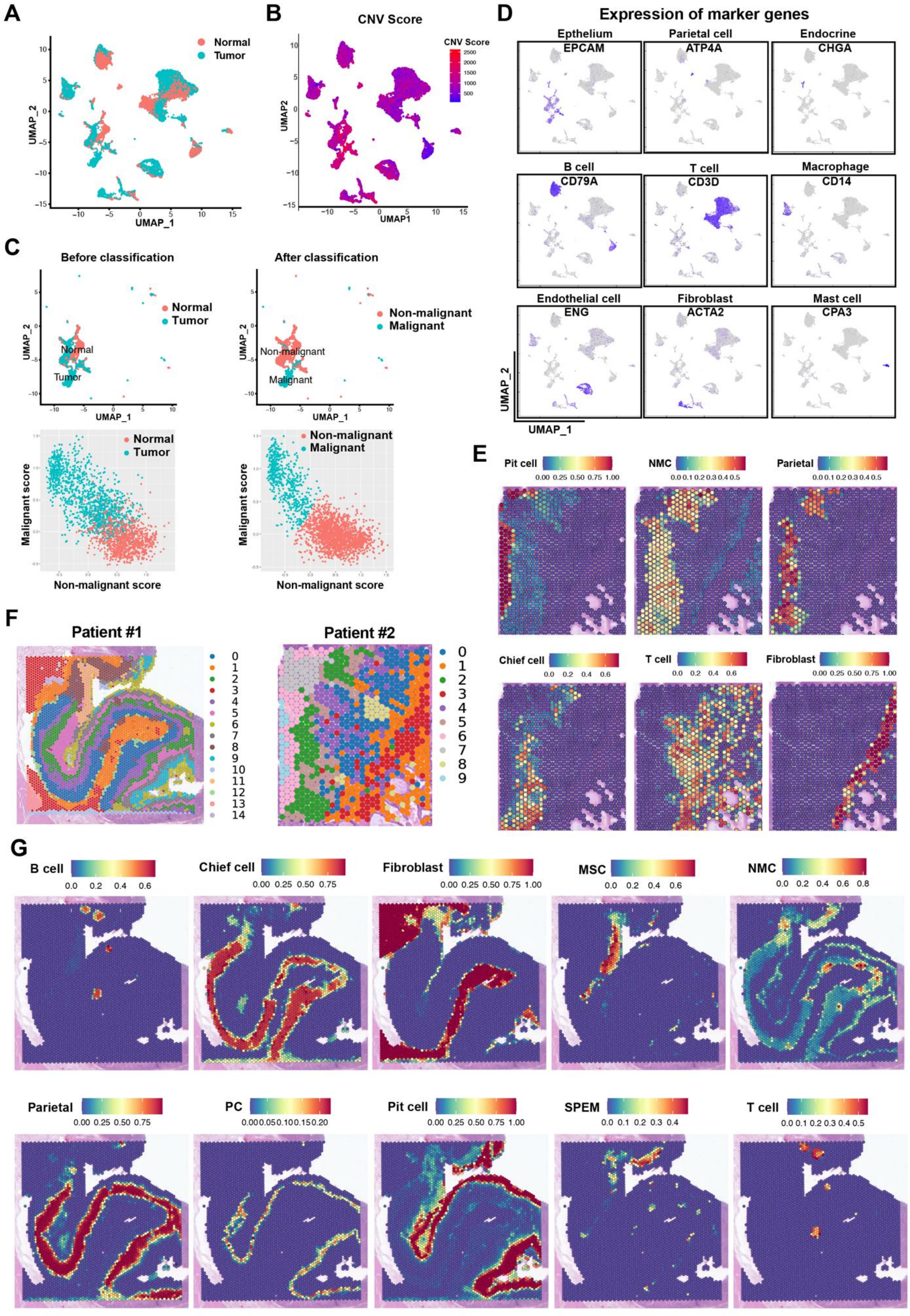


**Figure S13. scRNA-seq and spatial transcriptomics analysis in human samples**

(**A**) UMAP plot exhibiting the sample origin in human gastric corpus. (**B**) UMAP plot of colored by CNV score of all cells. (**C**) (Left top) UMAP plot exhibiting the sample origin of epithelial clusters (without endocrine or parietal cells) (Before classification). (Left bottom) Scatter plot exhibiting the initial malignant and non-malignant score for each epithelial cell based on differential expression genes according to sample origin from tumor or normal tissue samples. (Right top) UMAP plot of the classification of malignant and non-malignant cells using the k-means clustering algorithm. We repeated this process 7 times until the classification results were stabilized (After classification). (Right bottom) Scatter plot exhibiting the malignant and non-malignant final score for each epithelial cell after 7th classification of malignant and non-malignant epithelial cells. (**D**) UMAP plots showing the expression levels of canonical marker genes for nine cell types. The color key from gray to purplish blue indicates low to high expression levels, respectively. (**E**) Integration result by transferring the scRNA-seq dataset to the spatial transcriptomics dataset (patient #2). Each spot represents 2-10 cells on the corresponding HE section. Color-coded values (from blue to red) reflect low to high predicted values. NMC, mucous neck cell. (**F**) Louvain clustering of spatial transcriptomics in patient #1 (left) and patient #2 (right). (**G**) Integration result by transferring the scRNA-seq dataset to the spatial transcriptomics dataset (patient #1). Each spot represents 2-10 cells on the corresponding HE section. Color-coded values (from blue to red) reflect low to high predicted values. MSC, metaplastic stem-like cells; NMC, mucous neck cell; PC, normal proliferative cell cluster.

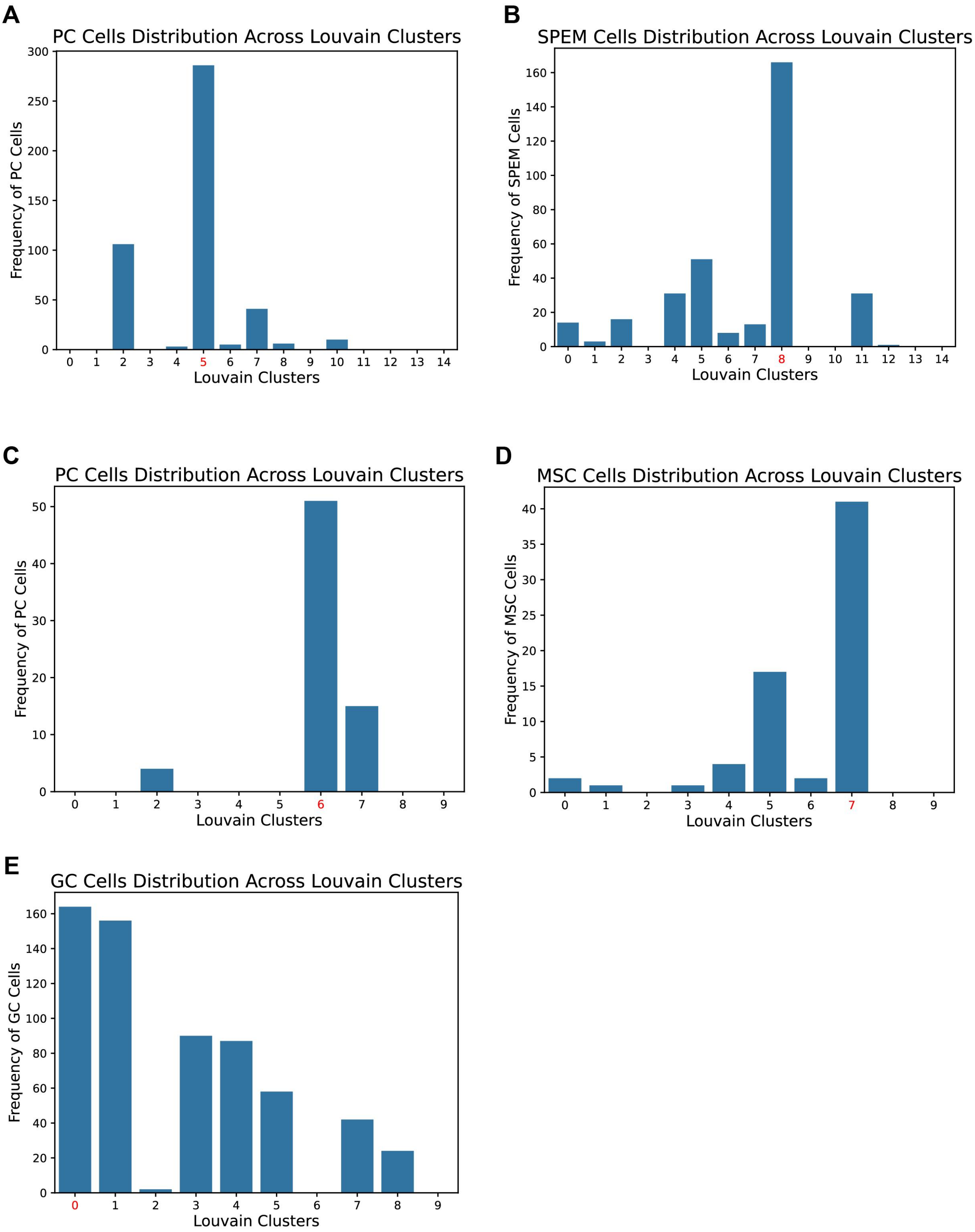


**Figure S14. Cell distribution across Louvain clusters**

(**A** and **B**) PC (A) and SPEM (B) cell distribution across Louvain clusters in patient #1.

(**C**, **D** and **E**) PC (A), MSC (B) and GC (C) cell distribution across Louvain clusters in patient #2.
