## Supplemental Table1 for "*Tff2* marks gastric corpus progenitors that give rise to pyloric metaplasia/SPEM following injury"

Primers used in this study

| Gene(Mus musculus) | Forward sequence | Reverse sequence |
|---|---|---|
| Gif | AAGCACAGCGCAAAAACTCC | GCAACCCCTTCATCCAAAGG |
| Lgr5 | CCTACTCGAAGACTTACCCAGT | GCATTGGGGTGAATGATAGCA |
| Pgc | ATGAAGAGTATCCGGGAGACC | TGGGCTCATAGAGTACACTGTAG |
| Muc6 | CAGCTCAACAAGGTGTGTGC | GGTCTCCTCGTAGTTGCAGG |
| Fabp5 | TGAAAGAGCTAGGAGTAGGACTG | CTCTCGGTTTTGACCGTGATG |
| Nme1 | AGGAGCACTACACTGACCTGA | GGTTGGTCTCTCCAAGCATCA |
| Myc | ATGCCCCTCAACGTGAACTTC | CGCAACATAGGATGGAGAGCA |
| Lgr4 | CCCGACTTCGCATTCACCAA | GCCTGAGGAAATTCATCCAAGTT |
| Wfdc2 | TGCCTGCCTGTCGCCTCTG | TGTCCGCACAGTCCTTGTCCA |
| Tff2 | TGCTCTGGTAGAGGGCGAG | CGACGCTAGAGTCAAAGCAG |
| Sox9 | GTGCAAGCTGGCAAAGTTGA | TGCTCAGTTCACCGATGTCC |
| Yap1 | ACCCTCGTTTTGCCATGAAC | TGTGCTGGGATTGATATTCCGTA |
| Cd44 | TCGATTTGAATGTAACCTGCCG | TCGATTTGAATGTAACCTGCCG |
| Gkn3 | CCGTTGCATTCGCTGGAGA | AACTGTCGCTAGTGTTCGTCA |
| Cxcr4 | GCTGGCTGAAAAGGCAGTCTAT | TGACGTCGGCAAAGATGAAGT |
| Prom1 | CTCCCATCAGTGGATAGAGAACT | ATACCCCCTTTTGACGAGGCT |
| Sox2 | GCGGAGTGGAAACTTTTGTCC | CGGGAAGCGTGTACTTATCCTT |
| Atp4a | GATGGAGATTAACGACCACCAG | ACGGGCAAACTTCACATACTC |
| Chga | ATCCTCTCTATCCTGCGACAC | GGGCTCTGGTTCTCAAACACT |
| Muc5ac | CAGGACTCTCTGAAATCGTACCA | AAGGCTCGTACCACAGGGA |
| Gapdh | AGGTCGGTGTGAACGGATTTG | TGTAGACCATGTAGTTGAGGTCA |
